# An increasing Arctic-boreal CO_2_ sink offset by wildfires and source regions

**DOI:** 10.1101/2024.02.09.579581

**Authors:** Anna-Maria Virkkala, Brendan M. Rogers, Jennifer D. Watts, Kyle A. Arndt, Stefano Potter, Isabel Wargowsky, Edward A. G. Schuur, Craig See, Marguerite Mauritz, Julia Boike, Syndonia M. Bret-Harte, Eleanor J. Burke, Arden Burrell, Namyi Chae, Abhishek Chatterjee, Frederic Chevallier, Torben R. Christensen, Roisin Commane, Han Dolman, Bo Elberling, Craig A. Emmerton, Eugenie S. Euskirchen, Liang Feng, Mathias Goeckede, Achim Grelle, Manuel Helbig, David Holl, Järvi Järveoja, Hideki Kobayashi, Lars Kutzbach, Junjie Liu, Ingrid Liujkx, Efrén López-Blanco, Kyle Lunneberg, Ivan Mammarella, Maija E. Marushchak, Mikhail Mastepanov, Yojiro Matsuura, Trofim Maximov, Lutz Merbold, Gesa Meyer, Mats B. Nilsson, Yosuke Niwa, Walter Oechel, Sang-Jong Park, Frans-Jan W. Parmentier, Matthias Peichl, Wouter Peters, Roman Petrov, William Quinton, Christian Rödenbeck, Torsten Sachs, Christopher Schulze, Oliver Sonnentag, Vincent St.Louis, Eeva-Stiina Tuittila, Masahito Ueyama, Andrej Varlagin, Donatella Zona, Susan M. Natali

**Affiliations:** Woodwell Climate Research Center, Falmouth, USA; Center for Ecosystem Science and Society, Northern Arizona University Flagstaff, USA; Department of Biological Sciences, Northern Arizona University Flagstaff, USA; Biological Sciences, University of Texas at El Paso, El Paso, USA; Permafrost Research Section, Alfred Wegener Institute Helmholtz Center for Polar and Marine Research, Potsdam, Germany; Department of Geography, Humboldt University, Berlin, Germany; Institute of Arctic Biology, University of Alaska Fairbanks, Fairbanks, USA; Met Office Hadley Centre, Exeter, UK; Institute of Life Science and Natural Resources, Korea University, Seoul, South Korea; NASA Jet Propulsion Laboratory, California Institute of Technology, USA; Laboratoire des Sciences du Climat et de l’Environnement, LSCE/IPSL, CEA-CNRS-UVSQ, Université Paris-Saclay, Gif-sur-Yvette, France; Department of Ecoscience, Arctic Research Center, Aarhus University, Roskilde, Denmark; Water, energy and environmental engineering research unit, University of Oulu, Oulu, Finland; Department of Earth and Environmental Sciences, Columbia University, NY, USA; Royal Netherlands Institute for Sea Research, Texel, Netherlands; Vrije Universiteit, Amsterdam, the Netherlands; Department of Geosciences and Natural Resource Management, University of Copenhagen, Copenhagen, Denmark; Department of Biological Sciences, University of Alberta, Edmonton, Canada; NCEO, School of GeoSciences, University of Edinburgh, UK; Max Planck Institute for Biogeochemistry, Jena, Germany; Linnaeus University, Växjö, Sweden; Department of Physics and Atmospheric Science, Dalhousie University, Halifax, Canada; Institute of Soil Science, Center for Earth System Research and Sustainability (CEN), Universität Hamburg, Hamburg, Germany; Department of Forest Ecology and Management, Swedish University of Agricultural Sciences, Umeå, Sweden; Research Institute for Global Change, Japan Agency for Marine-Earth Science and Technology; Department Biology, San Diego State University, San Diego, USA; Institute for Atmospheric and Earth System Research / Physics, Faculty of Science, University of Helsinki, Finland; Department of Environmental and Biological Sciences, University of Eastern Finland, Finland; Forestry and Forest Products Research Institute, Tsukuba, Ibaraki, Japan; Institute for Biological Problems of Cryolithozone of the Siberian Branch of the RAS - Division of Federal Research Centre “The Yakut Scientific Centre of the Siberian Branch of the Russian Academy of Sciences”, Yakutsk, Russia; Integrative Agroecology, Agroecology and Environment, Agroscope, Zurich, Switzerland; Département de géographie, Université de Montréal, Montréal, Québec, Canada; Environment and Climate Change Canada, Climate Research Division, Victoria, Canada; Earth System Division, National Institute for Environmental Studies/Department of Climate and Geochemistry Research, Meteorological Research Institute, Japan; Division of Atmospheric Sciences, Korea Polar Research Institute, Incheon, Republic of Korea; Centre for Biogeochemistry in the Anthropocene, Department of Geosciences, University of Oslo, Oslo, Norway; Centre for Isotope Research, Energy and Sustainability Research Institute Groningen, Groningen University, The Netherlands; Cold Regions Research Centre, Wilfrid Laurier University, Waterloo, Canada; GFZ German Research Centre for Geosciences, Germany; Department of Renewable Resources, University of Alberta, Edmonton, Canada; School of Forest Sciences, University of Eastern Finland; Graduate School of Agriculture, Osaka Metropolitan University; A.N. Severtsov Institute of Ecology and Evolution, Russian Academy of Sciences, Moscow, Russia; Dept. of Meteorology and Air Quality, Wageningen University, The Netherlands; Institute of Geoecology, Technische Universität Braunschweig, Braunschweig, Germany; Department of Environment and Minerals, Greenland Institute of Natural Resources, Nuuk, Greenland

## Abstract

The Arctic-Boreal Zone (ABZ) is rapidly warming, impacting its large soil carbon stocks. We use a new compilation of terrestrial ecosystem CO_2_ fluxes, geospatial datasets and random forest models to show that although the ABZ was an increasing terrestrial CO_2_ sink from 2001 to 2020 (mean ± standard deviation in net ecosystem exchange: −548 ± 140 Tg C yr^-1^; trend: −14 Tg C yr^-1^, p<0.001), more than 30% of the region was a net CO_2_ source. Tundra regions may have already started to function on average as CO_2_ sources, demonstrating a critical shift in carbon dynamics. After factoring in fire emissions, the increasing ABZ sink was no longer statistically significant (budget: −319 ± 140 Tg C yr^-1^; trend: −9 Tg C yr^-1^), with the permafrost region becoming CO_2_ neutral (budget: −24 ± 123 Tg C yr^-1^; trend: −3 Tg C yr^-1^), underscoring the importance of fire in this region.

## Main text

Estimating terrestrial net ecosystem CO_2_ exchange (NEE) of the Arctic-Boreal Zone (ABZ) poses a significant challenge^1–4^ due to their complex functions ^4–6^ and a limited network of field measurements ^7,8^. As a result, models show a wide range of CO_2_ budgets, from substantial net atmospheric sinks (−1,800 Tg C yr^-1^) to moderate atmospheric sources (600 Tg C yr^-1^) ^1,4,5,9,10^, a concerning discrepancy as northern permafrost soils hold nearly half of global soil organic carbon stocks ^11^. The release of this soil carbon to the atmosphere as CO_2_ could significantly exacerbate climate change ^12^. Thus, there is an urgent need to improve CO_2_ budget estimates across the ABZ.

The rapid climate change of the ABZ makes this discrepancy even more critical ^13^. Increasing air and soil temperatures in both summer and non-summer seasons are causing changes in the CO_2_ budget that remain poorly understood ^9^. Furthermore, it is not known how the widespread but spatially heterogeneous increase in vegetation productivity and greening ^14,15^ impacts the annual CO_2_ balance although links to enhanced CO_2_ sinks during the spring-summer period have been found ^16,17^. Some of the enhanced uptake might be offset by CO_2_ losses associated with vegetation dieback (‘browning’), and the escalating frequency and intensity of disturbances such as abrupt permafrost thaw (e.g., thermokarst), drought and fires, further complicating the understanding of ABZ carbon dynamics and climate feedbacks ^18–20^.

Current evidence on recent ABZ CO_2_ budget trends and their main drivers is limited to few in-situ data-driven synthesis and modeling studies without a regional perspective on where and why CO_2_ budgets are changing ^1,5,9,10^. These studies have focused primarily on ecosystem CO_2_ fluxes (i.e., not incorporating fire emissions), coarse annual or seasonal CO_2_ fluxes (i.e., overlooking the intra-annual dynamics), and spatial patterns in CO_2_ fluxes with data from only one to two decades. Most importantly, earlier studies have not extended into the 2020s, a period of time where warming has further accelerated and more fires have occurred ^21^. Thus, we lack a comprehensive understanding of the regional and seasonal patterns in recent ecosystem CO_2_ fluxes, including fire emissions, and their multidecadal trends, and the links to changing environmental conditions across the ABZ.

Here, we address this knowledge gap using the most comprehensive site-level ABZ CO_2_ flux dataset to-date —including monthly terrestrial photosynthesis (gross primary production; GPP), ecosystem respiration (R_eco_), and NEE data from 200 terrestrial eddy covariance and flux chamber sites (4,897 site-months). This dataset is at least four times larger than in earlier upscaling efforts and covers a longer time period with data extending to 2020. The same dataset was previously used to analyze in-situ CO_2_ flux trends in permafrost versus non-permafrost regions, with the conclusion that the annual net uptake is increasing in the non-permafrost region but not in the permafrost region ^22^. Here we extend that study from the site level to the full ABZ region by combining flux observations with meteorological, remote sensing, and soil data, together with random forest models to estimate CO_2_ budgets across the ABZ. We do this upscaling over two periods, 2001-2020 (1-km resolution) and 1990-2016 (8-km resolution); results in the main text are based on the 1-km models unless stated otherwise. We then assess regional and seasonal patterns and trends in ABZ ecosystem CO_2_ fluxes and their environmental drivers. We also integrate annual fire emissions from 2002 to 2020 ^23^ to provide near-complete terrestrial CO_2_ budget estimates (referred as NEE + fire).

## Results

### CO_2_ budgets across the ABZ

Using machine learning models that had a high predictive performance (up to two times higher cross-validated R^2^ compared to earlier efforts ^5,9^), we find that from 2001-2020 circumpolar tundra was on average CO_2_ neutral without accounting for fire emissions (in-situ NEE: −4 ± 44 g C m^-2^ yr^-1^; upscaled NEE: 7 ± 3 g C m^-2^ yr^-1^; upscaled budget 45 ± 53 Tg C yr^-1^; mean ± standard deviation; Table 1). In contrast, the boreal was a strong sink (in-situ NEE: −42 ± 82 g C m^-2^ yr^-1^; upscaled NEE: −43 ± 7 g C m^-2^ yr^-1^; upscaled budget −593 ± 101 Tg C yr^-1^). Including fire emissions (on average 237 Tg C yr^-1^ ^23^, i.e., 2% of R_eco_ and 43% of the ABZ net CO_2_ uptake budget) changed the budget to −383 ± 101 Tg C yr^-1^ in the boreal and to 64 ± 53 Tg C yr^-1^ in the tundra. With fire emissions included, the permafrost region turned into CO_2_ neutral (NEE: −249 ± 123 Tg C yr^-1^, NEE + fire: −24 ± 123 Tg C yr^-1^).

**Table 1.** Mean gross primary productivity (GPP), ecosystem respiration (R_eco_), and net ecosystem exchange (NEE) fluxes and budgets over 2001-2020, and NEE + fire budgets from 2002-2020. Uncertainties are standard deviations across sites or pixels (for the mean fluxes) or across bootstrapped budget estimates. Positive numbers for NEE indicate net CO_2_ loss to the atmosphere and negative numbers indicate net CO_2_ uptake by the ecosystem. Mismatches in the site-level versus upscaled CO_2_ fluxes are likely related to sites being biased to certain regions and years while upscaled summaries should provide more representative regional estimates but are influenced by model performance. Mismatches in the NEE vs. GPP-R_eco_ estimates are related to different numbers of sites and observations being available for the different fluxes. Supplementary Table 1 shows the budgets for different vegetation types and regions.

| Class | In-situ average |  |  | Upscaled per-area average |  |  | Average regional budget |  |  | The proportion of summer net uptake budget of non-summer net emissions | Average regional budget with fire | Area (x 10<sup>6</sup> km<sup>2</sup>) |
| --- | --- | --- | --- | --- | --- | --- | --- | --- | --- | --- | --- | --- |
| Flux and unit | NEE g C m<sup>-2</sup> yr<sup>-1</sup> | GPP g C m<sup>-2</sup> yr<sup>-1</sup> | $R_{eco}$ g C m<sup>-2</sup> yr<sup>-1</sup> | NEE g C m<sup>-2</sup> yr<sup>-1</sup> | GPP g C m<sup>-2</sup> yr<sup>-1</sup> | $R_{eco}$ g C m<sup>-2</sup> yr<sup>-1</sup> | NEE Tg C yr<sup>-1</sup> | GPP Tg C yr<sup>-1</sup> | $R_{eco}$ Tg C yr<sup>-1</sup> | % | NEE + fire Tg C yr<sup>-1</sup> | |
| Arctic-Boreal Zone | -32 (± 76) | 618 (± 396) | 588 (± 385) | -26 (± 5) | 482 (± 20) | 460 (± 15) | -548 (± 140) | 9970 (± 144) | 9525 (± 90) | 1.4 | -319 | 20.79 |
| Tundra | -4 (± 44) | 302 (± 125) | 312 (± 133) | 7 (± 3) | 300 (± 14) | 306 (± 12) | 45 (± 53) | 2049 (± 49) | 2090 (± 33) | 0.9 | 64 | 6.8 |
| Boreal | -42 (± 82) | 705 (± 402) | 664 (± 398) | -43 (± 7) | 572 (± 24) | 537 (± 17) | -593 (± 101) | 7920 (± 106) | 7435 (± 74) | 1.6 | -383 | 13.9 |
| Permafrost region | -21 (± 62) | 458 (± 197) | 445 (± 171) | -15 (± 5) | 416 (± 20) | 405 (± 16) | -249 (± 123) | 6918 (± 109) | 6719 (± 69) | 1.2 | -24 | 16.6 |

Although the entire ABZ domain was a terrestrial CO_2_ sink across all years during 2001-2020 with an average NEE of −548 ± 140 Tg C yr^-1^, our upscaling of NEE revealed a large areal fraction of annual ecosystem CO_2_ sources across the domain (34% of the total region, Fig. 1). For the permafrost domain, the fraction of annual CO_2_ sources was even higher (41% of the region). This large fraction is also seen in our in-situ CO_2_ flux database, with 29% of sites being CO_2_ sources (NEE between 0-142 g C m^-2^ yr^-1^). These CO_2_ source sites were mostly in Alaska (44%), but also in northern Europe (25%), Canada (19%), and Siberia (13%). One key factor driving CO_2_ sources is the long and persistent non-summer season (September-May) emissions in the tundra that, on average, exceed the short summer (June-August) net CO_2_ uptake (Table 1). In the boreal, longer summers with strong uptake still dominate over non-summer emissions.

**Figure 1.**
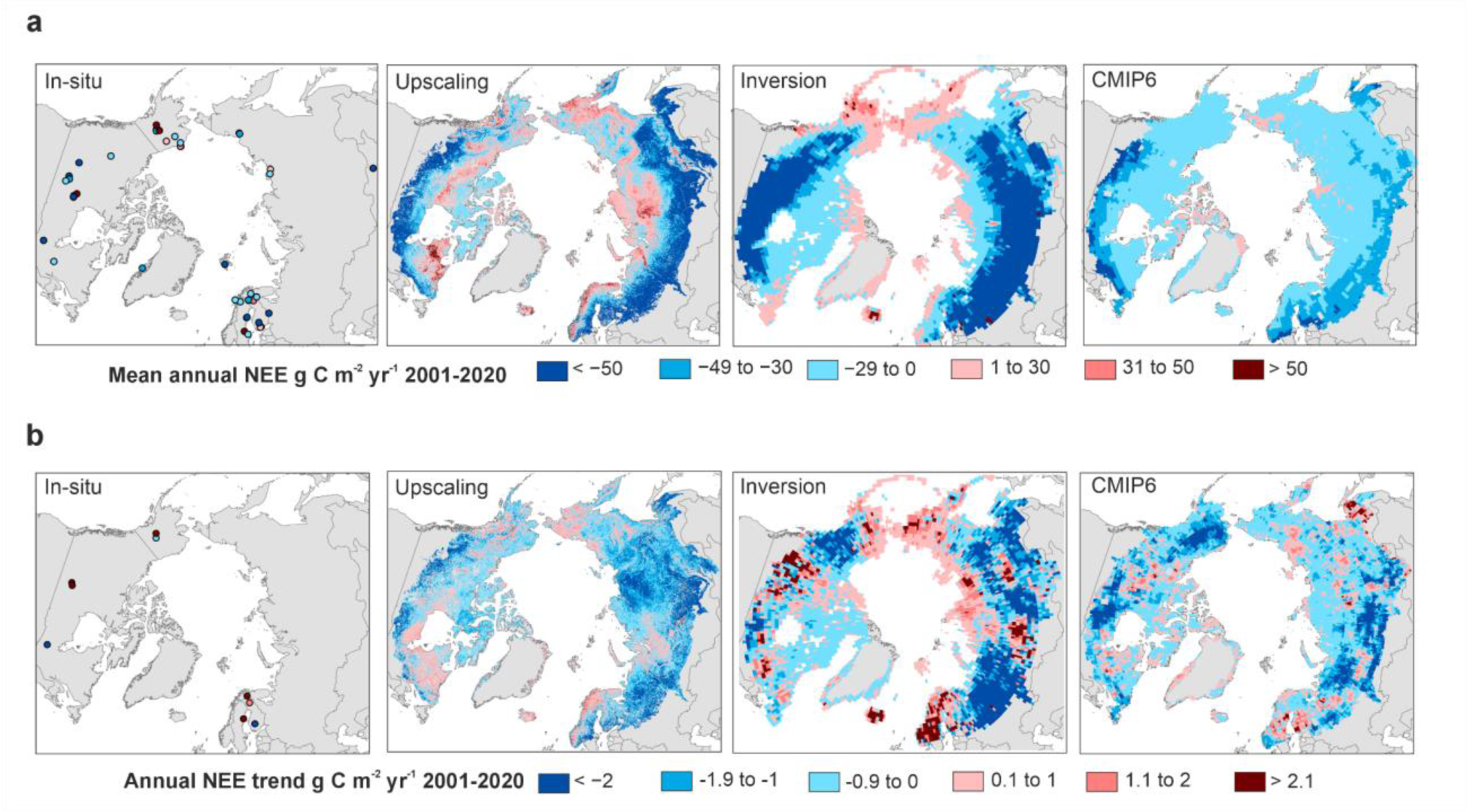
Maps showing the mean annual terrestrial NEE (a) and its trends (b) based on site-level data, our upscaling, atmospheric inversion ensemble, and CMIP6 process model ensemble. In-situ trends in b are based on sites that have >7 years of data. Supplementary Fig. 5c shows the significance of the trends. While the average upscaled NEE values can go up to 116 g C m^-2^ yr^-1^, most of the values are below 60 g C m^-2^ yr^-1^. While the NEE values of the inversion ensemble can go down to −1636 g C m^-2^ yr^-1^, most of the values are higher than −200 g C m^-2^ yr^-1^, similar to upscaling and CMIP6 model outputs.

### Model performance and comparison

We observed moderate correlation of our upscaled NEE results with an ensemble of atmospheric inversions ^24^ across space (Pearson’s correlation 0.5, p<0.001), but the correlation between the temporal trends was weaker (Pearson’s correlation 0.2, p<0.001) (Fig. 1). However, the ensemble net uptake budgets from the inversions, as well as from a global machine-learning based upscaling product (FLUXCOM-X-BASE ^25,26^) were 1.5 to 3 times larger than our upscaled budgets (Supplementary Section 5). Moreover, the global Coupled Model Intercomparison Project Phase 6 (CMIP6) process model ensemble ^27^ had barely any annual CO_2_ sources across the ABZ, indicating that the process models may not accurately simulate CO_2_ source situations (Fig. 1), especially given the prevalence of site-level sources. The cross-validated predictive performances of our random forest models for GPP, R_eco_, and NEE showed high correlations between observed and predicted fluxes (R^2^ varied from 0.5 to 0.78 and root mean square error from 19.4 to 37.3 g C m^-2^ month^-1^; Supplementary Fig. 1-3), but upscaling uncertainties remain. For example, areas with the most extensive strong sink or source estimates rarely had in-situ data and were thus largely extrapolated (e.g., sources in central Siberia, or sinks in southern Siberia, Supplementary Fig. 4). These areas also had the highest uncertainties in our analysis (approximately twice as large uncertainties as in the more densely measured areas; Supplementary Fig. 5).

### Temporal trends in upscaled ABZ CO_2_ budgets

The ABZ has been an increasing terrestrial CO_2_ sink based on NEE alone from 2001 to 2020 (temporal trend: −14 Tg C yr^-1^, p<0.001) (Fig. 2). However, the increasing sink strength was no longer statistically significant when fire emissions were added to NEE (average NEE + fire budget trend −9 Tg C yr^-1^ over 2002-2020). In the permafrost region, the NEE + fire trend was only −3.3 Tg C yr^-1^. Nevertheless, based on our NEE upscaling, 23% of the region increased (p<0.05) net CO_2_ uptake from 2001 to 2020 (Fig. 1), with increasing net sink pixels occurring across all key regions (Supplementary Fig. 31). Most of the increasing net sink activity was driven by an increase in GPP, especially in Siberia (Fig. 2). Some of the trends were also related to a declining R_eco_, likely associated with disturbed ecosystems (e.g., forest fires, harvesting) with high R_eco_ during the first post-disturbance years now recovering ^28^. However, evidence for the increasing overall net uptake trend from the in-situ data is limited due to the low number of long-term sites (>7 years of year-round measurements; 9 sites) out of which only one site showed a statistically significant trend (increasing uptake at a boreal forest site in Finland). Some of the relationships in our model are likely thus influenced by spatial differences across the sites rather than temporal and truly causal patterns, creating some uncertainty in upscaled trends ^29^. However, the model reproduces temporal patterns at individual sites well (see Supplementary Fig. 6), and our upscaled trends are similar to a recent in-situ time-series analysis ^22^ and somewhat similar to those estimated from the inversion ensemble (Fig. 1), providing confidence in our trend results.

**Figure 2.**
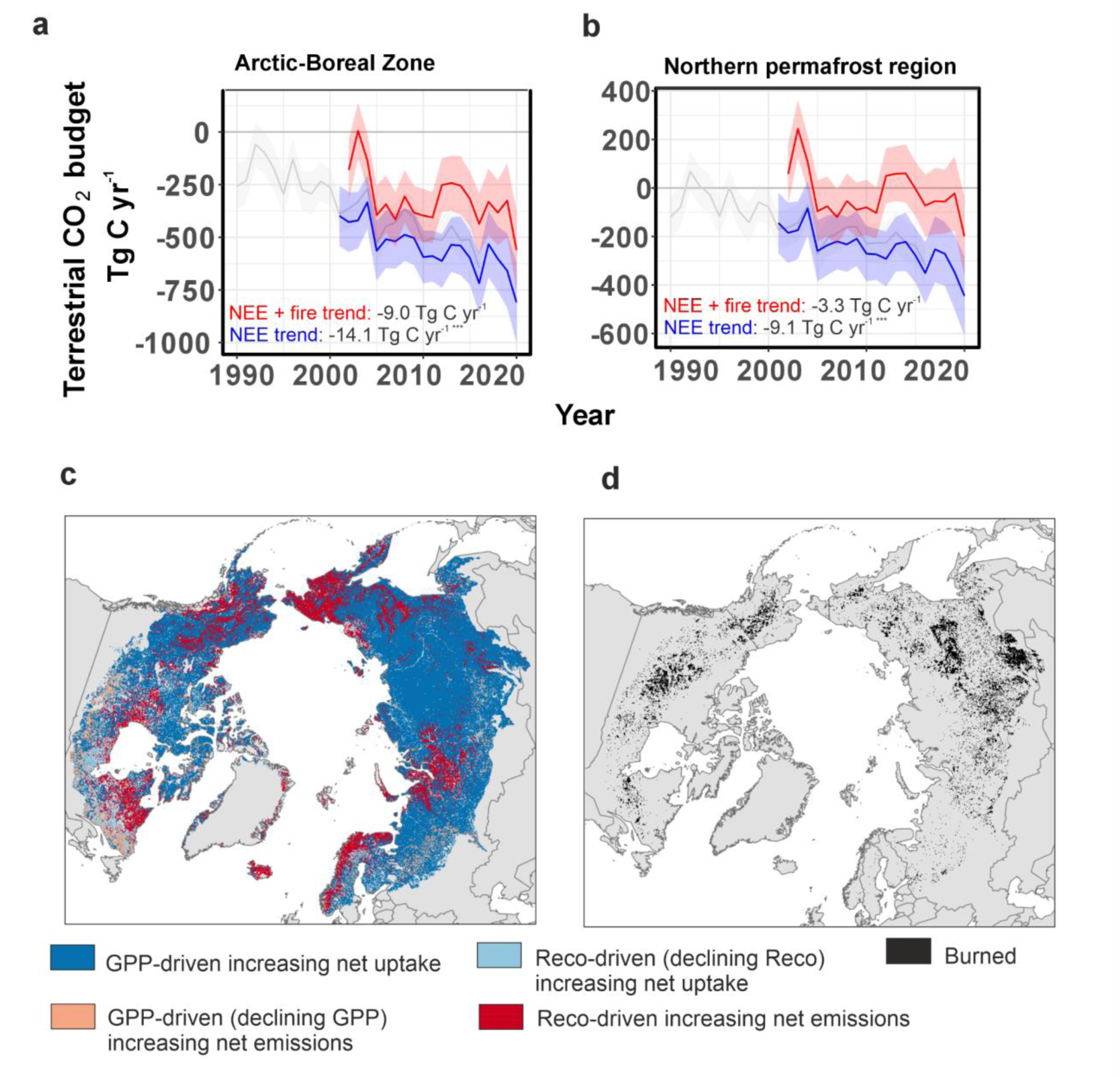
Terrestrial CO_2_ budgets for 1-km (blue; 2001-2020) and 8-km (grey; 1990-2016) NEE, and 1-km NEE + fire emissions (red; 2002-2020) across the ABZ (a) and permafrost region (b). An overlay analysis of NEE, GPP, Reco trend maps identifying how trends in GPP and Reco relate to trends in NEE over 2001-2020 (includes significant and non-significant trends; c), and a map showing also pixels burned during 2002-2020 (d). In a and b, trends are shown for the 2002-2020 (NEE + fire) and 2001-2020 (NEE) periods. Stars in the trend values depict the significance of the trend (*= p<0.05, **=p<0.01, ***=p<0.001).

Parts of the ABZ also show increasing annual net CO_2_ emissions over time (Fig. 1). Such trends have been observed at six long-term sites (2 to 17 g C m^-2^ yr^-1^, p>0.05), and in 2% of the upscaled region (p<0.05) from 2001 to 2020. Most of the increasing net emission trends were driven by an increase in R_eco_ instead of a decline in GPP (Fig. 2). Regions experiencing increased net CO_2_ emissions in upscaling were found especially in (i) northern Europe and Canada (dominated by evergreen needleleaf forests with mild and moderately wet climates), (ii) parts of central Alaska and northern Siberia (sparse boreal ecosystems and graminoid tundra with permafrost and high soil carbon stocks), and (iii) Hudson Bay and Siberian lowlands (wetlands with some permafrost and high soil organic carbon stocks) (Supplementary Fig. 31). Some sites in Alaska have increasing net emissions of CO_2_ due to permafrost thaw ^18,30^, but it is unclear if similar changes are occurring in other regions with increasing net CO_2_ emissions.

We calculated an overall 25% increase in seasonal amplitude of CO_2_ fluxes from the upscaled NEE time series from 2001 to 2020 across the ABZ, on par with earlier atmospheric and modeling studies ^31,32^. Both increasing summer uptake and non-summer season emissions—the key dynamics driving increasing annual sinks and sources—were evident in the tundra and boreal biomes (Fig. 3). However, over the 2001-2020 period, the increasing uptake (GPP) during summer months dominated over increasing net emissions (R_eco_) during non-summer months across most of the domain. On average across both biomes, net uptake increased the most during July (an average upscaled increase of −5 g C m^-2^ month^-1^ in the boreal and −3 g C m^-2^ month^-1^ in the tundra in 2011-2020 compared to 2001-2010), and increasing net emissions were occurring throughout the entire non-summer season, with no clear peaks (0.1-0.9 g C m^-2^ month^-1^). Although increases in early growing season (May-June) uptake were evident, late growing season (September) trends were absent or minimal (Fig. 3).

**Figure 3.**
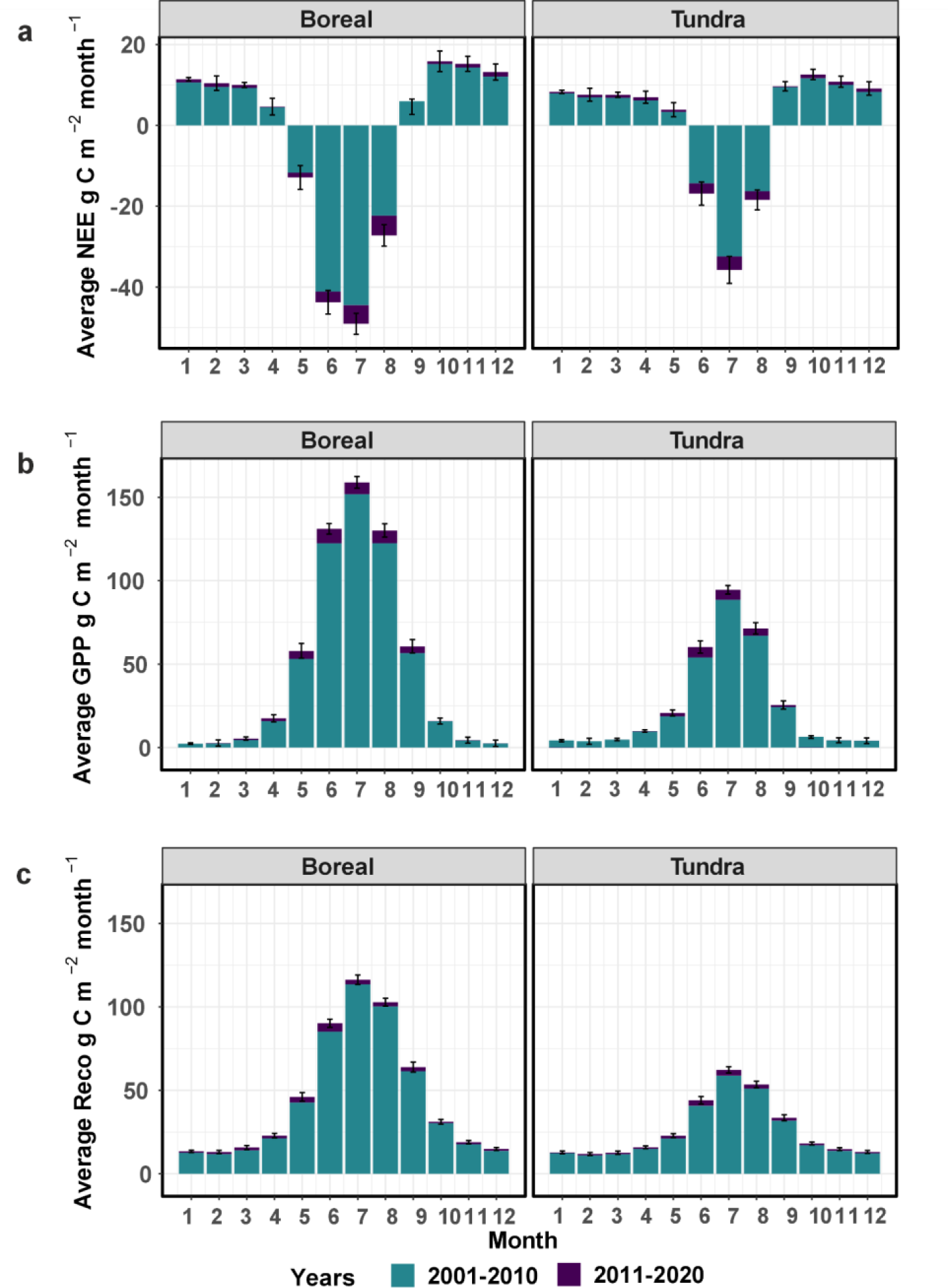
Average upscaled monthly NEE, GPP, and R_eco_ in boreal and tundra biomes during the past two decades. Negative NEE values represent net uptake and positive net release. Error bars are only shown for the 2011-2020 period but are similar for the 2001-2010 period. Note that NEE was 1.4 g C m^-2^ month^-1^ lower in September 2011-2020 compared to 2001-2020 in the boreal biome, but this is not shown in the figure. For a similar figure made based on the in-situ data, see Supplementary Fig. 7.

### Drivers of ABZ CO_2_ fluxes

There are several environmental conditions driving CO_2_ budgets across the ABZ. Our variable importance analysis showed CO_2_ fluxes, and thus the overall increasing sink strength, are explained by dynamic variables of air or surface temperatures, solar radiation, the Normalized Difference Vegetation Index (NDVI), and partially also by soil temperature, snow cover, and the vapor pressure deficit (Supplementary Fig. 8-10). Other less important dynamic variables were vegetation cover and atmospheric CO_2_ concentration. Volumetric soil water content was not important in our models, likely due to the large uncertainties and coarse spatial resolution in the gridded product, although in-situ studies have shown drier soils to be linked to larger net CO_2_ emissions and wetter soils to enhanced plant growth due to the lack of water limitation ^33^. Static variables (primarily vegetation type, soil carbon stock, soil pH) were also important in explaining spatial differences.

The most important dynamic variables had a positive overall effect on net uptake, GPP, and R_eco_ (Supplementary Fig. 8-10), however, these relationships are more nuanced in reality. In fact, the recent permafrost in-situ trend analysis of CO_2_ fluxes using the same database suggests that the CO_2_ flux response to warmer temperatures ranges from positive to negative, depending on the availability of water and nutrients at the site ^22^. Consequently, strong warming or greening trends did not always translate into increasing net CO_2_ sinks in our upscaling (Supplementary Fig. 11). For example, while 49% of the region experienced greening (June-August average NDVI; based on MODIS NDVI, p<0.05), only 12% of those greening pixels showed an annual increasing net CO_2_ uptake trend, and 29% an increasing June-August net uptake.

### Continental and regional patterns in CO_2_ budgets and their trends

Our upscaling showed clear continental patterns in NEE budgets and trends (Fig. 4), with the boreal biome primarily driving the budget and trend differences between the continents ^1,34^. The increasing net uptake trend was more pronounced in Eurasia (−11 Tg C yr^-1^, p<0.001) compared to North America (−3 Tg C yr^-1^, p<0.05), which corresponds with the smaller area and weaker warming, declining snow cover and greening trends in North America (Supplementary Fig. 12-14). We found statistically significant declining summer soil moisture trends in Siberian boreal (Supplementary Fig. 15), but this did not translate into stronger net emissions. When fire emissions were added, continental differences were less pronounced due to the much larger and more rapidly increasing CO_2_ emissions from Siberian fires (on average 160 compared to 76 Tg C yr^-1^ in North America; Supplementary Fig. 16). Fire emissions even reversed some NEE trends: the strong increasing sink in Siberia became a source when fire emissions were included (trend: +0.7 Tg C yr^-1^; p > 0.05). However, Siberian ecosystems have the largest uncertainty for both the upscaled fluxes and inversion-based estimates due to lack of in situ observations, making it challenging to accurately determine the magnitude of continental differences (Fig. 4, Supplementary Fig. 17).

**Figure 4.**
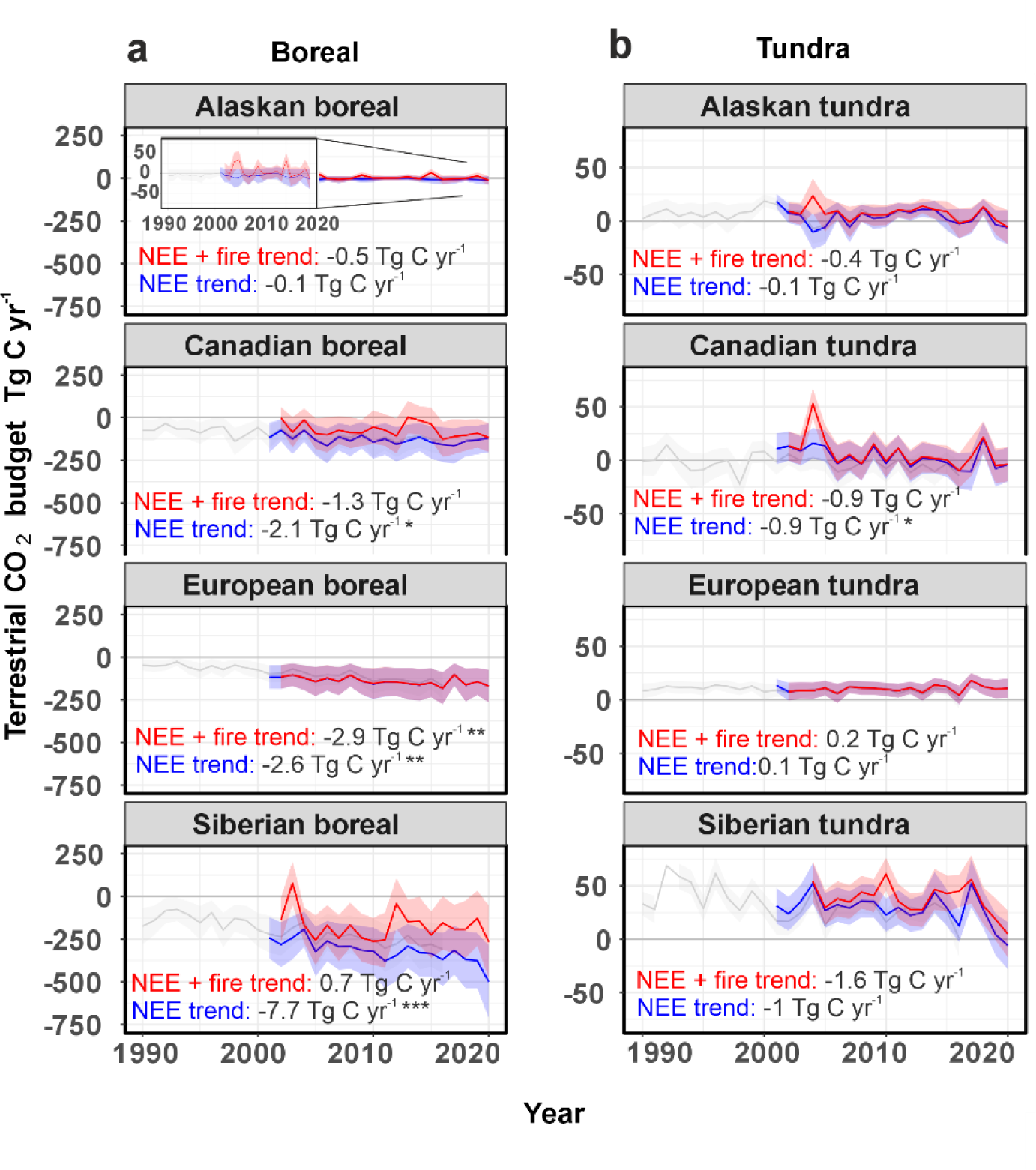
Terrestrial CO_2_ budgets for NEE and NEE + fire in key regions and biomes across the boreal (a) and tundra (b). Terrestrial CO_2_ budgets are shown for 1-km (blue; 2001-2020) and 8-km (grey; 1990-2016) NEE, and 1-km NEE + fire emissions (red; 2002-2020). Trends are shown for the 2002-2020 (NEE + fire) and 2001-2020 (NEE) periods. The inset in Alaskan boreal in (a) shows the time series with a narrower y axis compared to the main figure to better detect interannual variability. Stars in the trend values depict the significance of the trend (*= p<0.05, **=p<0.01, ***=p<0.001). Fire emissions alone are shown in Supplementary Fig. 16.

Alaska is an important contributor in the weaker North American CO_2_ sink. Based on our analysis, Alaska as a whole was consistently CO_2_ neutral or a source over 2002-2020 (NEE + fire emissions), both in the boreal (budget +5 Tg C yr^-1^) and tundra (budget +7 Tg C yr^-1^). Alaska has a relatively high density of observations, making this result more certain compared to other regions. Alaska is therefore different from the other ABZ regions where boreal regions still remain on average CO_2_ sinks. Potential reasons for the Alaskan CO_2_ source include Alaska having the most rapidly warming autumns and declining autumn snow cover, which also have high inter-annual variability (Supplementary Fig. 14 and 18). Further, field measurements suggest that many of the observed changes in Alaskan ecosystems can be attributed to permafrost thaw ^18,30^—a phenomenon that has accelerated significantly in response to Alaska’s pronounced warming trend since the 1950s ^35^. However, we were unable to incorporate permafrost thaw into our models as high-resolution geospatial data from 1990 to 2020 were not available. The question of whether analogous trends will manifest in other regions across the northern permafrost region remains an important research priority.

## Discussion

Our results show that the ABZ was on average an increasing terrestrial CO_2_ sink (GPP is increasing more than R_eco_ + fire), indicating that the region still creates an important negative feedback to global warming. However, our study also suggests some positive feedbacks to climate change that have been more regional and of shorter duration in recent decades. We show that the presence of annual sources was large as indicated by several site-level and regional studies ^36–38^, and even larger with fire emissions included ^39^. There were also extreme years when fire emissions exceeded annual net CO_2_ uptake (e.g., 2003 in Siberian boreal, 2012 in Canadian boreal, and several years in the permafrost region; Fig. 2). We also observed increasing shoulder season net emission trends, particularly in Alaska ^40^ (Supplementary Fig. 34). Moreover, while summer net uptake increase still dominates over non-summer CO_2_ emissions, net CO_2_ uptake is increasing only in the early and peak growing season (May-August in the boreal and June-August in the tundra) and not in the late growing season (September), because GPP does not increase later in the season due to plant physiological limitations, and drier and warmer conditions cause enhanced R_eco_ instead ^41–46^. A better understanding of how soil moisture and hydrology have been and will be changing, and the impact of these changes on CO_2_ fluxes is critical for more accurate ABZ CO_2_ budgets.

In the tundra, our findings reveal a noteworthy shift in carbon dynamics. While the tundra region has been a carbon sink for millennia ^47^, our results suggest that many tundra regions may now have started to function as CO_2_ sources. This transition from an ecosystem CO_2_ sink to a CO_2_ source may have begun prior to 1990 ^48^, yet the precise timing of this transformation remains uncertain. The main drivers of this pattern may be related to warming-induced permafrost thaw, the drying of soils, or vegetation shifts ^49–51^ but remain unresolved. Tundra regions are also progressing towards conditions where average annual soil (0-7 cm) temperatures are above freezing, resulting in more soil organic material being susceptible to decomposition (Supplementary Fig. 12). Overall, the primary reason behind the annual CO_2_ emissions from tundra ecosystems is the limited duration of the high net CO_2_ uptake period, and the substantial non-summer season net emissions. However, we observed lower in-situ and upscaled October-April season NEE fluxes and budgets compared to Natali et al. (2019) throughout the entire period (Supplementary Fig. 19; ^52^).

Our results demonstrate the need to further study Siberian CO_2_ flux trends. Our upscaling indicated that some of the strongest net sources and sinks, and strongest increasing sink trends occur in the Siberian boreal. Increasing sink trends in Siberian tundra were also the strongest across tundra regions. The Siberian sink trend might be explained by strong greening trends ^53^, earlier growing season starts and increasing carbon uptake due to declining spring snow cover (Supplementary Fig. 14), increases in tree growth and distribution ^54,55^, rapid recovery of ecosystems after fire ^56^, and high cover of larch forests that can rapidly take up CO_2_ (Supplementary Table 1)^8,57^. However, the large inversion model spread, sparse measurement network, and our upscaling uncertainties indicate that it remains challenging to conclude the magnitude of the Siberian CO_2_ balance ^2^. This is a significant problem given that Siberia stores more than half of the permafrost region’s C stocks and is now warming more rapidly than other ABZ regions.

In summary, our study reveals distinct spatial and temporal patterns in CO_2_ budgets across the ABZ and underscores the importance of three decades’ worth of data. Relatively robust spatial patterns can be seen, such as the Alaskan CO_2_ sources and southern Eurasian boreal sinks while the temporal trends remain more uncertain. While CO_2_ fluxes can be relatively well modeled using machine learning and existing gridded datasets, gaps persist, such as the incomplete characterization of fire, thermokarst and harvesting disturbances and their links to ecosystem CO_2_ fluxes, the lack of accurate predictors describing soil moisture ^1^, and the need to quantify landscape heterogeneity and carbon dynamics at even higher spatial (meters) and temporal resolutions (days). Sustaining long-term sites is crucial to accurately track trends in ABZ CO_2_ balance, while establishing new year-round sites in data-poor areas like Siberia and the Canadian Arctic is vital to fill knowledge gaps and enhance our understanding of carbon dynamics ^58^.

## Online Methods

### In-situ data overview

We used a recently compiled dataset of in-situ Arctic-boreal terrestrial ecosystem CO_2_ flux measurements (ABCflux, led by Virkkala et al. 2021 ^8^) within the ABZ (Supplementary Methods Section 1). The synthesized data were cumulative flux densities of gross NEE, GPP, and R_eco_ aggregated at the monthly timescale (3,968 to 4,897 site-months depending on the flux). In addition to eddy covariance data, we included fluxes estimated with the chamber method to increase data coverage especially during the growing season. The dataset included metadata out of which we used the site coordinate, biome, flux measurement method, and disturbance history information in the analysis. For further details on the dataset, see Virkkala et al. (2021) ^8^ and a description of additional data processing and screening in the Supplementary Methods Section 2. Note that our study does not include lateral transport of carbon, or vertical lake and river CO_2_ emissions which were recently summarized to be 93, 66, and 164 Tg C yr^-1^, respectively, in the northern permafrost region (i.e., greater than the NEE+fire budget calculated in this study) ^59^.

This dataset is more comprehensive than the ones used in earlier upscaling studies as it represents monthly fluxes from the entire year if available, while Virkkala et al. (2021) focused on coarse seasonal or annual fluxes ^5^, Natali et al. (2019) on monthly winter fluxes ^9^, and Mu et al. (2023) a more limited temporal period (2002-2017) ^60^. Furthermore, we included more data from recent years (805 monthly observations from 2015-2020 compared to 32 and 95 fluxes in Virkkala et al. 2021 and Natali et al. 2019, respectively), and the sample size in our models was 4 to 25 times larger here compared to the earlier upscaling efforts.

### Geospatial data

We used data from geospatial products as predictor variables to upscale fluxes. Our models had the following predictors: month, incident solar radiation, vapor pressure deficit, atmospheric CO_2_ concentration, vegetation type, snow cover (the fraction covered by snow), soil temperature (0-7 cm), soil moisture (0-7 cm), NDVI (MODIS- or AVHRR-based), land surface temperature (or air temperature; MODIS- or ERA5 Land-based), compound topographic index (i.e., topographic wetness index), continuous vegetation fields describing percent non-tree vegetation and non-vegetated fraction and percent tree cover (MODIS- or AVHRR-based), soil pH (0-5 cm), soil organic carbon stock in 0-2 meters, and permafrost probability. In our analysis, NDVI was the primary predictor describing disturbances, with declines in NDVI being related to disturbances ^5^. Data were in daily, weekly, monthly, annual, and static format (i.e., no temporal changes such as in the compound topographic index). If data were of higher temporal resolution than monthly, they were aggregated to monthly time steps. Gaps in MODIS and AVHRR NDVI time series were filled to produce a continuous time series. Data were re-projected to North Pole Lambert Azimuthal Equal Area Projection at 1 and 8 km spatial resolution and extracted at the flux sites. See Supplementary Section 3 for further descriptions and data sources.

We used the Global Fire Emissions Database (GFED) 500-m fire product ^23^ to calculate fire emissions. The product is based on a global fire emissions model with a spatial resolution of 500 m using MODIS data. The model was developed using an updated field measurement synthesis database of fuel load and consumption which included improvements, for example, in boreal soil carbon combustion. The higher resolution of the 500-m model compared to earlier coarser models improved the detection of small-scale fires and understanding of landscape heterogeneity, and reduced the scale mismatch in comparing field measurements to model grid cells. However, some small fires might still be undetected by this model, leading to potential underestimations in carbon emissions in this product.

### Machine learning modeling

We used random forest models to upscale GPP, R_eco_, and NEE to the ABZ from 1990 to 2020, the period with in-situ flux measurements. Two sets of predictive models were developed: (i) models using primarily predictors with a spatial resolution ≤1 km from 2001 to 2020 (i.e., the MODIS era) at 1-km spatial resolution (hereafter 1-km models;), and (ii) models using coarser-resolution predictors from the AVHRR GIMMS era (1990-2016;) from 1990 to 2016 at 8-km spatial resolution (hereafter 8-km models) (Supplementary Table 3). Each model included all available monthly fluxes from the entire year, i.e. no separate models for individual months or seasons were developed, as this approach resulted in the best predictive performance. All models included 17 predictors, but the sample sizes were variable depending on the amount of data available for each flux and time period; NEE models had the highest amount of model training data compared to GPP and R_eco_ models. For the 1-km model, coarsest predictors were at 9-km resolution but most important predictors were at 1 to 4-km resolution. For the 8-km model, the coarsest predictor resolution was 9 km, and the most important variables had a resolution of 1 to 9 km.

Model parameter tuning was performed based on leave-one-site out cross validation (CV) to achieve minimum predictive error. The models were run using the “caret” package in R version 4.2 ^61^. We assessed the predictive performance of the final models using the (1) R^2^, (2) root mean square error (RMSE), 3) mean absolute error (MAE), and 4) mean bias error (MBE) between predicted and observed values using the CV data. Larger RMSE and MAE values indicate larger errors, and positive MBE values indicate overestimation. The predictive performance of our models was good or high, with R^2^ ranging from 0.55 to 0.78 and RMSE from 19.4 to 37.3 g C m^-2^ month^-1^, but there were uncertainties in model performance at both ends of the flux gradient. Specifically, the model had a tendency to slightly overestimate fluxes, which was particularly evident with the model struggling with strong sink observations (observations of ca. −180 to −80 g C m^-2^ month^-1^ were predicted to be −80 to −20 g C m^-2^ month^-1^ in cross validation; Supplementary Fig. 1). Other uncertainties were mostly due to 1) our model not being able to identify landscape heterogeneity with nearby sites showing large differences in CO_2_ fluxes (e.g., a forest and wetland site), and 2) our model not capturing inter-annual variability at individual sites, both of which are likely attributed to the coarse, uncertain, and missing predictors characterizing such conditions (e.g., soil moisture, disturbances) (Supplementary Fig. 1-3).

We evaluated the uncertainty of predictions by creating 20 bootstrapped model training datasets (with replacement; same sample size as in the original model training data) and using those to develop 20 individual models and predictions. Out of the 20 predictions, we calculated the standard deviation to represent prediction uncertainty. The uncertainty ranges in NEE + fire budgets only represent NEE uncertainties. We further assessed the area of extrapolation of the models, and the influence of the flux measurement method and disturbance history information on flux predictions by training models with different subsets. Further details of the uncertainty analyses can be found in the Supplementary Methods Section 4.

### Model performance in burned ecosystems

In addition to direct fire emissions from combustion (i.e., burning) derived with GFED, fires have a profound impact on carbon budgets by modulating post-fire ecosystem CO_2_ fluxes ^28,67^. Our current database has 21 sites that reported fire disturbance. Only four of those were longer-term sites (operating for >3 years) with recent (<10 years since burn) fire history. All four of these sites in young recovering ecosystems were measured year-round and originally had an evergreen (black spruce) forest cover which underwent a shift to a more deciduous shrub and tree-dominated cover after a stand-replacing fire. These include (i) CA-NS7 with 4 years of data starting 4 years after the fire, (ii) CA-SF3 with 6 years of data starting 3 years after the fire, (iii) US-Rpf with a 6-year time series starting 4 years after the fire, and (iv) US-CR-Fire with a 4-year time series starting the next year after the fire.

Across all the burned sites, the in-situ flux data and remotely-sensed NDVI time series show a clear pattern of July NDVI values, GPP and net carbon uptake steadily increasing after the fire (Supplementary Fig. 24). This post-fire recovery signal is captured by our upscaling, as we see our upscaled GPP and net uptake drop after a fire, and then return to higher levels after the fire (Supplementary Fig. 24 and 25). However, while our random forest model fits the time series of the longer-term sites with recent fire history relatively well, the predictions based on cross validation (i.e., model training data excluding each site) are variable (Supplementary Fig. 26), indicating that our model might struggle in extrapolating post-fire ecosystem CO_2_ fluxes in other areas. The model performance at sites experiencing recent fire or other disturbances was also lower than at sites without disturbance or disturbance information, as the model had a lower R_2_ and a tendency to underestimate NEE values (i.e., underestimate net CO_2_ emissions or overestimate net CO_2_ uptake) (Supplementary Fig. 27).

### Spatial upscaling of fluxes

We upscaled fluxes across the Arctic-boreal terrestrial area ≥49° N ^62^, which comprises 20.69 × 10^6^ km^2^ of land (excluding glaciers and ice sheets; Fig. 1) with lake and glacier areas removed. The models were applied at a monthly time step from 2001 to 2020 for the 1-km models and from 1990 to 2016 for the 8-km models.

We analyzed the upscaled flux maps as well as fire emission and environmental predictor rasters for temporal trends using the nonparametric Mann–Kendall test using the “zyp” package ^63,64^ with pre-whitening (Zhang method ^65^) to remove autocorrelation. We report the significance of Kendall’s correlation coefficient (the strength of the time-series) and the Theil–Sen slope to describe trends over time. Finally, we calculated zonal statistics of average annual, seasonal, and monthly fluxes and trends across key regions (Siberia defined as all land east from the Ural mountains, including a small portion of Mongolia; the rest of Eurasia, including Greenland are grouped within the European classes), biomes (tundra and boreal) ^62^, permafrost region ^66^, and vegetation types ^8^.

### Comparison to process models and atmospheric inversions

We compared our estimates with the CMIP6 process models ^27^, atmospheric inversions used in the Global Carbon Project’s Global Carbon Budget 2022 ^68^, and a global upscaling product FLUXCOM-X-BASE ^26^. We included a subset of CMIP6 process models (13 in total) that had soil thermal processes at several depths to assure they had some information about the freeze-thaw patterns in the permafrost region. We included inversions with data from the whole 2001-2020 period (i.e., included five inversions and excluded four). Fire CO_2_ emissions ^23^ were subtracted from the inversions. CMIP6 process model outputs were only available for the 2001-2014 period. The final model outputs used here represent terrestrial NEE (GPP-R_eco_) in a similar way across the models except for inversions that also include vertical CO_2_ fluxes from water bodies. There is some heterogeneity between individual inversions and CMIP6 models within the ensembles, but overall the ensemble results can be considered robust ^69,70^.

## Supporting information

Analysis codes

## Acknowledgements

This work was supported by funding from the Gordon and Betty Moore Foundation (grant #8414) and funding catalyzed by the TED Audacious Project (Permafrost Pathways). We additionally acknowledge the funding from the NASA Arctic-Boreal Vulnerability Experiment and Carbon Cycle Science programs (NNX17AE13G), NSF PLR Arctic System Science Research Networking Activities (RNA, Grant#1931333), Minderoo Foundation, NSF (grants DEB LTREB 1354370, 2011257, DEB-0425328, DEB-0724514, and DEB-0830997), the US Geological Survey Climate R&D Program, NSF Arctic Observatory Network (grants 1936752, 1503912, 1107892), KAKENHI(19H05668),The Danish National Research Foundation (CENPERM DNRF100), EU HORIZON GreenFeedBack, grant agreement No. 101056921, Danish Arctic Climate support through Greenland Ecosystem Monitoring and ICOS grants, Natural Sciences and Engineering Research Council, the Deutsche Forschungsgemeinschaft (DFG, German Research Foundation) under Germany‘s Excellence Strategy – EXC 2037 ‘CLICCS - Climate, Climatic Change, and Society’ – Project Number: 390683824, NASA Grant/Cooperative Agreement Number: NNX17AD69A, The Research Council of Norway (BioGov, project nr. 323945), ERC synergy project Q-Arctic (grant agreement no. 951288), the Copernicus Atmosphere Monitoring Service, implemented by ECMWF on behalf of the European Commission (Grant: CAMS2 55), the Environment Research and Technology Development Fund of the Environmental Restoration and Conservation Agency of Japan (JPMEERF21S20810), ArCSII(JPMXD142031886), Financial support from the Swedish Research Council (VR) and consortium partners to ICOS Sweden (grants 2015−06020 and 2019−00205) and SITES (grant 2017−00635), VR grant 2019-04676 and 2018-03966, ArcticNet and NSERC,NOAA Cooperative Agreement NA16SEC4810008, Research Council of Finland (NPERM project nrs 341349, 330840, 349503 ICOS-FIRI), ICOS-FI via University of Helsinki funding, the EU Horizon Europe (GreenFeedback nr. 101056921 and LiweFor nr. 101079192), NRF-2021M1A5A1065425 (KOPRI-PN23011), NRF-2021M1A5A1065679, NRF-2021R1I1A1A01053870, the Dutch Research Council (NWO) (project number VI.Vidi.213.143), and The Natural Environment Research Council through the National Centre for Earth Observation (NE/R000115/1). Part of the research was carried out at the Jet Propulsion Laboratory, California Institute of Technology, under a contract with the National Aeronautics and Space Administration (80NM0018D0004). Part of the inverse analyses were performed on the supercomputer systems at the National Institute for Environmental Studies and Meteorological Research Institutes (NEC SX-Aurora TSUBASA and FUJITSU PRIMERGY CX2550M5) and at the HPC cluster Aether at the University of Bremen, financed by DFG within the scope of the Excellence Initiative (Germany).

## Data availability

In-situ data used here can be accessed from ORNL DAAC ^71^ and geospatial data from the links and references provided in the Supplementary Tables 3 and 6. The 1-km and 8-km upscaled rasters of NEE, GPP, and R_eco_ together with their uncertainties will be published via ORNL DAAC.

## Code availability

The main analysis codes can be found in the Supplement.

