## Supplementary Materials and Methods for "An increasing Arctic-boreal CO_2_ sink offset by wildfires and source regions"

### Supplementary Material and Methods for “An increasing Arctic-boreal $CO_2$ sink offset by wildfires and source regions”

Anna-Maria Virkkala, Brendan M. Rogers, Jennifer D. Watts, Kyle A. Arndt, Stefano Potter, Isabel Wargowsky, Edward A. G. Schuur, Craig See, Marguerite Mauritz, Julia Boike, Syndonia M. Bret-Harte, Eleanor J. Burke, Arden Burrell, Namyi Chae, Abhishek Chatterjee, Frederic Chevallier, Torben R. Christensen, Roisin Commane, Han Dolman, Bo Elberling, Craig A. Emmerton, Eugenie Euskirchen, Liang Feng, Mathias Goeckede, Achim Grelle, Manuel Helbig, David Holl, Järvi Järveoja, Hideki Kobayashi, Lars Kutzbach, Junjie Liu, Ingrid Liujkx, Efrén López-Blanco, Kyle Lunneberg, Ivan Mammarella, Maija E. Marushchak, Mikhail Mastepanov, Yojiro Matsuura, Trofim Maximov, Lutz Merbold, Gesa Meyer, Mats B. Nilsson, Yosuke Niwa, Walter Oechel, Sang-Jong Park, Frans-Jan W. Parmentier, Matthias Peichl, Wouter Peters, Roman Petrov, William Quinton, Christian Rödenbeck, Torsten Sachs, Christopher Schulze, Oliver Sonnentag, Vincent St.Louis, Eeva-Stiina Tuittila, Masahito Ueyama, Andrej Varlagin, Donatella Zona, and Susan M. Natali

### 1. Study domain

Our study covers the Arctic-Boreal zone (ABZ) which was delineated based on the tundra and boreal biomes included in Dinerstein et al. (2017) [1]. The tundra consists of treeless Arctic and sub-Arctic ecosystems, and the boreal biome is dominated by forests. Both of the biomes also include wetlands. In total, 23% of the ABZ is covered by larch forest, 38% by other types of forests (evergreen, mixed, deciduous broadleaved), 6% by wetlands, and 14% by sparse boreal vegetation that is not classified as a forest, 13% as tundra vegetation covered by shrubs, grasses, mosses, and lichens, and 6% as barren tundra with minimal vegetation, based on the land cover dataset used in this study (Supplementary Table 3). 77% of the ABZ contains permafrost which is ground that remains frozen for at least two consecutive years [2]. Our study does not focus on the entire permafrost region that also includes the alpine permafrost regions further south in the Tibetian plateau, Alps, and Rockies mountain chains, for example.

### 2. In-situ flux data summary

#### 2.1. Data screening and filtering

We used the full Arctic-boreal $CO_2$ flux (ABCflux) database [3,4] with the following modifications. In instances where chamber plots from the same site had the same coordinates, we calculated an average flux for each year and month across the chamber plots (1-12 plots per site). This was done to assure that these measurements better represent average landscape-scale conditions of the site that could be more easily linked to the geospatial data sets and compared with eddy covariance measurements. We further filled flux columns that remained NA even though the two other flux columns had data by subtracting the two flux values using variations of the equation NEE= GPP-$R_{eco}$. We only included sites within our Arctic-boreal domain (i.e. tundra and boreal biomes as defined in Dinerstein et al. (2017) [1]), thus a few hemiboreal sites were excluded.

We removed outlier observations showing extremely high July NEE uptake in the tundra with the Study_ID_Short identifier “Lund_Kobbefjord_Ch” with uptake values < -300 g C $m^{-2}$ $month^{-1}$ in Greenland (range of tundra July NEE primarily between -25 to -100 g C $m^{-2}$ $month^{-1}$). Moreover, we removed observations with the Study_ID_Short “Goulden_CA-NS2_tower2”, “Goulden_CA-NS3_tower3”, “McCaughey_CA-Man” showing high GPP values in the peak winter months (GPP 100-300 g C $m^{-2}$ $month^{-1}$ in January or February compared to the overall December-February range between -20 to 74 g C $m^{-2}$ $month^{-1}$). The final list of sites can be found in Supplementary Table 2.

#### 2.2 Data description

The final data used in 1-km models included 199 sites and 4,981 months in total. The sample sizes for the different fluxes and model resolutions can be found in Supplementary Table 4. The majority of the data for the 1-km models was based on eddy covariance: 55% of sites and 88% of months represented this approach. Each site had from one to 213 months of measurements in our database, with the average number of months per site being 25. Long-term sites with more than 7 years of year-round data included boreal forest sites (FI-Hyy, FI-Sod, CA-Oas, CA-Obs, CA-Gro, US-Uaf, SE-Deg), a wetland site (FI-Kaa), and a tundra site (US-EML). Across all the sites, most of the sites were in recently undisturbed ecosystems (i.e., ecosystems without known abrupt changes related to, e.g., fires, forest harvesting, thermokarst). There were 21 sites that had experienced a fire; two of these had data from recent years (from Alaska). 3 sites reported thermokarst; 2 of these were in Alaska and 1 in Siberia, but gradual permafrost thaw was present in many more sites. At least 5 forest sites had been harvested. In total, 14% of the sites in ABCflux have experienced some level of natural or anthropogenic disturbances. This proportion is likely less than the overall proportion of disturbances across the entire ABZ. For example, 11% of the ABZ was burned during the 2002-2020 period [5], and the areas experiencing thermokarst and harvesting are also extensive. Thus, the flux site distribution might be biased towards non-disturbed or only moderately disturbed sites [6,7], leading to potential underestimations in disturbance effects on $CO_2$ emissions.

The most represented vegetation types in our database were evergreen needleleaf forests (26% of sites and 41% of months), shrub tundra (15% of sites and 10% of months), and wetlands (11% of sites and 15% of data). Note that these vegetation type statistics were based on the information extracted from the gridded land cover data set and were thus slightly different from those reported in Virkkala et al. (2021).

#### 2.3. A description of the in-situ flux variability

The distribution of monthly $CO_2$ fluxes was not normal; Shapiro-Wilk normality test for NEE was W = 0.88015, p-value < 0.001, for GPP W = 0.79242, p-value < 0.001, and for Reco W = 0.7835, p-value < 0.001. For NEE, the tails of the distribution are extending towards the sink (Supplementary Fig. 28). Average net $CO_2$ uptake during 2001-2020 was highest in July (Supplementary Fig. 7), both in the tundra and boreal, during which time almost all observations (95%) were net sinks. In the tundra, net uptake was high primarily in July while it was high during all the summer months in the boreal: during June and August in the tundra, the rate of carbon uptake was two to three times less compared to boreal ecosystems (Supplementary Fig. 7). Non-summer season (September-May) net emissions were highest in October, both in the tundra (average in-situ NEE 13 ± 12 g C $m^{-2}$ $month^{-1}$) and the boreal regions (average in-situ NEE 17.7 ± 17 g C $m^{-2}$ $month^{-1}$). In both biomes, average non-summer season NEE and $R_{eco}$ were greater than zero in all months (excluding May in the boreal). 8% of the in-situ fluxes also show net uptake in autumn and spring months (Supplementary Fig. 19; excluding May). The magnitude of average monthly $CO_2$ fluxes was relatively similar in both biomes across the peak winter months (December-February, in-situ NEE ranging from 9 to 14 g C $m^{-2}$ $month^{-1}$ in the boreal and 7 to 8 g C $m^{-2}$ $month^{-1}$ in the tundra). Furthermore, net emissions in the tundra in May and September were even higher than in the boreal.

It is worth noting that while we had up to 10 years of data from three Siberian larch forest sites, the only year-round site was located in an ecotone close to the tundra experiencing permafrost thaw, and was a small annual $CO_2$ source [8]. The other two larch sites had some of the strongest growing season fluxes that might also indicate strong annual $CO_2$ sinks (e.g., -240 g C $m^{-2}$

month$^{-1}$ for July compared to ca. -150 g C m$^{-2}$ month$^{-1}$ for the evergreen forests). Furthermore, Siberian tundra sites in relatively similar coastal graminoid-dominated vegetation types had a large variability from relatively strong sinks to large $CO_2$ sources with average annual NEE ranging from -37 to +60 g C m$^{-2}$ year$^{-1}$.

### 3. Geospatial data

Predictors included in our models and their main theoretical links to fluxes are listed in Supplementary Table 3.

#### 3.1 Processing geospatial data

To build a continuous dataset of gridded predictor variables, we had to gap-fill some of the predictor data (NDVI) and/or include some lower-level quality data (LST). We gap-filled and smoothed MODIS NDVI at their original temporal resolution using the method developed by Kong et al. (2019) [9]. This denoising method uses weighted Whittaker approach [10] with parameter λ across space for reconstructing gap-filled and smoothed remote sensing vegetation index time series and was efficiently employed in Google Earth Engine (see here https://github.com/gee-hydro/gee_whittaker_kong2019_validation). All the available NDVI data were used but were weighted to account for good and poor quality data (e.g., snow), and the algorithm was run three times to optimize performance. We used the most recent and efficient version of this algorithm with a constant parameter λ found here (https://code.earthengine.google.com/09fef6c8c16919f8ecaa455aae5362b0). The gap-filling and smoothing method with constant λ used here was originally developed for LAI. Therefore, the available scripts developed by Kong et al. (2019) had to be adjusted for NDVI. The λ parameter was manually optimized by comparing the effects of λ set to 10-700.

The GIMMS3g NDVI data includes gap-filled data provided by the data developers (flag 1: NDVI retrieved from spline interpolation, flag 2: NDVI retrieved from seasonal profile, possible snow/cloud). We used the NDVI values with a quality flag 2 if no other information was available (during the winter). During the summer, we used NDVI values with flag 1 to gap-fill data. Small remaining gaps were filled with linear interpolation.

We acknowledge the uncertainties associated with gap-filling NDVI data throughout the snowy shoulder and winter seasons but took this approach to assure that our predictors have data throughout the entire year, as most of the machine learning models cannot deal with missing data accurately. We justified this decision further by the fact that we are using these data to train multivariate models where, for example, climate data is likely the most important predictor instead of the gap-filled (and temporally not variable) optical remotely sensed data during the winter. We further supported this approach with the idea that vegetation biomass (and greenness) generally stays the same (or is lower) after the last good-quality autumn pixel value throughout the winter. We verified that the winter values were higher in highly productive ecosystems (e.g., forests, where winter NDVI values were close to 0.5) compared to sparse ecosystems (e.g., tundra, where winter NDVI values were close to 0-0.2) to make sure that the vegetation indices differ spatially in expected ways. We extracted the final predictor values at our flux sites from this gap-filled and smoothed data. These correlated significantly with the non-gap-filled values in the summer (Pearson correlation 0.88).

The geospatial data had differences in spatial resolution and terrestrial surface coverage (e.g., lake and ocean distribution) and ABCflux site coordinate accuracy was also variable. Therefore, in cases where some sites and/or monthly observations would have received NA values, we extracted the closest non-NA pixel values. This way we were able to keep all the sites from this sparsely measured region in the analysis.

##### 3.2 Additional predictors that were tested

We tested several other data sources as predictors for our models in addition to the ones listed on Supplementary Table 4. Those were dropped because (i) they were highly correlated with the other more powerful predictors that we already had (e.g., MOD13A2v006-based NDVI chosen over EVI or MOD11A2v006 surface temperature over TerraClimate-based air temperature for 1-km models; a Pearson correlation higher than 0.8 was considered as a cutoff value), (ii) they had data from a temporal period that was shorter than our study period (e.g. fractional open water cover 2002-2015 or ESA CCI annual permafrost layers 1998-2017 [11,12]), (iii) they had unrealistic values within our study domain (e.g., pixels indicating frozen status along the Swedish coastline in July [13], or (iv) they were missing a lot of data from the northernmost latitudes (e.g., northern Greenland) or coastlines.

Some potentially important variables were difficult to aggregate to ecologically meaningful predictors to be used in our monthly models. Describing fire history was one of those predictors as no accurate circumpolar burn history products exist that would extend beyond our study period that would allow us to accurately describe how, for example, a site/pixel that burned in 1970 is recovering after the fire. We tested including a ‘time since fire’ predictor based on MCD64A1v006 that goes up to 2020 [14] or GFED4 that goes up to 1997 [15] for each monthly observation to our machine learning model, but this variable was among the least important variables for all fluxes during the test runs (results not shown), likely due to its limitations in long-term data coverage. Additional predictors that were tested but excluded due to low importance included thermokarst coverage [16] and forest age in 2010 [17].

#### 4. Machine learning models

##### 4.1 Model structure

We had three response variables (GPP, $R_{eco}$, and NEE) and two different spatial resolutions and time periods of models. Consequently, we built a total of six models. We used the same predictors for all the response variables and a similar set of predictors for the 1-km and 8-km models to allow for straightforward model and prediction comparisons. For example, we used MODIS tree cover for the 1-km model and AVHRR tree cover for the 8-km model, and MODIS LST for the 1-km model and TerraClimate air temperature for the 8-km model.

We used random forest models which are a powerful machine learning model. They utilize several decision trees in an ensemble model framework and thus avoid overfitting, have high accuracy, are highly adaptable, and are not significantly impacted by outliers. Random forest models bootstrap the data several times and sample the predictor variables at each split during the tree building, after which the algorithm builds an ensemble prediction [18] . However, random forest models may suffer from overfitting and extrapolating outside the conditions present in the training data [19]. We tested other machine learning models (e.g., support vector machine, generalized boosted regression tree, generalized additive model, neural networks; results not shown). Random forest models outperformed those in terms of cross-validated predictive performance and produced the most realistic flux maps, which is why we only used random forest models.

For all the random forest models, we assumed Gaussian error distribution. Parameters for machine learning models were tuned separately for each response variable with the “caret” package in R [20,21] using the leave-one-site-out cross validation. We tuned the number of variables randomly sampled as candidates at each split from three options in each model. The best model with the final set of parameters was chosen based on the lowest root mean square error (RMSE) values. The only parameter that was tuned was the number of variables to randomly sample as candidates at each split, and it varied from 2 to 17 in the final models depending on the response variable.

We used partial dependence plots (i.e., response graphs) using the “pdp” package [22] and estimated variable importance of the predictors from each of the models using the “vip” package [23] (Section Machine learning models in the main text).The values on the y axis of each partial dependence plot can be interpreted as followed: yhat is conditional on other predictors in the model and their relationships with the predictor in the plot in question. Therefore, yhat values should not be directly compared with observed or predicted values, rather the patterns in yhat should be explored more generally. The x-axis represents the actual predictor values and can be used to infer, for example, conditions that lead to changes in yhat (tipping points), and the strength and direction of the relationship. Variable importance scores were estimated by randomly permuting the values of the predictor in the training data and exploring how this influenced model performance based on RMSE values, with the idea that random permutation would decrease model performance [18]. We used 100 simulations to calculate 100 importance scores which are shown in Supplementary Fig. 5-7.

#### 4.2 Model predictions

We used the random forest models to predict (i.e., upscale) fluxes with the 1-km model from 2001 to 2020 and 8-km model from 1990 to 2016. In total we produced 1692 upscaled flux maps. 8-km upscaled maps were further multiplied by the terrestrial surface cover within each 8-km pixel based on the 1-km ESA CCI+CAVM land cover dataset to remove fluxes from water bodies. These flux maps were robust across the two pixel resolutions, and a comparison of 2001-2016 average annual NEE maps showed that NEE was similar across the two pixel resolutions (Supplementary Fig. 20). The 1- and 8-km predictions had the largest differences in Siberia, as shown by the annual budget mismatches in Fig. 4, which also prevented us from merging the two predictions and calculating trends for the entire 1990-2020 period. We also compared upscaled NEE maps from two approaches: based on modeling NEE directly, and deriving it indirectly from the upscaled GPP and $R_{eco}$ maps. NEE from these two approaches yielded similar results, providing confidence in our results. Budgets estimates from the NEE and GPP-$R_{eco}$ approaches are also similar (Table 1, Supplementary Table 1, Supplementary Fig. 20, Supplementary Fig. 21-22). Overall, our upscaling results revealed a latitudinal pattern of average $CO_2$ fluxes, with stronger sinks in the south and weaker sinks or sources in the north (Fig. 1). However, the correlation between latitude and average NEE was moderate (Pearson's correlation for in-situ NEE: 0.26, $p=0.053$; for upscaled NEE: 0.55, $p<0.001$), suggesting that the latitudinal climate and radiation gradients were not the sole drivers of spatial $CO_2$ flux patterns.

##### 4.3 Model predictive performance and uncertainty

The predictive performance and uncertainty analysis was described in detail in the Machine learning modeling section of the Online methods. Here we provide a longer description of the strengths and limitations of our random forest models.

Overall, our models show good predictive performance. Compared to earlier ABZ upscaling efforts, our cross-validated performance metrics (Supplementary Figs. 1-3) indicate better performance. For example, the $R^2$ of our models ranged from 0.5 to 0.78, whereas Natali et al. (2019) [24] had an $R^2$ of 0.49 for winter NEE and Virkkala et al. (2021) [3] an $R^2$ of 0.07 for annual NEE, $R^2$ of 0.5 for annual GPP and annual $R_{eco}$; note though that the cross validation in Natali et al. 2019 was not based on a leave-one-site out approach. However, our performance metrics also indicate that strong sinks and sources, and high GPP and $R_{eco}$ were underestimated - a common issue in any kind of modeling [25]. As described with the mean bias error (MBE) metric, the models had a small tendency to overestimate fluxes (i.e., predict too small net sinks or too high net emissions), as reflected by the small and positive MBE values. Further, the distribution of NEE residuals was slightly skewed towards negative residuals (i.e., NEE overestimated; Supplementary Fig. 29). However, the majority of the observed and predicted values were close to the 1:1 line, and issues associated with the model underestimating strong net sinks were clearly larger (deviation up to -150 g C $m^{-2}$ $month^{-1}$) than the model underestimating strong net sources (deviation up to 80 g C $m^{-2}$ $month^{-1}$). For NEE, it was clear that situations where the modeled month differed significantly from the average monthly flux at the site were predicted worst (shown with yellow and light green values or dark blue values; Supplementary Figs. 1-3). It is thus possible that we are missing predictors that accurately describe conditions from such different months.

We also evaluated how differences in flux measurement method and the exclusion of disturbed sites impact model predictive performance (Supplementary Table 5). Overall, performance statistics were similar across the approaches.

We evaluated the uncertainty of predictions by creating 20 bootstrapped datasets (with replacement; same sample size as in the original model training data) and using those to develop 20 individual models and predictions. For these bootstrapped datasets and models, we did not include the categorical month and land cover datasets as predictors due to bootstrapping resulting in situations where a factor level was entirely missing from the model training data (e.g., for barren class that had little data) which prevented us from predicting fluxes across the entire domain (i.e., predictions to barren class not possible when the model had no information about it). Out of the 20 predictions, we calculated the standard deviation to represent prediction uncertainty. Similar to the predictive performance metrics (largest issues in our models related to predicting strong sinks), the uncertainty analysis also points towards highest uncertainties in areas with strong sinks, such as in northern Europe and southwestern Russia. However, when the uncertainty estimates were presented relative to the average flux, uncertainties were highest in tundra regions and parts of northern boreal Canada which generally have low in-situ flux data coverage. In some areas of these regions, our upscaling shows unrealistically high NEE values. For example, some sparsely vegetated or barren mountainous regions in northern Siberia (Kolyma mountains) or northern Europe (Scandes mountains) showed net emissions of 30-50 g C $m^{-2}$ $yr^{-1}$, which appears unrealistically high compared to the low vegetation carbon inputs and overall soil carbon pools. However, we did not mask the sparsely vegetated or barren areas away from our upscaling because we had data from these vegetation classes indicating that there is small but significant growing season and annual uptake in these regions [26,27]. Overall, the spatial uncertainty maps thus emphasize uncertainties both associated with model performance with strong sinks, and data gaps.

To further understand the uncertainties related to data gaps, we used a multivariate environmental dissimilarity surface analysis (MESS) to define the area of extrapolation in our models [28]. We used average annual environmental conditions over 2001-2020 of the 7 most important variables for this analysis (solar radiation, NDVI, land surface temperature, soil temperature, snow cover, soil organic carbon stock, soil pH, permafrost probability); average NDVI conditions were calculated for the June-August period alone. MESS represents how similar a point (i.e., a site) is to a reference set of points (i.e., all the ABZ conditions), with respect to a set of predictor variables. Negative values represent sites where at least one variable has a value that is outside the range of environments over the reference set (i.e., areas where the model extrapolates). The values in MESS are influenced by the full distribution of the reference points, so that points within the environmental range of the reference points but in relatively unusual environments will have a smaller value than those in common environments. Large positive values represent common conditions across the sites and ABZ. The analysis was done for the sites with data from January (primarily year-round sites). Our results show that 35% of the region was extrapolated (Supplementary Figure 4). If we limit our budget estimates to the area that was not extrapolated (i.e., 65% of the region), the annual NEE budget was -390 Tg C $yr^{-1}$.

Despite these uncertainties, our results show that machine learning-based upscaling is a promising approach for understanding recent trends in $CO_2$ fluxes as the models can easily integrate the most recent flux data and new predictor datasets while operating at high spatial and temporal resolutions. One uncertainty in upscaling remains how natural (e.g., thermokarst, insect outbreaks) and anthropogenic (e.g., forest management) disturbances are covered by the flux sites and explained with gridded data [29]. New predictors describing disturbances as well as supporting and extending the year-round flux network are critical to improve this upscaling and other synthesis and modeling efforts.

##### 4.4 Clustering of environmental rasters and trends

To understand what types of pixels are showing sinks or sources, or increasing sources or increasing sinks, we used clustering to visualize environmentally similar locations and their conditions. We included the most important predictors (solar radiation, June-August NDVI, LST, annual range in LST, soil temperature, snow cover, soil organic carbon stock, soil pH, and permafrost probability) averaged annually from 2001 to 2020 in clustering. A similar clustering analysis was done for the annual temporal trends of the predictors (solar radiation, June-August NDVI, LST, range in LST, soil temperature and snow cover). Trends were calculated with the nonparametric Mann–Kendall test using the “zyp” [30,31] package with pre-whitening (Zhang method [32]) to remove autocorrelation. All datasets were first re-scaled between 0 and 1 to ensure units are not the driving factor, after which we run a principal component analysis to condense data. Finally, we ran a k-means clustering analysis with the first four principal components using the Lloyd algorithm with 6 clusters, 10 chosen random sets and 500 maximum iterations; this analysis was done separately for the average environmental rasters and their temporal trends. Environmental conditions and upscaled fluxes were then visualized across the 6 clusters.

#### 5. Comparison with CMIP6 process models, inversions and earlier upscaling efforts

Supplementary Table 6 lists the CMIP6 process models and inversions included in our model intercomparison, and Supplementary Table 7 the budgets estimated with those. The models had variable inputs and structures, which causes differences in model outputs. We used an ensemble of these models (i.e. mean model output) for the two model categories (process and inversion models) in the main text because the uncertainty of the ensemble is expected to be lower than the uncertainty of a single model. A similar inversion ensemble was used in Liu et al. (2022) where the ensemble size and individual members were assessed and the temporal trends evaluated by fitting a linear mixed model to the ensemble, with the conclusion that the inversion ensemble remained robust for trend detection. Pixel-wise trends also correlated with EC data at the local scale [33]. Further, Treat and Virkkala et al. (2024) visualized the proportion of individual inversion pixels showing terrestrial net $CO_2$ sinks, sources, or neutrals [34]. They showed that most of the inversions suggest an annual $CO_2$ sink for large parts of the boreal despite some local disagreements, but areas in eastern and western parts of the continents were also net annual $CO_2$ sources. Most of the individual CMIP6 models suggested annual $CO_2$ sink or neutral values across the pixels.

##### 5.1. Inversions

The number of assimilated sites in the inversions in the Arctic-boreal region varies from ca. 10 up to 30 over the study period. At large scales, the inversions that have priors (i.e., prior values given by a process model; included in four out of five inversions) are hardly constrained by the priors but at regional scales and in areas with poor data coverage (e.g., Siberia), the posterior flux might be reflecting the prior flux. The average inversion NEE budget for the entire ABZ, including aquatic ecosystems while excluding fires, indicated a considerably stronger sink strength (-1054 Tg C $yr^{-1}$; range of average individual inversion estimates -790 to -1259 Tg C $yr^{-1}$).

We observed good agreement with our upscaling compared to inversion models. However, some disagreements were also apparent, especially in some parts of central and northern Siberia, where our upscaling suggested the region to be primarily a net annual $CO_2$ source and the inversion ensemble a $CO_2$ sink; however, the sparsity of year-round atmospheric or terrestrial flux data from this region prevents us from reliably concluding what the current sink status of the region is. Similarly, our upscaling showed sub-Arctic Canada in the Northwest Territories to have a large distribution of annual $CO_2$ sources whereas inversions suggested sinks. Overall, our combined NEE+fire estimates were on the higher end compared to inversions (i.e. weaker net $CO_2$ sinks or stronger net $CO_2$ emissions), especially in Canadian boreal, Siberian boreal, and Siberian tundra regions (Supplementary Fig. 17).

There are some similarities and differences in the long-term trends in our upscaling compared to inversions. Interannual variability in upscaled NEE and inversion-based NEE is highest in Siberia. However, inversions had more interannual variability in flux budgets overall compared to our upscaling. For example, NEE + fire budgets varied by 350 Tg C $yr^{-1}$ in our upscaling whereas those could range by 750 Tg C $yr^{-1}$ in inversion estimates. In our NEE upscaling, interannual variability in NEE was strongly related to air temperature. For example, in 2020 with a record-warm year in Siberia, the NEE budget changed from ca. -400 to -500 Tg C $yr^{-1}$ in Siberian boreal. It is possible that in 2020 Siberian ecosystems also experienced drought that should have decreased uptake as indicated by some of the inversions (Supplementary Fig. 17), which our models did not capture. However, during an extreme disturbance year in 2003 in Siberia with a high extent of fires and a decline in NDVI [36], our upscaling shows an increase in net $CO_2$ emissions with NEE changing from ca. -300 to -200 Tg C $yr^{-1}$ in the boreal biome and 25 to 40 Tg C $yr^{-1}$ in the tundra biome; this increase of net emissions was shown for most inversions in the Siberian boreal region as well. This provides confidence that our upscaling captures the impact of some of the extreme years that are increasingly important for ABZ $CO_2$ budgets. A visualization of the pixel-wise fluxes in two extreme years: the 2003 fire year and 2020 warm year are shown in Supplementary Fig. 23.

##### 5.2 CMIP6 process models

The ensemble mean of CMIP6 process models 29 showed consistently stronger tundra CO2 sink strength (-48 Tg C yr-1) than found in this study and weaker sink strength in the boreal zone (-391 Tg C yr-1) despite the mean NEE budget being very close to ours (-501 Tg C yr-1) (Fig. 1). However, variability in individual CMIP6 was high, with average NEE budgets for the ABZ ranging between -1256 to 963 Tg C yr-1 depending on the model.

##### 5.3 FLUXCOM-X-BASE

FLUXCOM-X-BASE-based estimates of annual net uptake budgets derived with global upscaling and gradient boosting trees [29,37,38] were twice as large compared to our upscaling over 2001-2020 (-1179 compared to -548 Tg C $yr^{-1}$), both in boreal (-1077 compared to -593 Tg C $yr^{-1}$) and tundra (-103 compared to 45 Tg C $yr^{-1}$) biomes. Furthermore, the range of pixel-wise NEE values was much broader in FLUXCOM-X-BASE, with upscaled average annual NEE varying from -749 to +242 g C $m^{-2}$ $yr^{-1}$ whereas our values ranged between -235 to +160 g C $m^{-2}$ $yr^{-1}$ (Supplementary Fig. 32). A comparison between the average annual NEE maps showed FLUXCOM-X-BASE to have both larger sinks and sources compared to our upscaling. However, the mean difference (-31 g C $m^{-2}$ $yr^{-1}$) suggested FLUXCOM-X-BASE predicted stronger net uptake or weaker net emissions compared to our upscaling. Differences in mean fluxes were particularly high in deciduous broadleaved and mixed forests, where FLUXCOM-X-BASE predicts stronger growing season uptake (Supplementary Fig. 33).

The differences between our upscaling and FLUXCOM-X-BASE stem from the differences in model training data and model structures. Specifically, there were 295 global eddy covariance sites and 40 Arctic-boreal sites with year-round NEE data in FLUXCOM-X-BASE whereas we had 56 year-round eddy covariance sites with NEE data. FLUXCOM-X-BASE had six tundra sites (five sites from Alaska, one in Russia) and our study 14 year-round tundra sites spanning Greenland, Norway, northern and sub-Arctic Alaska, and Russia. The number of site-months remained relatively similar across years in FLUXCOM-X-BASE model training data (132 to 252 site-months depending on the year, median 168), with a good coverage of site-months from the most recent 2015-2020 period (132 to 180, median 162). Our upscaling was based on a more temporally unevenly distributed number of monthly eddy covariance measurements (6 to 357, median 224), and there was less data from the 2015-2020 period (6 to 210, median 113); note that we also include chamber observations which were not compared here. This difference in the data distribution is likely due to our database compilation being done before the FLUXNET2015 ONEFlux data products that included more recent measurements.

The biggest difference between FLUXCOM-X-BASE and our upscaling is that their extreme gradient boosting algorithm was trained using all global sites, whereas our model was regionally trained, focusing on the Arctic-boreal region. Thus, eddy covariance sites outside the Arctic-boreal region are likely also influencing FLUXCOM-X-BASE's upscaled patterns and might be one of the reasons, in addition to the overall smaller number of Arctic sites included in model training, for the stronger sink strength suggested by FLUXCOM-X-BASE. Across the entire global FLUXCOM-X-BASE model training dataset, annual NEE estimates vary from -1889 to 1693 g C $m^{-2}$ $yr^{-1}$, i.e. the maximum net sink or source estimates in FLUXCOM-X-BASE model training data are more than three times higher than the ABZ estimates (see below).

Differences might also be related to the temporal resolution of the models, and the range of flux estimates included in model training data. FLUXCOM-X-BASE is based on hourly fluxes and our upscaling is based on monthly fluxes. Higher temporal resolution of the data used in FLUXCOM-X-BASE may increase the amount of noise in the data, whereas averaging over longer time resolutions likely smooths out extreme values. Indeed, when the hourly fluxes that were used in FLUXCOM-X-BASE model training were summed to annual cumulative NEE, we identified sites that had unrealistically high or low annual NEE estimates. Annual NEE estimates in the model training data within the ABZ in FLUXCOM-X-BASE varied between -650 to 638 g C $m^{-2}$ $yr^{-1}$ (mean -41 g C $m^{-2}$ $yr^{-1}$), whereas in our model training data, annual NEE ranged between -359 to 158 g C $m^{-2}$ $yr^{-1}$ (mean -45 g C $m^{-2}$ $yr^{-1}$), which is much more in line with annual NEE estimates published in scientific literature by site PIs (range -367 to 191, mean -44 $m^{-2}$ $yr^{-1}$; see Virkkala et al. 2021). For example, there were some years in FLUXCOM-X-BASE model training data when RU-Cok and RU-Che were estimated to have an annual NEE of around -400 g C $m^{-2}$ $yr^{-1}$, which is unrealistically strong uptake for tundra sites, whereas US-BZB and FI-Let had values close to 600 g C $m^{-2}$ $yr^{-1}$.

Additionally, the types of predictors and the spatial resolution in upscaling might explain some of the differences. Although many of the predictors are quite similar between the two efforts (MODIS-based LST and vegetation indices, or ERA5-based meteorology variables), a key difference between the upscaling frameworks is that FLUXCOM-X-BASE uses in-situ meteorological data to train the models instead of the data extracted from geospatial datasets. Further, the coarser resolution of FLUXCOM models might miss some of the spatial heterogeneity of the ABZ, such as the different post-disturbance recovery conditions.

#### 6. Aggregating results

We calculated in-situ cumulative average fluxes by first calculating mean fluxes across years at each site to avoid biasing the statistics by long-term sites. Annual fluxes were calculated for sites that had the full year of monthly flux estimates. We used the package “terra” [41] to derive zonal statistics (mean fluxes and budgets) across the key regions.

### Supplementary Tables and Figures

Supplementary Table 1. Average gross primary productivity (GPP), ecosystem respiration ($R_{eco}$), and net ecosystem exchange (NEE) fluxes and budgets over 2001-2020 across vegetation types. Uncertainties represent standard deviations across sites (for the in-situ data), or across bootstrapped upscaled estimates. Positive numbers for NEE indicate net $CO_2$ loss to the atmosphere and negative numbers indicate net $CO_2$ uptake by the ecosystem. Mismatches in the site-level versus upscaled $CO_2$ fluxes are likely related to sites being biased to certain regions and years while upscaled summaries should provide more representative regional estimates but are influenced by model performance. NAs occurred in situations when flux data was entirely non-existent, not partitioned to GPP and $R_{eco}$, or when statistics were based on a single site (not possible to calculate standard deviation).

| Class | In-situ average NEE g C $m^{-2}$ $yr^{-1}$ | In-situ average GPP g C $m^{-2}$ $yr^{-1}$ | In-situ average $R_{eco}$ g C $m^{-2}$ $yr^{-1}$ | Upscaled average NEE g C $m^{-2}$ $yr^{-1}$ | Upscaled average GPP g C $m^{-2}$ $yr^{-1}$ | Upscaled average $R_{eco}$ g C $m^{-2}$ $yr^{-1}$ | Average NEE budget Tg C $yr^{-1}$ | Average GPP budget Tg C $yr^{-1}$ | Average $R_{eco}$ budget Tg C $yr^{-1}$ |
|---|---|---|---|---|---|---|---|---|---|
| Barren and prostrate shrub | -74 (± 61) | NA | NA | -4 (± 8) | 537 (± 23) | 513 (± 17) | -4 (± 17) | 482 (± 18) | 461 (± 15) |
| Graminoid | 10 (± 28) | 272 (± 4) | 269 (± 0) | 9 (± 9) | 519 (± 31) | 525 (± 23) | 6 (± 9) | 346 (± 9) | 350 (± 6) |
| Shrub | 35 (± 37) | 244 (± 44) | 288 (± 81) | 23 (± 9) | 572 (± 35) | 598 (± 28) | 9 (± 6) | 215 (± 5) | 225 (± 2) |
| Sparse boreal vegetation | -33 (± 124) | 443 (± 229) | 442 (± 135) | 35 (± 6) | 669 (± 29) | 698 (± 27) | 50 (± 24) | 962 (± 20) | 1003 (± 12) |
| Tree cover, broadleaved, deciduous | -112 (± NA) | 1100 (± NA) | 988 (± NA) | -185 (± 23) | 1568 (± 64) | 1433 (± 47) | -90 (± 10) | 765 (± 11) | 699 (± 10) |
| Tree cover, needleleaved, deciduous | -17 (± 29) | NA | NA | -61 (± 18) | 931 (± 67) | 875 (± 52) | -148 (± 45) | 2239 (± 32) | 2105 (± 26) |

| | | | | | | | | | |
|---|---|---|---|---|---|---|---|---|---|
| Tree cover, needleleaved, evergreen | -36 (± 81) | 773 (± 423) | 734 (± 431) | -85 (± 12) | 1270 (± 47) | 1211 (± 34) | -222 (± 37) | 3309 (± 48) | 3153 (± 29) |
| Wetland | -22 (± 31) | 281 (± 78) | 256 (± 67) | -78 (± 11) | 778 (± 39) | 697 (± 28) | -47 (± 10) | 464 (± 9) | 415 (± 6) |
| Mosaic and mixed vegetation | -54 (± 74) | 697 (± 265) | 643 (± 223) | -117 (± 15) | 1358 (± 53) | 1274 (± 38) | -102 (± 14) | 1188 (± 16) | 1114 (± 12) |
| Alaskan boreal | -7 (± 60) | 592 (± 195) | 615 (± 148) | -12 (± 10) | 495 (± 41) | 486 (± 32) | -6 (± 4) | 228 (± 4) | 224 (± 3) |
| Alaskan tundra | 20 (± 31) | 277 (± 90) | 298 (± 113) | 5 (± 10) | 354 (± 28) | 360 (± 20) | 4 (± 7) | 270 (± 6) | 274 (± 3) |
| Canadian boreal | -39 (± 72) | 635 (± 255) | 598 (± 202) | -32 (± 6) | 557 (± 27) | 534 (± 20) | -129 (± 26) | 2214 (± 31) | 2125 (± 22) |
| Canadian tundra | NA | NA | NA | 1 (± 4) | 282 (± 13) | 278 (± 10) | 3 (± 20) | 644 (± 20) | 636 (± 16) |
| European boreal | -53 (± 94) | 837 (± 583) | 778 (± 614) | -64 (± 11) | 737 (± 40) | 684 (± 30) | -140 (± 20) | 1603 (± 23) | 1488 (± 16) |
| European tundra | -42 (± 52) | 440 (± 215) | 421 (± 232) | 14 (± 4) | 313 (± 15) | 328 (± 12) | 10 (± 5) | 225 (± 6) | 236 (± 4) |
| Siberian boreal | -105 (± 154) | 703 (± NA) | 572 (± NA) | -44 (± 9) | 535 (± 32) | 497 (± 23) | -319 (± 61) | 3875 (± 55) | 3599 (± 40) |

| | | | | | | | | | |
|---|---|---|---|---|---|---|---|---|---|
| Siberian tundra | -9 (± 30) | 241 (± 66) | 245 (± 92) | 9 (± 5) | 296 (± 18) | 307 (± 14) | 28 (± 24) | 910 (± 20) | 944 (± 12) |

Supplementary Table 2. The sites included in the analysis. For information about the sites see the Virkkala et al. (2021) [4] dataset.

| **Study ID Short** | **Site name** | **Site reference** | **Latitude** | **Longitude** | **Country** | **Flux method** |
|---|---|---|---|---|---|---|
| Adkinson_CA-WP2_tower1 | Alberta - Western Peatland - Poor Fen (Sphagnum moss) | CA-WP2 | 55.5375 | -112.334 | Canada | Eddy covariance |
| Adkinson_CA-WP3_tower2 | Alberta - Western Peatland - Rich Fen (Carex) | CA-WP3 | 54.47 | -113.32 | Canada | Eddy covariance |
| Alekseychik_RU-Murk_tower1 | Mukhrino field station | RU-Murk | 60.9 | 68.7 | Russia | Eddy covariance |
| Aurela_FI-Kaa_tower1 | Kaamanen | FI-Kaa | 69.14057 | 27.26985 | Finland | Eddy covariance |
| Aurela_FI-Ken_tower2 | Kenttarova | FI-Ken | 67.98723 | 24.24305 | Finland | Eddy covariance |
| Aurela_FI-SamFell_tower3 | Sammaltunturi fell | FI-SamFell | 67.9733 | 24.11565 | Finland | Eddy covariance |
| Aurela_FI-Sod_tower1 | Sodankyla | FI-Sod | 67.36239 | 26.63859 | Finland | Eddy covariance |
| Aurela_RU-Tks_tower1 | Tiksi | RU-Tks | 71.59427 | 128.8878 | Russia | Eddy covariance |
| Backstrand_StordalenMire_Ch | Stordalen Mire | Palsa Site,Sphagnum Site,Eriophorum Site | 68.36667 | 19.05 | Sweden | Chamber |
| BangYong_US-KOC_tower1 | US-KOC, Council | US-KOC, Council | 64.8439 | -163.711 | USA | Eddy covariance |
| Bergeron_CA-sOBS_tower1 | Southern Old Black Spruce | CA-sOBS | 53.99 | -105.12 | Canada | Eddy covariance |
| Bjoerkman_Adventdalen_Diff | Adventdalen, Svalbard | heath shallow,meadow shallow | 78.167 | 16.067 | Norway | Diffusion through snow |

| | | | | | | |
|---|---|---|---|---|---|---|
| Bjoerkman_Latnjajaure_Diff | Latnjajaure | heath snowbed,meadow snowbed,heath meadow,mesic meadow, heath shallow | 68.333 | 18.5 | Sweden | Diffusion through snow |
| Boike_NO-Blv_tower1 | Bayelva, Spitsbergen | NO-Blv | 78.9216 | 11.8311 | Norway | Eddy covariance |
| Bret-Harte_US-ICs_tower1 | Imnavait Creek Watershed | US-ICs | 68.6058 | -149.311 | USA | Eddy covariance |
| Bret-Harte_US-ICt_tower2 | Imnavait Creek Watershed | US-ICt | 68.6063 | -149.304 | USA | Eddy covariance |
| Cannone_Adventdalen1_Ch | Adventdalen | P1 | 78.18506 | 15.92633 | Norway | Chamber |
| Cannone_Adventdalen2_Ch | Adventdalen | P2 | 78.18511 | 15.92577 | Norway | Chamber |
| Cannone_Adventdalen3_Ch | Adventdalen | P3 | 78.18517 | 15.92551 | Norway | Chamber |
| Cannone_Adventdalen4_Ch | Adventdalen | P4 | 78.18529 | 15.92486 | Norway | Chamber |
| Cannone_Adventdalen5_Ch | Adventdalen | P5 | 78.18534 | 15.92644 | Norway | Chamber |
| Cannone_Adventdalen6_Ch | Adventdalen | P6 | 78.18539 | 15.92581 | Norway | Chamber |
| Cannone_Adventdalen7_Ch | Adventdalen | P7 | 78.18541 | 15.92515 | Norway | Chamber |
| Celis_EML_Ch | Eight Mile Lake | moist acidic tundra | 63.88306 | -149.226 | USA | Chamber |
| Chae_US-KOC_Ch | Council | US-KOC | 64.8439 | -163.711 | USA | Chamber |
| Christensen_NO-Adv_tower1 | Adventdalen | NO-Adv | 78.186 | 15.923 | Norway | Eddy covariance |
| Christiansen_DaringLake_Ch | Daring Lake | Low birch hummock | 64.833 | -111.633 | Canada | Chamber |
| Christiansen_DiskoIsland_Ch | Disko Island | Arctic Station | 69.254 | -53.514 | Greenland | Chamber |
| Christiansen_Zackenberg1_Ch | Zackenberg | dry heath | 74.467 | -20.577 | Greenland | Chamber |
| Christiansen_Zackenberg2_Ch | Zackenberg | Cassiope heath; NY-ITEX heath | 74.475 | -20.543 | Greenland | Chamber |
| Christiansen_Zackenberg3_Ch | Zackenberg | Salix heath; NY-ITEX heath | 74.475 | -20.54 | Greenland | Chamber |
| Davydov_Cherskiy1_Ch | Cherskiy | Larch-shrub forest, low density | 68.7 | 161.55 | Russia | Chamber |
| Davydov_Cherskiy2_Ch | Cherskiy | Post-fire shrub | 68.72 | 161.53 | Russia | Chamber |
| Davydov_Cherskiy3_Ch | Cherskiy | Old larch forest | 68.73 | 161.4 | Russia | Chamber |

| | | | | | | |
|---|---|---|---|---|---|---|
| Davydov_Cherskiy4_Ch | Cherskiy | Dense larch 'bamboo' stand | 68.75 | 161.45 | Russia | Chamber |
| Dolman_RU-Cok_tower1 | Chokurdakh | RU-Cok | 70.82914 | 147.4943 | Russia | Eddy covariance |
| Dolman_RU-Ypn_tower1 | Yakutsk | Larix cajanderii stand 160 yr old | 62.255 | 129.619 | Russia | Eddy covariance |
| Dyukarev_Siberia_Ch | Middle Taiga Zone | large hollow,small ridge | 60.9 | 68.7 | Russia | Chamber |
| Eckhardt_LRD_Ch | Lena River Delta | Wet tundra - polygon center,Dry tundra - polygon rim | 72.36667 | 126.4667 | Russia | Chamber |
| Egan/Risk_ImnavaitCreek_Ch | Imnavait Creek | heath | 68.607 | -149.296 | USA | Chamber |
| Elberling_Endalen_Ch | Endalen, Svalbard | Moist Cassiope heath,Dry Dryas heath,Salix snow bed | 78.2 | 15.6 | Norway | Chamber |
| Elberling_GL-Dsk_tower1 | Disko Island | GL-Dsk | 69.253 | -53.514 | Greenlan d | Eddy covariance |
| Emmerton_CA-LHazen1_tower1 | Lake Hazen, Ellesmere Island | CA-LHazen1 | 82.82255 | -71.3809 | Canada | Eddy covariance |
| Emmerton_CA-LHazen2_tower2 | Lake Hazen, Ellesmere Island | CA-LHazen2 | 81.83447 | -71.3846 | Canada | Eddy covariance |
| Euskirchen_RU-Eusk_cher1_tower1 | Chersky Tower 1 | RU-Eusk_cher1 | 68.51351 | 161.5312 | Russia | Eddy covariance |
| Euskirchen_RU-Eusk_cher2_tower2 | Chersky Tower 2 | RU-Eusk_cher2 | 68.69808 | 161.5388 | Russia | Eddy covariance |
| Euskirchen_US-TFBog_tower2 | Bonanza Creek Thermokarst Bog | US-BZB | 64.69555 | -148.321 | USA | Eddy covariance |
| Euskirchen_US-TFBS_tower1 | Bonanza Creek Rich Fen | US-BZF | 64.69635 | -148.324 | USA | Eddy covariance |
| Euskirchen_US-TFRF_tower3 | Bonanza Creek Rich Fen | US-BZF | 64.70373 | -148.313 | USA | Eddy covariance |
| Falk_Zackenberg_Ch | Zackenberg | | 74.5 | -20.5 | Greenlan d | Chamber |
| Friborg_Se-St1_tower1 | Stordalen grassland | Se-St1 | 68.35415 | 19.05033 | Sweden | Eddy covariance |
| Friborg_Seida_tower1 | Seida | Mixed tundra with upland tundra heath, peat plateau and wetlands | 67.05 | 62.93333 | Russia | Eddy covariance |
| Friborg_Svalbard_Ch | Svalbard | Björnedalen | 78.224 | 15.324 | Norway | Chamber |
| Gasovic_FI-Salm_tower1 | Salmisuo | FI-Salm | 62.7833 | 30.9333 | Finland | Eddy covariance |
| Goeckede_RU-Ch2_tower2 | Cherski reference | RU-Ch2 | 68.61689 | 161.3509 | Russia | Eddy covariance |

| | | | | | | |
|---|---|---|---|---|---|---|
| Goulden_CA-NS1_tower1 | UCI-1850 burn site | CA-NS1 | 55.87917 | -98.4839 | Canada | Eddy covariance |
| Goulden_CA-NS2_tower2 | UCI-1930 burn site | CA-NS2 | 55.90583 | -98.5247 | Canada | Eddy covariance |
| Goulden_CA-NS3_tower3 | UCI-1964 burn site | CA-NS3 | 55.91167 | -98.3822 | Canada | Eddy covariance |
| Goulden_CA-NS4_tower4 | UCI-1964 burn site wet | CA-NS4 | 55.91437 | -98.3806 | Canada | Eddy covariance |
| Goulden_CA-NS5_tower5 | UCI-1981 burn site | CA-NS5 | 55.86306 | -98.485 | Canada | Eddy covariance |
| Goulden_CA-NS6_tower6 | UCI-1989 burn site | CA-NS6 | 55.91667 | -98.9644 | Canada | Eddy covariance |
| Goulden_CA-NS7_tower7 | UCI-1998 burn site | CA-NS7 | 56.63583 | -99.9483 | Canada | Eddy covariance |
| Goulden_CA-Oas_tower1 | Saskatchewan - Western Boreal, Mature Aspen | CA-Oas | 53.62889 | -106.198 | Canada | Eddy covariance |
| Harazano_US-Cms_tower1 | Central Marsh | US-Cms | 71.32019 | -156.622 | USA | Eddy covariance |
| Helbig_CA-SCB_tower1 | Scotty Creek Bog | CA-SCB | 61.3089 | -121.298 | Canada | Eddy covariance |
| Helbig_CA-SCC_tower1 | Scotty Creek Landscape | CA-SCC | 61.3079 | -121.299 | Canada | Eddy covariance |
| Huemmrich_Utqia?vik_Ch | Utqia?vik | wet sedge tundra | 71.322 | -156.602 | USA | Chamber |
| Humphreys_CA-CB_tower1 | Cape Bounty | CA-CB | 74.915 | -109.574 | Canada | Eddy covariance |
| Iwata_US-Rpf_tower1 | Poker Flat Research Range: Succession from fire scar to deciduous forest | US-Rpf | 65.11983 | -147.512 | USA | Eddy covariance |
| Iwata_US-Uaf_tower1 | University of Alaska, Fairbanks | US-Uaf | 64.86627 | -147.856 | USA | Eddy covariance |
| Jarveoja_DegeroStormyr_Ch | Degerö Stormyr | oligotrophic minerogenic mire complex | 64.18333 | 19.55 | Sweden | Chamber |
| Kade_ImnavaitCreek1_Ch | Imnavait Creek | wet sedge | 68.606 | -149.311 | USA | Chamber |
| Kade_ImnavaitCreek2_Ch | Imnavait Creek | tussock | 68.606 | -149.304 | USA | Chamber |
| Kade_ImnavaitCreek3_Ch | Imnavait Creek | heath | 68.607 | -149.296 | USA | Chamber |
| Kim_Coldfoot1_Ch | Coldfoot | Young Black Spruce | 67.183 | -150.297 | USA | Chamber |
| Kim_Coldfoot2_Ch | Coldfoot | Young Black Spruce | 67.18 | -150.31 | USA | Chamber |
| Kim_Council_Ch | Council, AK | tundra sphagnum,tundra lichen,tundra tussock | 64.861 | -163.711 | USA | Chamber |

| | | | | | | |
|---|---|---|---|---|---|---|
| Kim_Fairbanks1_Ch | Fairbanks | Old Black Spruce | 65.644 | -147.471 | USA | Chamber |
| Kim_Fairbanks2_Diff | Fairbanks | black spruce forest | 64.867 | -147.85 | USA | Diffusion through snow |
| Kim_InteriorAlaska1_Ch | Interior Alaska | Gold Creek White Spruce | 67.74 | -149.76 | USA | Chamber |
| Kim_InteriorAlaska2_Ch | Interior Alaska | Lower Yukon Black Spruce | 65.84 | -149.65 | USA | Chamber |
| Kim_InteriorAlaska3_Ch | Interior Alaska | Upper Yukon Black Spruce | 66.08 | -150.17 | USA | Chamber |
| Kim_NorthSlope1_Ch | North Slope | Subalpine tundra | 68.175 | -149.441 | USA | Chamber |
| Kim_NorthSlope2_Ch | North Slope | Upland tundra | 68.905 | -148.876 | USA | Chamber |
| Kim_NorthSlope3_Ch | North Slope | Subalpine tundra | 68.18 | -149.44 | USA | Chamber |
| Kim_NorthSlope4_Ch | North Slope | Upland tundra | 68.9 | -148.88 | USA | Chamber |
| Kim_NorthSlope5_Ch | North Slope | Coastal tundra | 69.84 | -148.71 | USA | Chamber |
| Kim_SouthBrooksRange1_Ch | South Brooks Range | Tundra-boreal ecotone | 67.991 | -149.76 | USA | Chamber |
| Kim_SouthBrooksRange2_Ch | South Brooks Range | Tundra-boreal ecotone | 67.99 | -149.76 | USA | Chamber |
| Kljun_CA-Ojp_tower3 | Saskatchewan - Western Boreal, Mature Jack Pine | CA-Ojp | 53.91634 | -104.692 | Canada | Eddy covariance |
| Kljun_CA-sOBS_tower2 | Southern Old Black Spruce | CA-sOBS | 53.99 | -105.12 | Canada | Eddy covariance |
| Kolari_FI-Var_tower1 | Varrio | FI-Var | 67.7549 | 29.69014 | Finland | Eddy covariance |
| Kutzbach_RU-LRD1_tower1 | Samoylov Island | RU-Sam | 72.37398 | 126.4967 | Russia | Eddy covariance |
| Kutzbach_RU-Sam_tower1 | Samoylov Island | RU-Sam | 72.37398 | 126.4967 | Russia | Eddy covariance |
| Kutzbach_RU-Sam_tower2 | Samoylov Island | RU-Sam | 72.37037 | 126.4817 | Russia | Eddy covariance |
| Kutzbach_Samoylov_Tower_3_closedpath | Samoylov Island | RU-Sam | 72.37382 | 126.4958 | Russia | Eddy covariance |
| Kutzbach_Samoylov_Tower_3_openpath | Samoylov Island | RU-Sam | 72.37382 | 126.4958 | Russia | Eddy covariance |
| Kwon_US-BEO_tower2 | Barrow-Bes (Biocomplexity Experiment South tower) | US-BEO | 71.2809 | -156.597 | USA | Eddy covariance |
| Kwon_US-BES _tower1 | Barrow-Bes (Biocomplexity Experiment South tower) | US-BES | 71.281 | -156.6 | USA | Eddy covariance |
| López-Blanco_GL-NuF_tower1 | Kobbefjord | GL-NuF | 64.1382 | -51.3784 | Greenland | Eddy covariance |

| | | | | | | |
|---|---|---|---|---|---|---|
| López-Blanco_GL-ZaF_tower1 | Zackenberg | GL-ZaF | 74.48143 | -20.5545 | Greenland | Eddy covariance |
| Lafleur_CA-DL1_tower1 | Daring Lake | CA-DL1 | 64.8689 | -111.575 | Canada | Eddy covariance |
| Lafleur_CA-DL3_tower3 | Daring Lake | CA-DL3 | 64.8722 | -111.549 | Canada | Eddy covariance |
| Lafleur_CA-DL4_tower4 | Daring Lake | CA-DL4 | 64.8631 | -111.65 | Canada | Eddy covariance |
| Lafleur_CA-Iqa_tower1 | Iqaluit | CA-Iqa | 63.79025 | -68.5601 | Canada | Eddy covariance |
| Lafleur_CA-Pin_tower1 | Pond Inlet | CA-Pin | 72.69275 | -77.9576 | Canada | Eddy covariance |
| Larsen_Abisko1_Ch | Abisko | | 68.35 | 18.81667 | Sweden | Chamber |
| Larsen_Abisko2_Ch | Abisko | Abisko Scientific Research Station | 68.3 | 18.82 | Sweden | Chamber |
| Laurila_FI-Kns_tower1 | Kalevansuo | FI-Kns | 60.64683 | 24.35617 | Finland | Eddy covariance |
| Laurila_FI-Let_tower1 | Lettosuo | FI-Let | 60.64183 | 23.95952 | Finland | Eddy covariance |
| Laurila_FI-Lom_tower1 | Lompolojankka | FI-Lom | 67.99724 | 24.20918 | Finland | Eddy covariance |
| Leffler_YKD_Ch | Yukon-Kuskokwim Delta | Tutakoke River | 61.25 | -165.62 | USA | Chamber |
| Lindroth_SE-Fla_tower1 | Flakaliden | SE-Fla | 64.11278 | 19.45694 | Sweden | Eddy covariance |
| Lindroth_SE-Kno_tower1 | Knottåsen | SE-Kno | 60.99825 | 16.21728 | Sweden | Eddy covariance |
| Lindroth_SE-Nor_tower1 | Norunda | SE-Nor | 60.0865 | 17.4795 | Sweden | Eddy covariance |
| Lund_DK-ZaH_tower1 | Zackenberg | DK-Zah, Heath | 74.47328 | -20.5503 | Greenland | Eddy covariance |
| Maanavilja_Kaamanen_Ch | Kaamanen | | 69.13333 | 27.28333 | Finland | Chamber |
| Machimura_RU-Nel_tower1 | Nelegel | RU-Nel | 62.31583 | 129.4997 | Russia | Eddy covariance |
| Margolis_CA-Obs_tower1 | Saskatchewan - Western Boreal, Mature Black Spruce | CA-Obs | 53.98717 | -105.118 | Canada | Eddy covariance |
| Margolis_CA-Qfo_tower1 | Quebec - Eastern Boreal, Mature Black Spruce | CA-Qfo | 49.6925 | -74.3421 | Canada | Eddy covariance |
| Marushchak_Seida_Ch | Seida | Upland Tundra Heath,Dry Peatlands, Wetlands | 67.05 | 62.93333 | Russia | Chamber |
| Mastepanov_Zackenberg_Ch | Zackenberg | fen | 74.479 | -20.555 | Greenland | Chamber |
| Maximov_RU-Elg_tower1 | Elgeeii | RU-Elg | 60.016 | 133.824 | Russia | Eddy covariance |

| | | | | | | |
|---|---|---|---|---|---|---|
| Maximov_RU-SkP_tower1 | Yakutsk Spasskaya Pad larch | RU-SkP | 62.255 | 129.168 | Russia | Eddy covariance |
| McCaughey_CA-Gro_tower1 | Ontario - Groundhog River, Boreal Mixedwood Forest | CA-Gro | 48.2167 | -82.1556 | Canada | Eddy covariance |
| McCaughey_CA-Man_tower1 | Manitoba - Northern Old Black Spruce (former BOREAS Northern Study Area) | CA-Man | 55.87962 | -98.4808 | Canada | Eddy covariance |
| Merbold_RU-Che_tower1 | Cherskiy | RU-Che | 68.61304 | 161.3414 | Russia | Eddy covariance |
| Miyazaki_MO-UFRS_tower1 | Mongolia | MO-UFRS | 48.27333 | 106.8508 | Mongolia | Eddy covariance |
| Mkhabela_CA-OJP_tower4 | Saskatchewan - Western Boreal, Mature Jack Pine | CA-Ojp | 53.916 | -104.69 | Canada | Eddy covariance |
| Mkhabela_CA-SF1_tower1 | Saskatchewan - Western Boreal, forest burned in 1977 | CA-SF1 | 54.48503 | -105.818 | Canada | Eddy covariance |
| Mkhabela_CA-SF2_tower2 | Saskatchewan - Western Boreal, forest burned in 1989 | CA-SF2 | 54.25392 | -105.878 | Canada | Eddy covariance |
| Mkhabela_CA-SF3_tower3 | Saskatchewan - Western Boreal, forest burned in 1998 | CA-SF3 | 54.09156 | -106.005 | Canada | Eddy covariance |
| Mkhabela_HJP02_tower7 | HJP02 Jack Pine | CA-HJP02 | 53.15 | -104.1 | Canada | Eddy covariance |
| Mkhabela_HJP75_tower5 | HJP75 Jack Pine | CA-HJP75 | 53.15 | -104.1 | Canada | Eddy covariance |
| Mkhabela_HJP94_tower6 | HJP94 Jack Pine | CA-HJP94 | 53.15 | -104.117 | Canada | Eddy covariance |
| Morgner_Adventdalen_Ch | Adventdalen, Svalbard | heath control,meadow control | 78.167 | 16.067 | Norway | Chamber |
| Nakai_US-Prr_tower1 | Poker Flats | US-Prr | 65.12367 | -147.488 | USA | Eddy covariance |
| Nielsen_Abisko_Ch | Abisko | Wet NE-facing slope | 68.35 | 18.81667 | Sweden | Chamber |
| Nilsson_SE-Deg_tower1 | Degerö | SE-Deg | 64.18203 | 19.55654 | Sweden | Eddy covariance |
| Olivas10_Utqia?vik_Ch | Utqiavik North,Utqiavik South,Utqia?vik Central | North,South,Central | 71.32 | -156.62 | USA | Chamber |
| Olivas11_Utqia?vik_Ch | Utqia?vik | Vascular-dominated,Intermediate,Polygon Rim | 71.32 | -156.62 | USA | Chamber |
| Parmentier_NO-And_tower1 | Andøya | NO-And | 69.14278 | 16.02222 | Norway | Eddy covariance |
| Pirk_Adventdalen_Diff | Adventdalen, Svalbard | Advent-fen, active low center ice wedge polyons | 78.183 | 15.917 | Norway | Diffusion through snow |
| Pirk_Zackenberg_Ch | Zackenberg | fen | 74.5 | -21 | Greenland | Chamber |
| Pirk_Zackenberg_Diff | Zackenberg | fen | 74.5 | -21 | Greenland | Diffusion through snow |

| | | | | | | |
|---|---|---|---|---|---|---|
| Poyatos_Petsikko_Ch | Petsikko | Several hummocks and hollows | 69.35983 | 27.23136 | Finland | Chamber |
| Rebmann_RU-Zot_tower1 | Zotino | RU-Zot | 60.8008 | 89.3507 | Russia | Eddy covariance |
| Rocha_US-An1_tower1 | Anaktuvuk River Severe Burn | US-An1 | 68.99 | -150.28 | USA | Eddy covariance |
| Rocha_US-An2_tower2 | Anaktuvuk River Moderate Burn | US-An2 | 68.95 | -150.21 | USA | Eddy covariance |
| Rocha_US-An3_tower3 | Anaktuvuk River Unburned | US-An3 | 68.93 | -150.27 | USA | Eddy covariance |
| Schuur_EML_Ch | Eight Mile Lake | minimal thaw,moderate thaw,extensive thaw | 63.88306 | -149.226 | USA | Chamber |
| Schuur_US-EML_tower1 | Eight Mile Lake | US-EML | 63.8784 | -149.254 | USA | Eddy covariance |
| Semenchuk_Adventdalen_Ch | Adventdalen, Svalbard | dry heath | 78.167 | 16.067 | Norway | Chamber |
| Shaver_US-ICh_tower1 | Imnavait Creek Watershed | US-ICh | 68.6068 | -149.296 | USA | Eddy covariance |
| Sonnentag_CA-SMC_tower1 | Smith Creek | CA-SMC | 63.153 | -123.252 | Canada | Eddy covariance |
| Sonnentag_CA-TVC_tower1 | Trail Valley Creek | CA-TVC | 68.74617 | -133.502 | Canada | Eddy covariance |
| Startsev_Anzac_Ch | Mackenzie Valley, Anzac, Mid-Boreal - Peat Plateau,MacKenzie Valley, Anzac, Mid-Boreal - Upland | mid boreal - peat plateau,mid boreal - upland | 56.4 | -111.03 | Canada | Chamber |
| Startsev_FortSimpson_Ch | Mackenzie Valley, Fort Simpson, High Boreal - Peat Plateau,Mackenzie Valley, Fort Simpson, High Boreal - Upland | boreal forest - peat plateau ,boreal forest - upland | 61.63 | -121.4 | Canada | Chamber |
| Startsev_Inuvik_Ch | Mackenzie Valley, Inuvik, High Sub-Arctic - Peat Plateau,Mackenzie Valley, Inuvik, High Sub-Arctic - Upland | high subarctic - peat plateau,high subarctic - upland | 68.32 | -133.43 | Canada | Chamber |
| Startsev_NormanWells_Ch | Mackenzie Valley, Normal Wells, Low Sub-Arctic - Peat Plateau,Mackenzie Valley, Norman Wells, Low Sub-Arctic - Upland | low subarctic - peat plateau,low subarctic - upland | 65.21 | -127.01 | Canada | Chamber |
| Startsev_NormanWells_Ch | Mackenzie Valley, Norman Wells, Low Sub-Arctic - Upland | low subarctic - upland | 65.21 | -127.01 | Canada | Chamber |
| Strachan_CA-LLC_tower1 | Lac Le Caron (hereafter referred to as LLC) peatland, an ombrotrophic bog | CA-LLC | 52.29028 | -75.2542 | Canada | Eddy covariance |
| Strebel_Adventdalen_Ch | Adventdalen, Svalbard | | 78.167 | 16.1 | Norway | Chamber |
| Sullivan_AgashashokRiver_Diff | Agashashok River, Noatak National Preserve | NTL, treeline low density spruce,STL, treeline low density white | 67.48 | -162.2 | USA | Diffusion through snow |

| | | | | | | |
|---|---|---|---|---|---|---|
| | | spruce,SNE, white spruce forest,NSE, white spruce forest,NNE, white spruce forest,SSE, white spruce forest,TER, low density white spruce | | | | |
| Sullivan_ToolikLake1_Diff | Toolik Lake | tussock tundra | 68.62 | -149.605 | USA | Diffusion through snow |
| Sullivan_ToolikLake2_Diff | Toolik Lake | dry heath tundra | 68.622 | -149.598 | USA | Diffusion through snow |
| Sullivan_ToolikLake3_Diff | Toolik Lake | wet sedge tundra | 68.625 | -149.6 | USA | Diffusion through snow |
| Sullivan_ToolikLake4_Diff | Toolik Lake | riparian willow tundra | 68.626 | -149.596 | USA | Diffusion through snow |
| Sullivan_ToolikLake5_Diff | Toolik Lake | dwarf birch tundra | 68.632 | -149.573 | USA | Diffusion through snow |
| Syed_CA-WP1_tower1 | Alberta - Western Peatland - LaBiche River,Black Spruce/Larch Fen | CA-WP1 | 54.95384 | -112.467 | Canada | Eddy covariance |
| TornDengel_US-NGB_tower1 | NGEE Arctic Barrow | US-NGB | 71.28333 | -156.616 | USA | Eddy covariance |
| TornDengel_US-NGC_tower1 | NGEE Arctic Council | US-NGC | 64.85196 | -163.7 | USA | Eddy covariance |
| Tuittila_FI-Sii_tower1 | Siikaneva | FI-Sii | 61.83265 | 24.19285 | Finland | Eddy covariance |
| Uchida_Svalbard_Ch | E. Brogger Glacier | | 79 | 12 | Norway | Chamber |
| Ueyama_US-CR-Fire_tower1 | Cascaden Ridge Fire Scar | US-Fcr | 65.39678 | -149.121 | USA | Eddy covariance |
| Vesala_FI-Hyy_tower1 | Hyytiala | FI-Hyy | 61.84741 | 24.29477 | Finland | Eddy covariance |
| Voigt_Seida_Ch | Northeast Russia | bare peat,peat plateau,upland tundra | 67.05 | 62.91667 | Russia | Chamber |
| Voigt_Seida_Ch | Northeast Russia | bare peat | 67.05 | 62.91667 | Russia | Chamber |
| Waldrop_BonanzaCreek_Diff | Bonanza Creek | Sphagnum bog | 64.69 | -148.32 | USA | Diffusion through snow |
| Welp_US-Bn1_tower1 | Delta Junction | Populus tremuloides; understory: salix; Epilobium angustifolium and Festuca altaica | 63.90111 | -145.373 | USA | Eddy covariance |

| | | | | | | |
|---|---|---|---|---|---|---|
| Welp_US-Bn2_tower2 | Delta Junction | US-Bn2 | 63.88806 | -145.739 | USA | Eddy covariance |
| Wickland_BonanzaCreek_Ch | Bonanza Creek | permafrost plateau PP,thermokarst wetland TW | 64.41 | -148.19 | USA | Chamber |
| Zona_US-Atq_tower1 | Atqasuk | US-Atq | 70.4696 | -157.409 | USA | Eddy covariance |
| Zona_US-Brw_tower1 | Barrow Environmental Observatory (BEO) tower | US-Brw | 71.281 | -156.596 | USA | Eddy covariance |
| Zona_US-Brw_tower2 | Barrow-Bes (Biocomplexity Experiment South tower) | US-Brw | 71.281 | -156.596 | USA | Eddy covariance |
| Zona_US-Brw_tower3 | Barrow | US-Brw | 71.323 | -156.609 | USA | Eddy covariance |
| Zona_US-Ivo_tower1 | Ivotuk | US-Ivo | 68.4865 | -155.75 | USA | Eddy covariance |
| Zyryanov_RU_IG_tower1 | Igarka | RU-IG | 67.4812 | 86.43727 | Russia | Eddy covariance |
| Zyryanov_RU_Tura_tower1 | Tura; Nizhnyaya Tunguska River | RU-Tur | 64.20889 | 100.4636 | Russia | Eddy covariance |

Supplementary Table 3. Predictor details.

| **Data product and name** | **Spatial resolution** | **Temporal resolution and period: Static, Monthly (or higher), Annual** | **Quality flags** | **Model (1 km or 8 km)** | **Reference** | **Mechanism for driving the flux** |
|---|---|---|---|---|---|---|
| TerraClimate meteorological variables: air temperature , vapor pressure deficit, and solar radiation | 1/24°, ~4 km | Monthly 1/1958-> | - | 1 and 8 km | [42] | Air temperatures control enzymatic processes and thus GPP and $R_{eco}$ [43,44]. Vapor pressure deficit is linked to GPP: higher moisture levels increase GPP [45]. GPP is dependent on solar radiation (and in particular diffuse radiation) as a resource for photosynthesis [46]. |
| Day-time land surface | 1 km | Monthly from 2/2000-> | We used bit 0-1 value | 1 km | [47] | Surface temperatures are more tightly linked to vegetation and soil conditions |

| | | | | | | |
|---|---|---|---|---|---|---|
| temperature MOD11A2v006 | | | <=1: Pixel produced, unreliable or unquantifiable quality, recommend examination of more detailed QA | | | than air temperatures and control enzymatic processes and thus GPP and $R_{eco}$ [48] |
| ERA5 land soil moisture and temperature at 0-5 cm depth, snow cover | 0.1°, ~9 km | Monthly from 1/1950-> | - | 1 and 8 km | [49,50] | Soil moisture is an important resource for GPP and regulates $R_{eco}$ ; drier soils often have higher $R_{eco}$ than water-saturated soils [51]. Soil temperatures control soil respiration which can occur at temperatures lower than 0 C and can contribute to $R_{eco}$ up to 70% [24,52]. Snow cover reflects both the amount of snow, and timing of snowmelt and snowfall which are important drivers of not only winter but also growing season fluxes [53,54]. |
| Barrow atmospheric CO2 concentrations | Assuming one location represents the entire atmosphere | Monthly from 1/1973 | - | 1 and 8 km | [55] | Increasing $CO_2$ concentrations ($CO_2$ fertilization) accelerate GPP[56] |
| ESA CCI vegetation type + Circumpolar Arctic Vegetation Map (CAVM) vegetation type | 1 km (ESA CCI originally 300 m; CAVM 1 km) | Static | - | 1 and 8 km | Following [3] based on ESA CCI (2017) and [57] | Vegetation composition and structure are important drivers of $CO_2$ fluxes [58] and also act as a surrogate for many other environmental conditions (e.g., soil wetness, soil nutrients). Classes included in our vegetation type map are: barren and prostrate shrub tundra, graminoid tundra, shrub tundra, wetland, sparse boreal vegetation, needleleaved evergreen tree cover, broadleaved deciduous tree cover, needleleaved deciduous tree cover, mosaic and mixed vegetation type. |
| NDVI MOD13A1v006 | 500 m | Monthly from 2/2000-> | We used SummaryQA bit 0-1 value 0 together with value 1 with smaller weights | 1 km | [59]; gap-filled and smoothed with weighted Whittaker & constant lambda approach [9] | NDVI represents vegetation greenness and productivity patterns, and is a widely used vegetation index that is strongly correlated with GPP and partly also $R_{eco}$ and NEE [60,61]. |
| GIMMS3g NDVI | ca. 8 km | Monthly from 7/1981 to 12/2017 | We used data covering all the flags 0-2; poorer-quality data used only to gap-fill high-quality data | 8 km | [62] | See above. |
| MOD44B | 250 m | Annual from | - | 1 km | [63] | Vegetation cover is linked to the amount |

| | | | | | | |
|---|---|---|---|---|---|---|
| Percent Tree Cover, Percent Non-Tree Cover, Percent Non-Vegetated Cover | | 2000-> | | | | of green biomass and thus $CO_2$ fluxes [64] |
| AVHRR VCF5KYR Percent Tree Cover, Percent Non-Tree Cover, Percent Non-Vegetated Cover | Ca. 8 km | Annual from 1982 to 2016 | - | 8 km | [65] | See above. |
| SoilGrids v2 variables: pH (water solution) at the topsoil (0-5 cm), soil organic carbon stock in the uppermost 2 m | 250 m | Static | - | 1 and 8 km | [66,67] | Soil pH may be associated with soil nutrient content and thus regulates the availability of resources for plants and microbes (lower pH potentially correlates with stronger net $CO_2$ sinks [68]). Soil organic carbon stock describes the amount of material available for decomposition and may thus be correlated with $R_{eco}$ [69]. |
| Topographic indices calculated from MERIT DEM: compound topographic index (CTI) | 250 m | Static | - | 1 and 8 km | [70] | CTI is a topographic index that describes the accumulation of water in topographic depressions (synonym to topographic wetness index), and might thus be correlated with GPP and $R_{eco}$ [71]. |
| Permafrost probability | 1 km | Static | - | 1 and 8 km | [72] | Permafrost protects organic matter from decomposition and thus defines how much material is available for decomposition in the soil [73]. |

Supplementary Table 4.

| | GPP | Reco | NEE |
|---|---|---|---|
| Sample size for 1 km model | 3869 | 3869 | 4765 |
| Sample size for 8 km model | 3968 | 3970 | 4897 |

Supplementary Table 5. Model performance based on different subsets of data for the 1-km models.

| Flux | Model training data | Performance estimates |
|---|---|---|
| NEE | All data | RMSE 15.9<br>R2 0.66<br>MAE 12.6 |
| NEE | Eddy covariance only | RMSE 17.5<br>R2 0.69<br>MAE 13.7 |
| NEE | Non-disturbed sites only | RMSE 14.6<br>R2 0.66<br>MAE 11.5 |
| GPP | All data | RMSE 35.0<br>R2 0.82<br>MAE 27.2 |
| GPP | Eddy covariance only | RMSE 32.3<br>R2 0.87<br>MAE 24.4 |
| GPP | Non-disturbed sites only | RMSE 35.2<br>R2 0.79<br>MAE 28.0 |
| Reco | All data | RMSE 28.9<br>R2 0.74<br>MAE 23.4 |
| Reco | Eddy covariance only | RMSE 27.5<br>R2 0.76<br>MAE 21.5 |
| Reco | Non-disturbed sites only | RMSE 28.0<br>R2 0.70<br>MAE 23.2 |

Supplementary Table 6. Details related to the process and inversion models included in the model intercomparison.

| Model type | Model name | Details and reference |
|---|---|---|
| Atmospheric inversions | CAMS | Release v21r1 of the inversion produced by the Copernicus Atmosphere Monitoring Service (https://atmosphere.copernicus.eu/), |

| | | |
|---|---|---|
| | | driven by air-sample measurements and included in the Global Carbon Budget 2022; total land CO2 flux adjusted for fossil fuel emissions, cement carbonation sink, and lateral fluxes[74]. |
| Atmospheric inversions | sEXTocNEET | Contribution to the Global Carbon Budget 2022; total land CO2 flux adjusted for fossil fuel emissions, cement carbonation sink, and lateral fluxes[74] |
| Atmospheric inversions | CTE | Contribution to the Global Carbon Budget 2022; driven by atmospheric observations in Obspack Globalviewplus v7.0 and NRT v7.2 [75]; total land CO2 flux adjusted for fossil fuel emissions, cement carbonation sink, and lateral fluxes[74] |
| Atmospheric inversions | NISMON | Contribution to the Global Carbon Budget 2022; total land CO2 flux adjusted for fossil fuel emissions, cement carbonation sink, and lateral fluxes[74] |
| Atmospheric inversions | UoE | Contribution to the Global Carbon Budget 2022; total land CO2 flux adjusted for fossil fuel emissions, cement carbonation sink, and lateral fluxes[74] |
| Process models: coupled CMIP6 models | ACCESS-ESM1-5 | Based on historical model runs with model outputs from 2001 to 2014 [35] |
| Process models: coupled CMIP6 models | BCC-ESM1 | Based on historical model runs with model outputs from 2001 to 2014 [35] |
| Process models: coupled CMIP6 models | BCC-CSM2-MR | Based on historical model runs with model outputs from 2001 to 2014 [35] |
| Process models: coupled CMIP6 models | CanESM5 | Based on historical model runs with model outputs from 2001 to 2014 [35] |

| | | |
|---|---|---|
| Process models: coupled CMIP6 models | CESM2 | Based on historical model runs with model outputs from 2001 to 2014 [35]; CESM2 includes permafrost carbon in the model |
| Process models: coupled CMIP6 models | CMCC-ESM2 | Based on historical model runs with model outputs from 2001 to 2014 [35] |
| Process models: coupled CMIP6 models | CNRM-ESM2 | Based on historical model runs with model outputs from 2001 to 2014 [35] |
| Process models: coupled CMIP6 models | GFDL-ESM4 | Based on historical model runs with model outputs from 2001 to 2014 [35] |
| Process models: coupled CMIP6 models | IPSL-CM6A | Based on historical model runs with model outputs from 2001 to 2014 [35] |
| Process models: coupled CMIP6 models | MIROC-ES2L | Based on historical model runs with model outputs from 2001 to 2014 [35] |
| Process models: coupled CMIP6 models | MPI-ESM1-2-LR | Based on historical model runs with model outputs from 2001 to 2014 [35] |
| Process models: coupled CMIP6 models | NorESM2-LM | Based on historical model runs with model outputs from 2001 to 2014 [35]; NorESM2-LM includes permafrost carbon in the model |
| Process models: coupled CMIP6 models | UKESM1-0-LL | Based on historical model runs with model outputs from 2001 to 2014 [35] |

Supplementary Table 7. Average annual regional budgets of individual inversions and CMIP6 models. BCC models were shown to produce outlier budgets in another high-latitude study as well [76].

| Model type | Model | Arctic-Boreal Zone | Tundra average regional | Boreal average regional | Permafrost region average |
|---|---|---|---|---|---|

| | | average regional budget NEE Tg C $yr^{-1}$ | budget NEE Tg C $yr^{-1}$ | budget NEE Tg C $yr^{-1}$ | regional budget NEE Tg C $yr^{-1}$ |
|---|---|---|---|---|---|
| This study | | -548 | 45 | -593 | -249 |
| Global upscaling | FLUXCOM-X-BASE | -1179 | -103 | -1077 | -621 |
| Inversion | CAMS | -1259 | -106 | -1061 | -784 |
| | CTE2021 | -1045 | -206 | -776 | -815 |
| | NISMON | -1245 | 13 | -1178 | -576 |
| | UoE | -790 | -34 | -683 | -488 |
| | sEXTocNEET | -930 | -6 | -841 | -614 |
| CMIP6 | ACCESS-ESM1-5 | -594 | -50 | -492 | -349 |
| | BCC-CSM2-MR | 963 | 171 | 731 | 690 |
| | BCC-ESM1 | 236 | 116 | 117 | 259 |
| | CanESM5 | -278 | -19 | -239 | -137 |
| | CESM2 | -624 | -38 | -515 | -292 |
| | CMCC-ESM2 | -440 | -14 | -367 | -171 |
| | CNRM-ESM2 | -1256 | -186 | -933 | -789 |
| | GFDL-ESM4 | -864 | -178 | -572 | -571 |
| | IPSL | -814 | -94 | -620 | -448 |

| | | | | | |
|---|---|---|---|---|---|
| | MIROC-ES2L | -835 | -122 | -621 | -531 |
| | MPI-ESM1 | -694 | -62 | -561 | -398 |
| | NorESM2-LM | -489 | -1 | -422 | -177 |
| | UKESM1 | -826 | -153 | -588 | -578 |

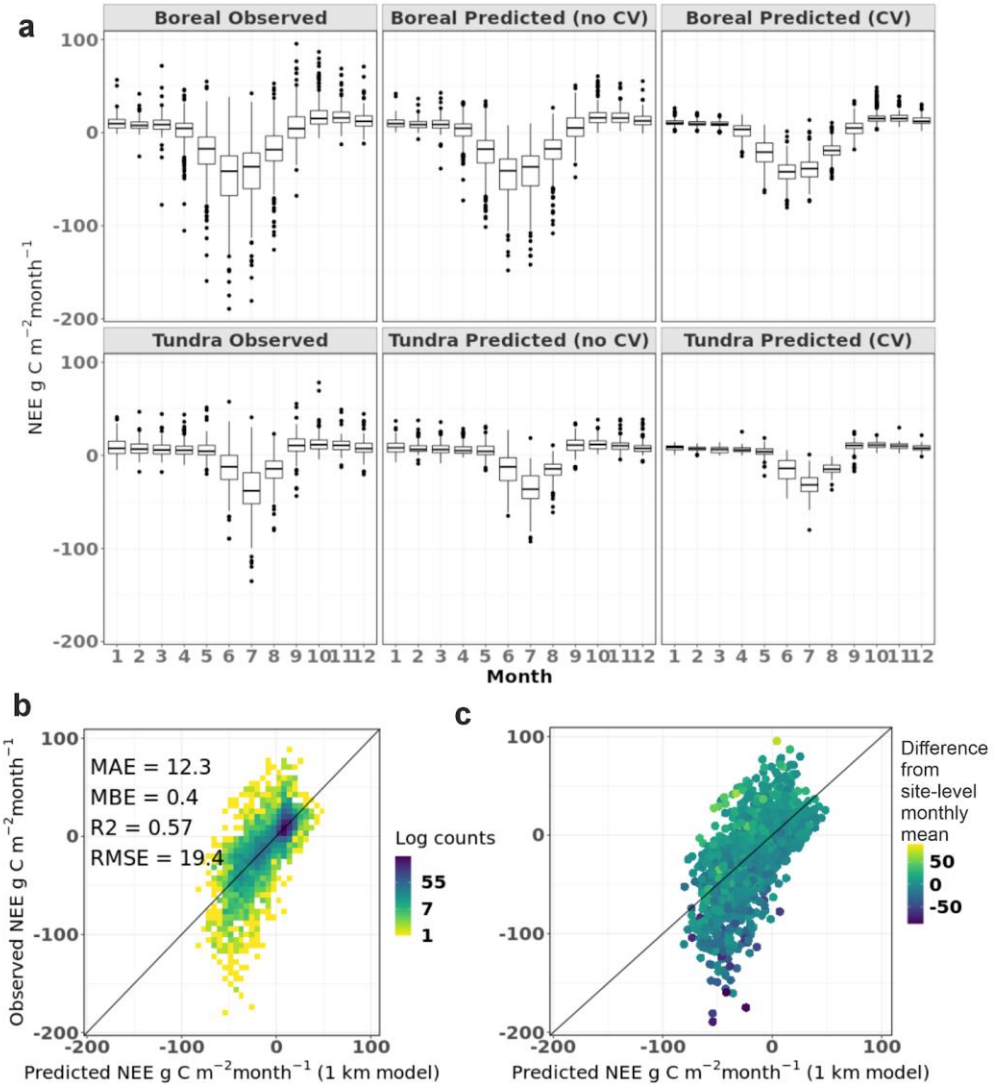

Supplementary Fig. 1. Predictive performance of the NEE model estimated using leave-one-site-out approach. Colors in subplot c indicate deviance from average site-level monthly flux and indicate that the model struggles the most when observations from individual sites have a large deviance from the mean.

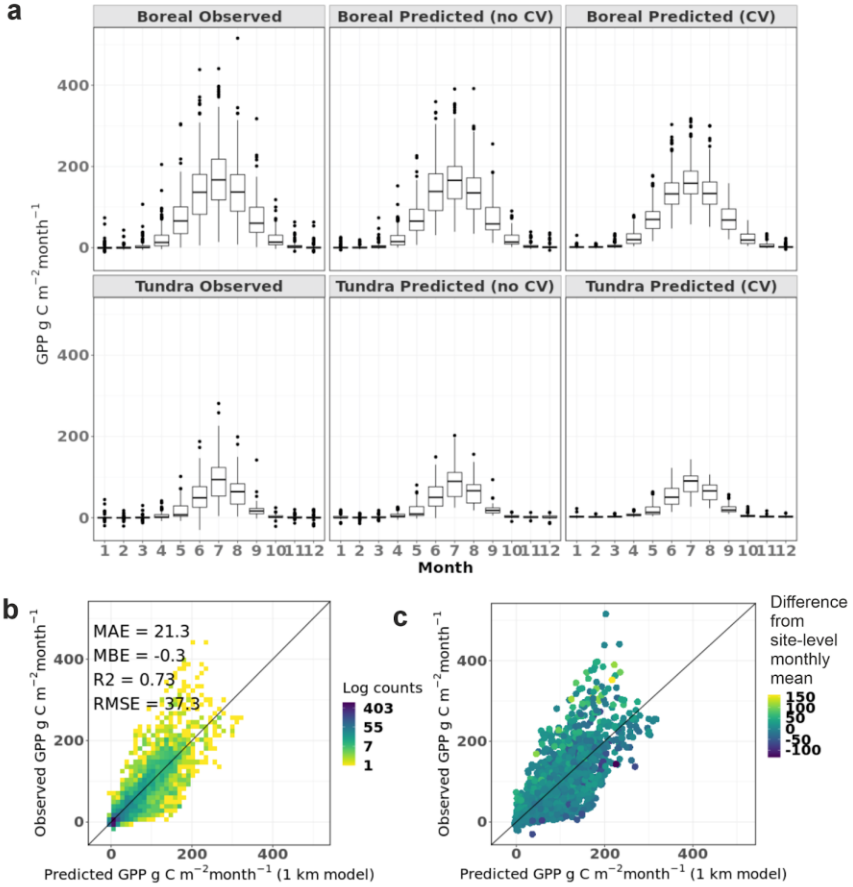

Supplementary Fig. 2. Predictive performance of the GPP model estimated using leave-one-site-out approach.

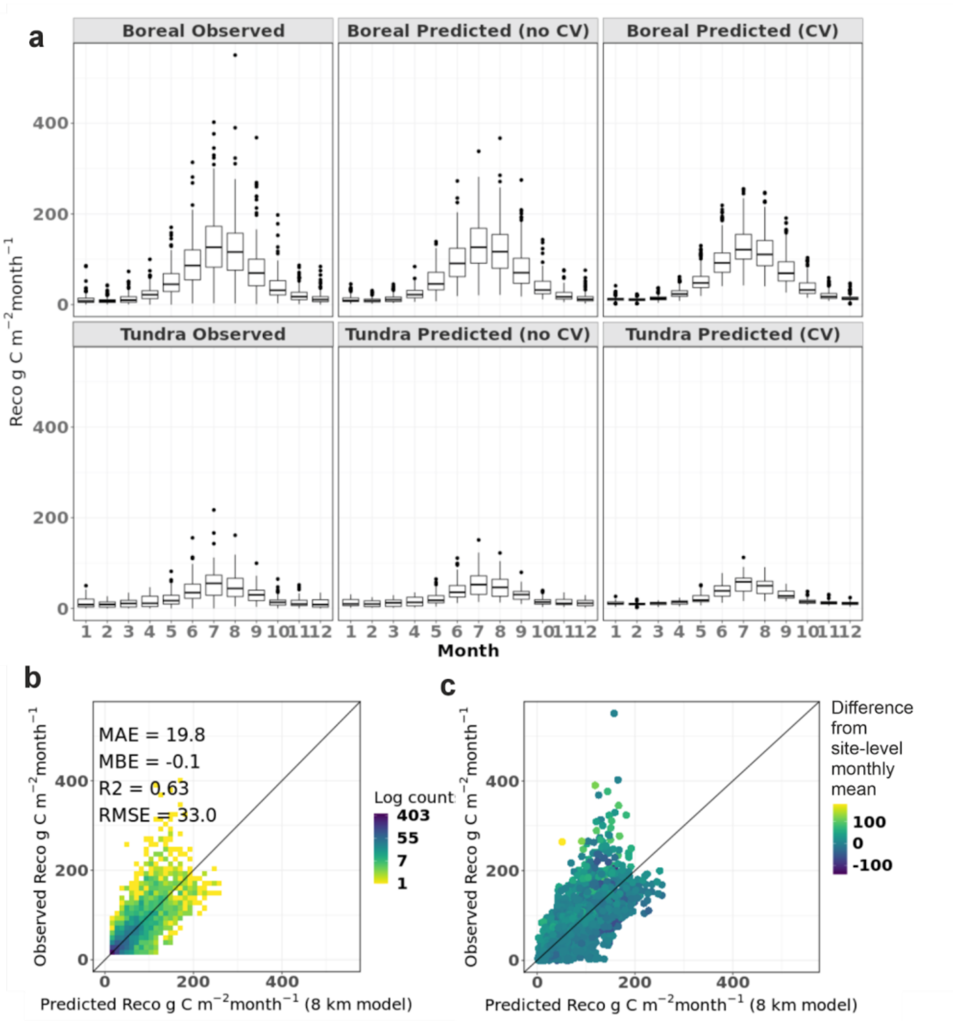

Supplementary Fig. 3. Predictive performance of the $R_{eco}$ model estimated using leave-one-site-out approach.

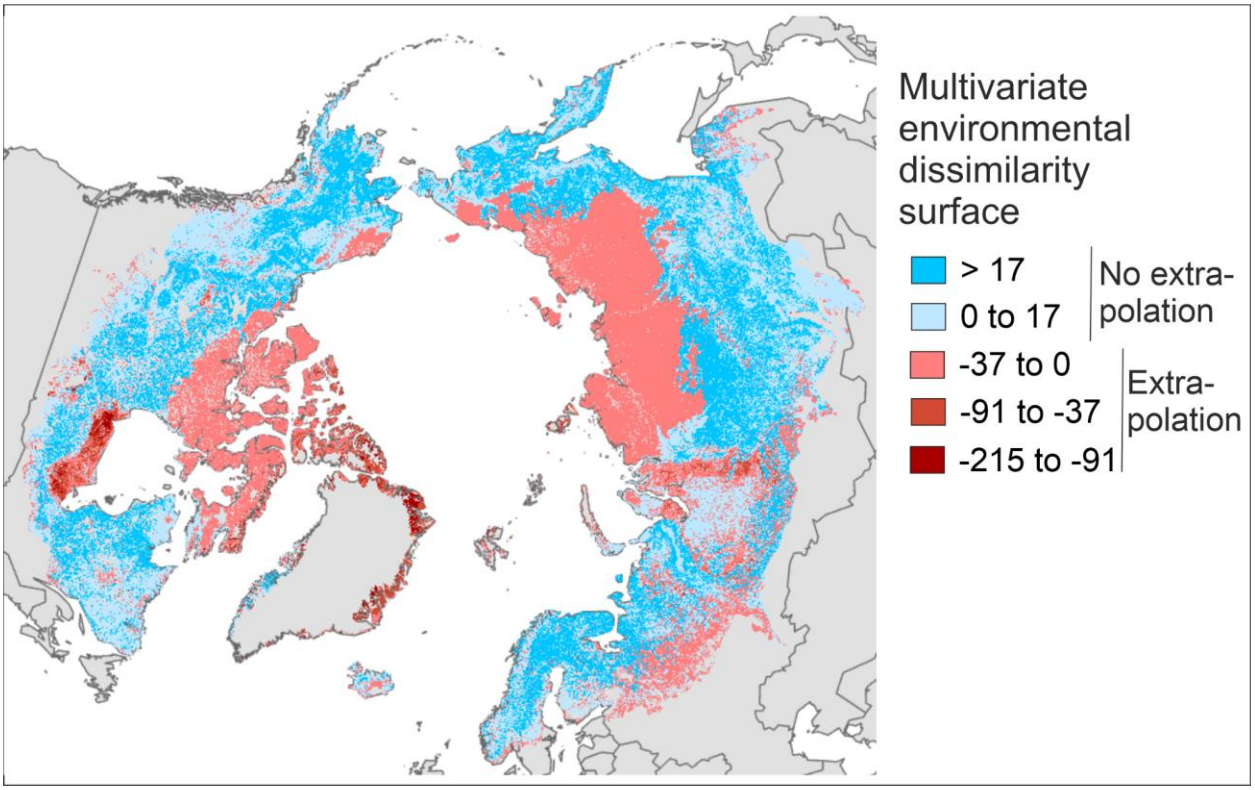

Supplementary Fig. 4. Maps showing the area of extrapolation for NEE models based on sites that have data at least from one January (i.e., year-round sites).

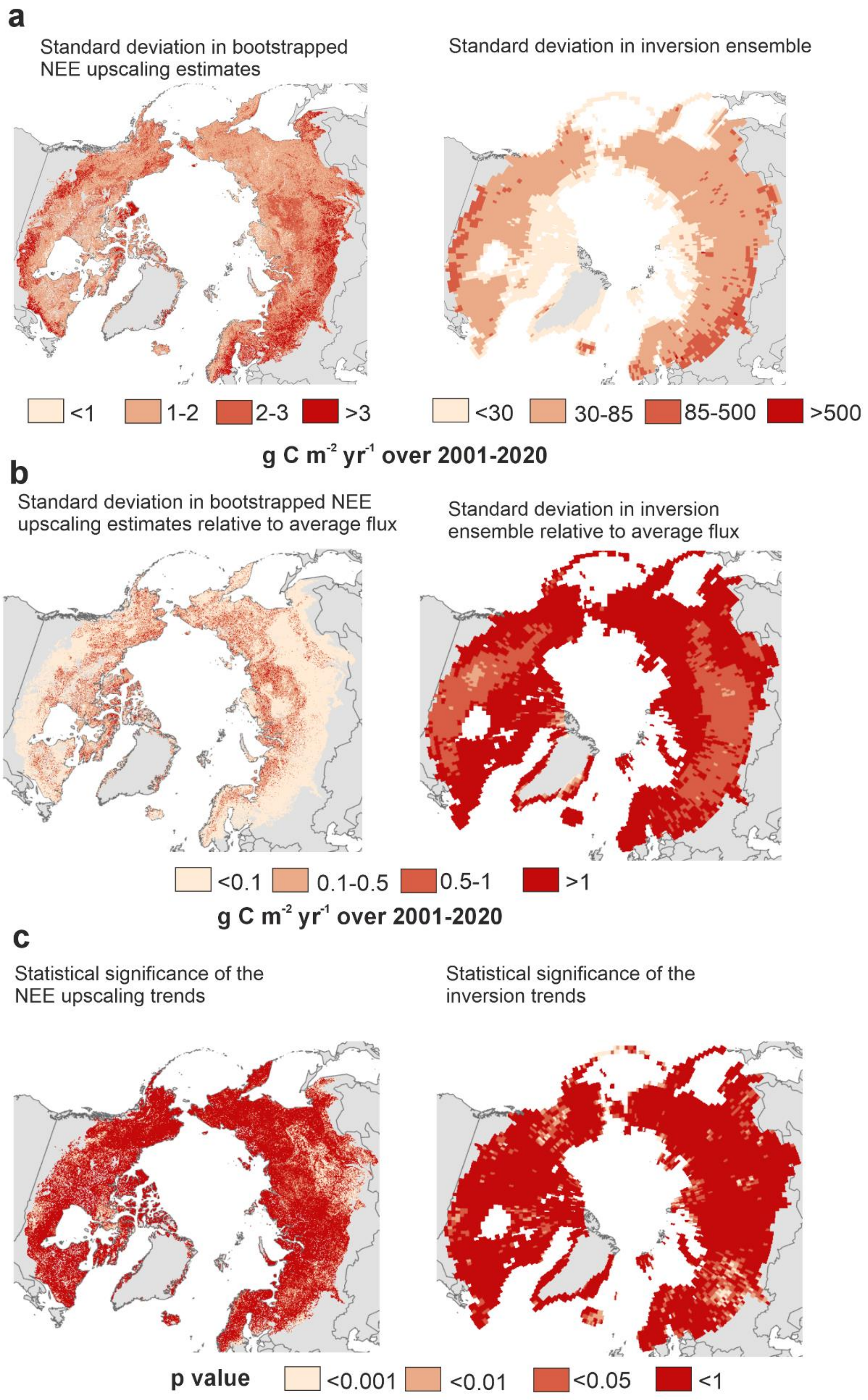

Supplementary Fig. 5. Uncertainties for the upscaled and inversion NEE.

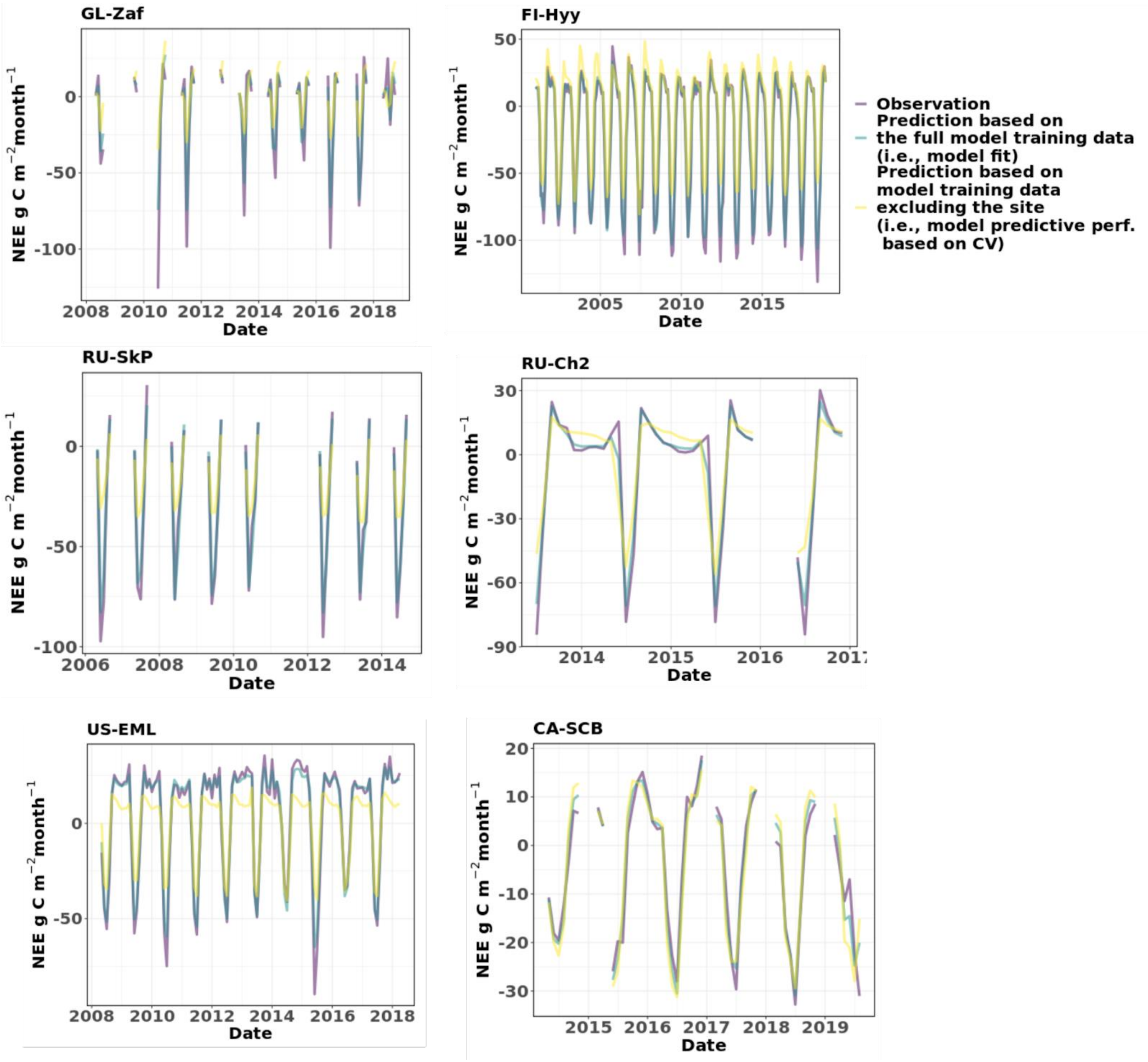

Supplementary Fig. 6. Time series of NEE from a subset of sites and their agreement with model predictions. Model fit indicates how well the model trained with the entire model training data predicts to the same data and model predictive performance shows how the models perform when a dataset excluding the specific site is used to train the model.

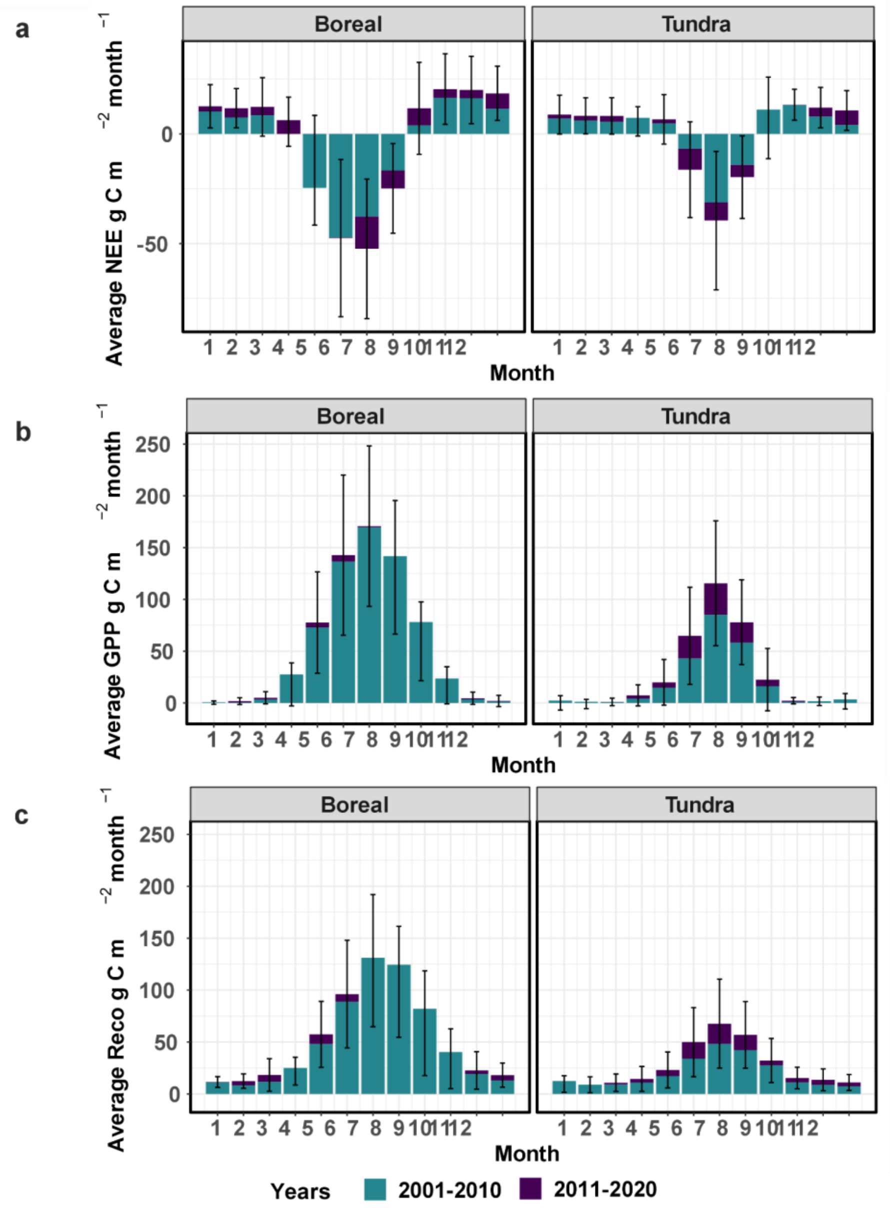

Supplementary Fig 7. Average in-situ monthly NEE, GPP, and $R_{eco}$ in boreal and tundra biomes during the past two decades. Note that this figure is highly uncertain as it does not account for the differences in the site distribution across the two decades. Fig. 3 in the main text shows the upscaled monthly fluxes that should better represent the average fluxes across the entire ABZ.

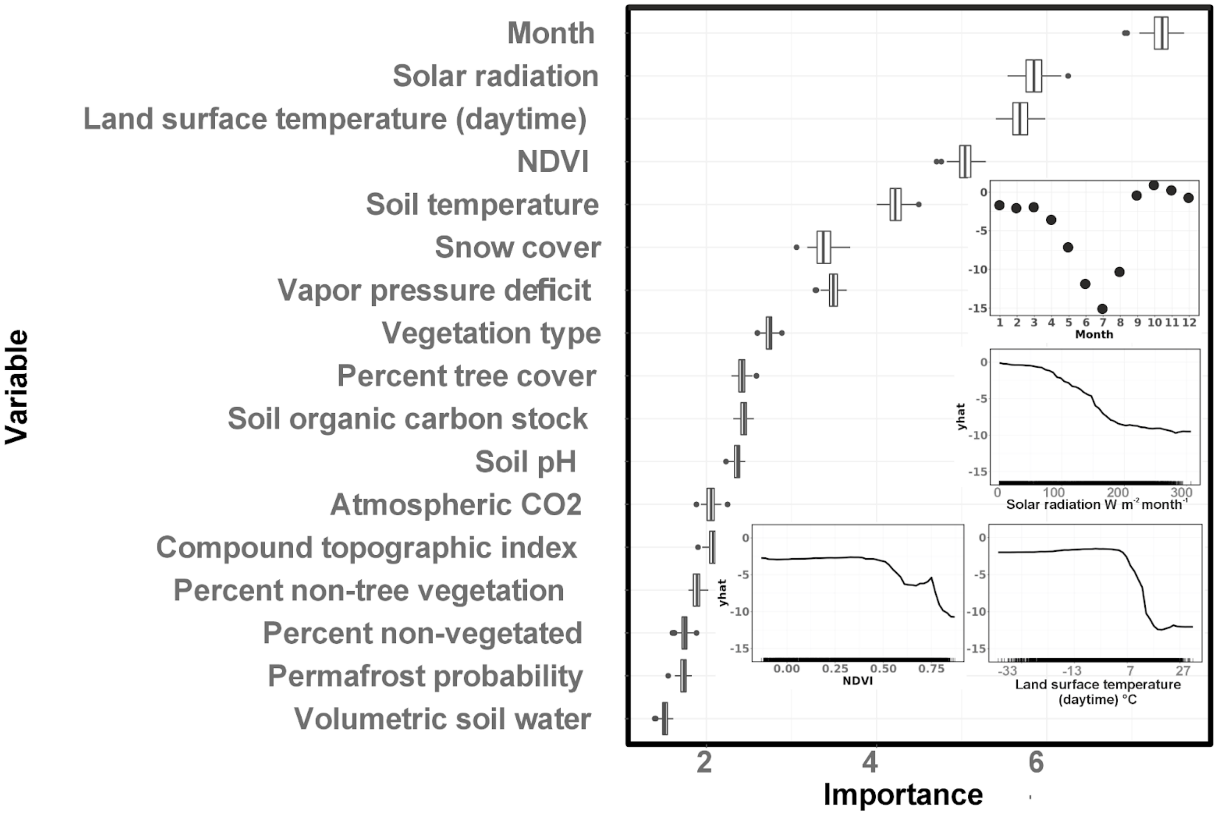

Supplementary Fig. 8. Variable importance plots and the partial dependence plots for the most important predictors of the 1-km NEE model.

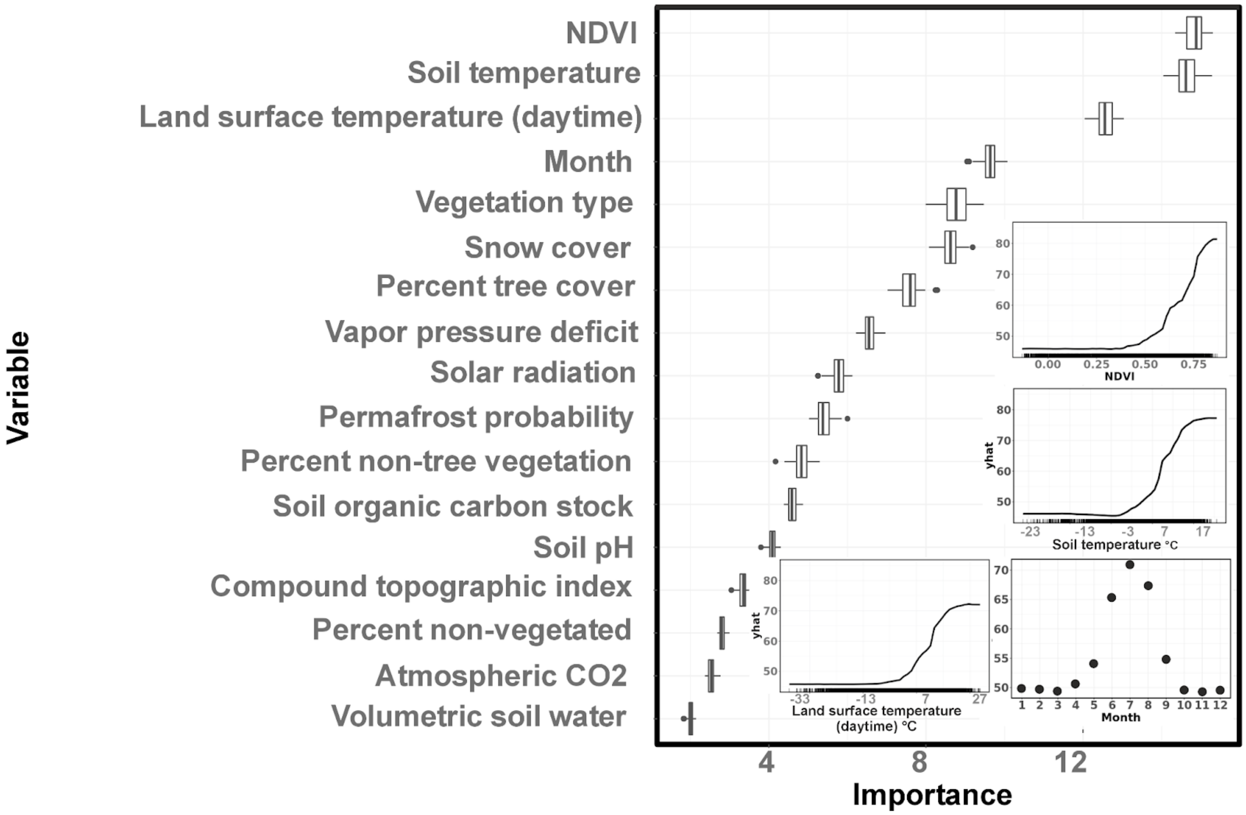

Supplementary Fig. 9. Variable importance plots and the partial dependence plots for the most important predictors of the GPP model.

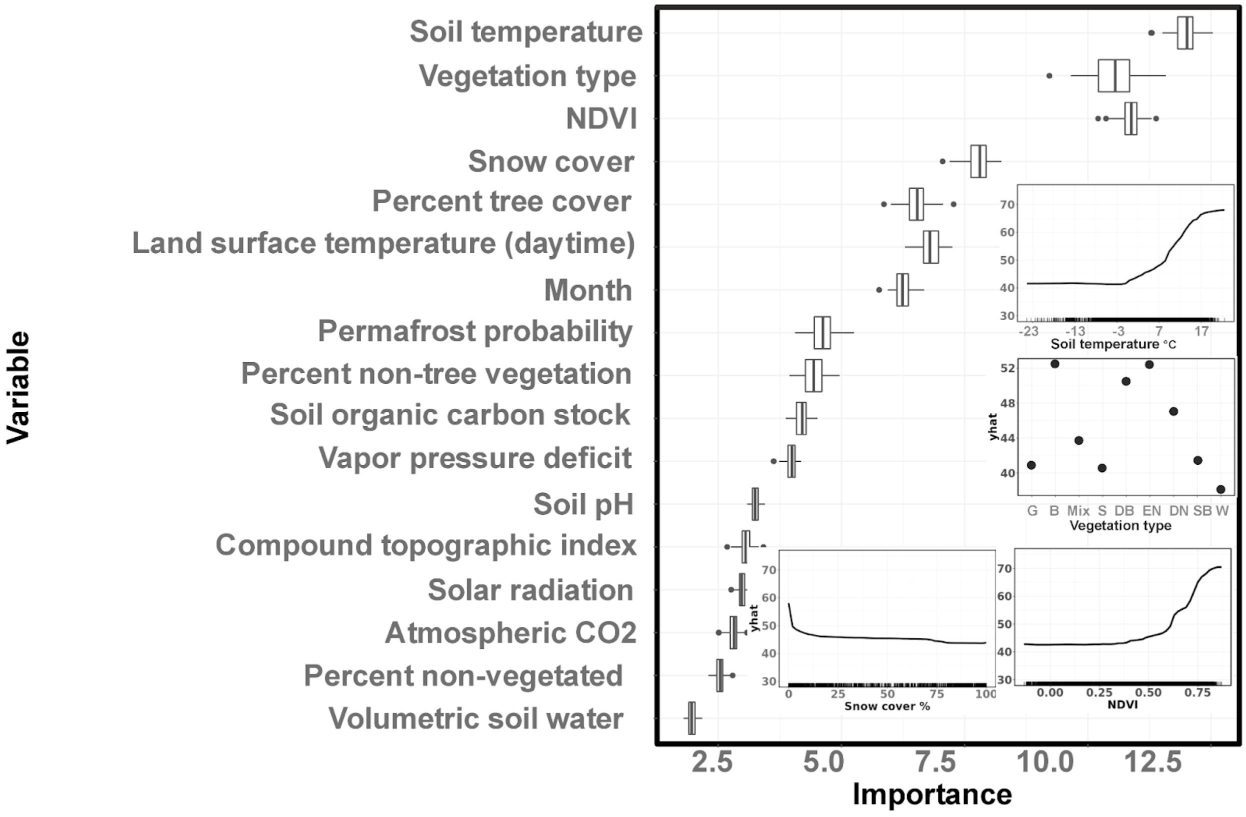

Supplementary Fig. 10. Variable importance plots and the partial dependence plots for the most important predictors of the $R_{eco}$ model. Vegetation types include G=graminoid, B=barren, Mix=mixed forest and mosaic vegetation type, S=shrub, DB=deciduous broadleaf forest, EN=evergreen needleleaf forest, DN=deciduous needleleaf forest, SB=sparse boreal vegetation, W=wetland.

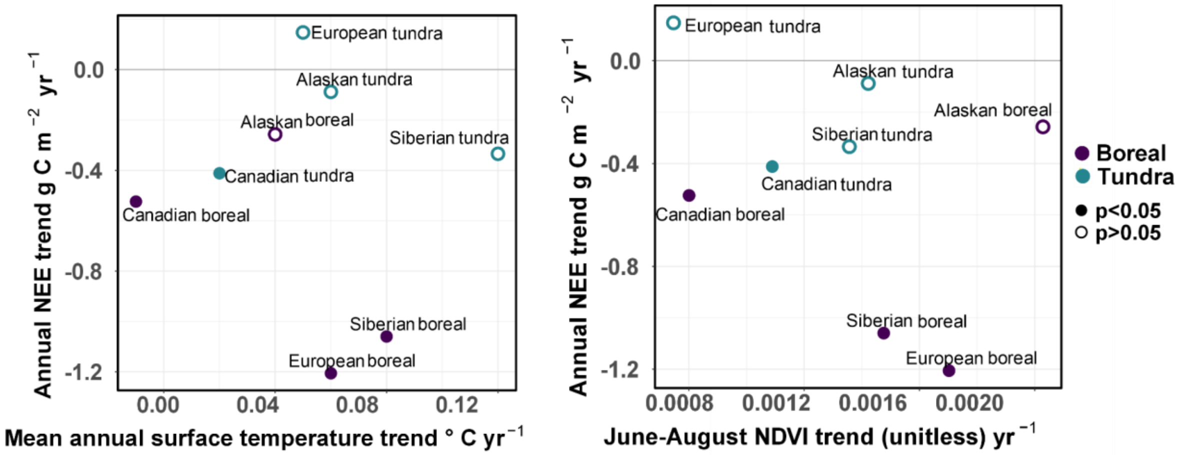

Supplementary Fig. 11. Correlation between average temperature and NDVI trends with upscaled average annual NEE trends over 2001-2020. The statistical significance of the NEE trend is shown with full and empty circles.

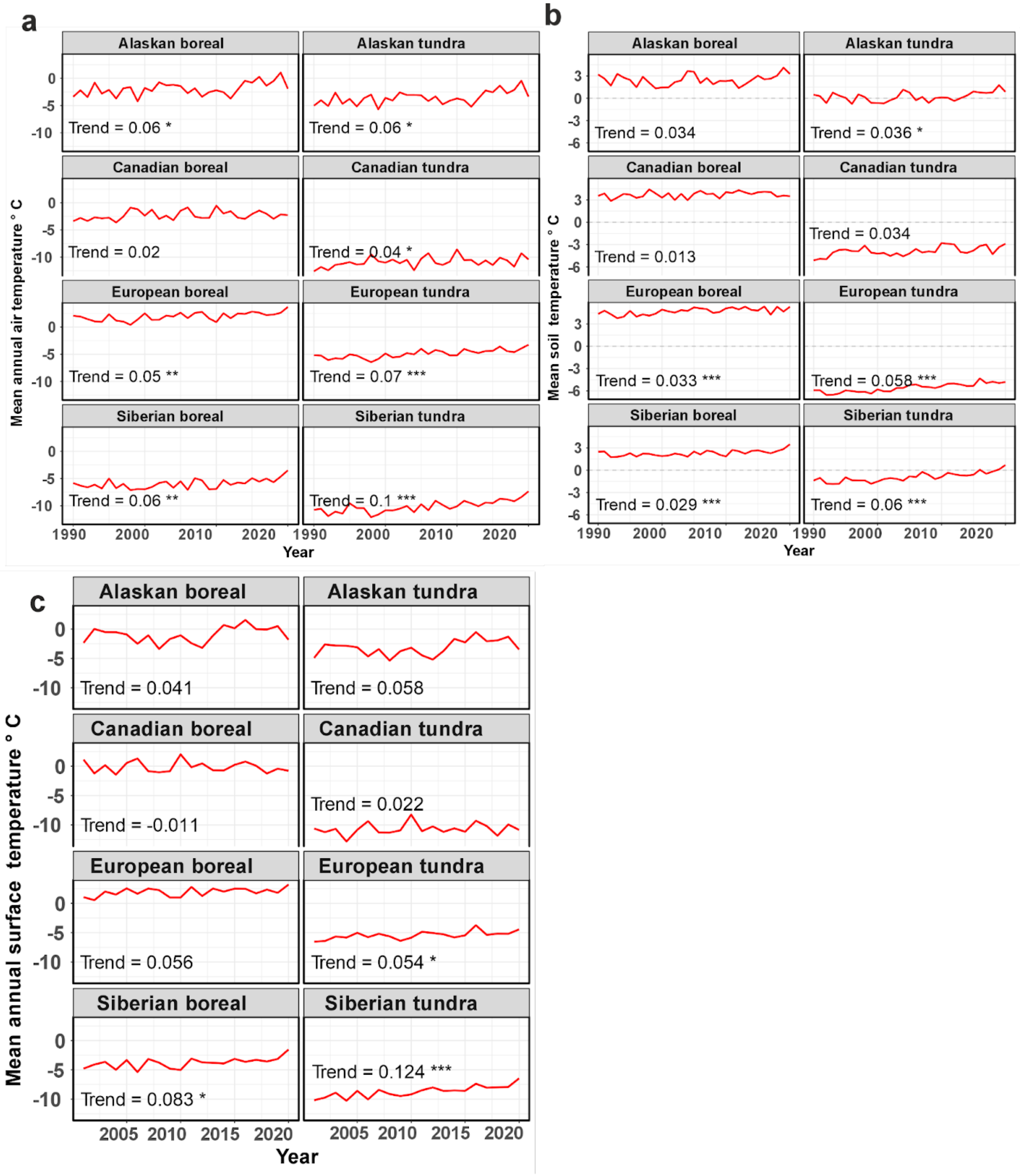

Supplementary Fig. 12. Trends (°C $yr^{-1}$) for the air, land surface and soil temperature variables included in the models. Figures show that all regions are showing increases in air and soil temperatures. The Siberian tundra has the strongest air and soil temperature trends whereas the Canadian boreal has the weakest non-significant trends. Stars in the trend values depict the significance of the trend (*= p<0.05, **=p<0.01, ***=p<0.001).

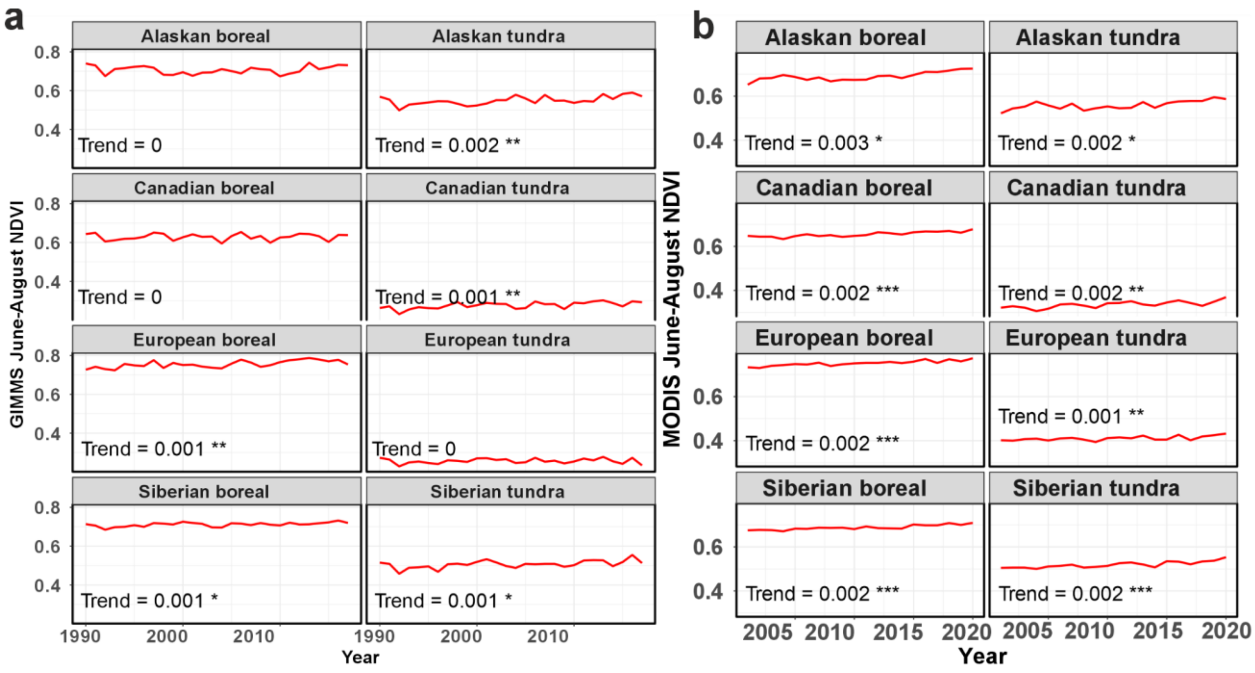

Supplementary Fig. 13. Trends for the NDVI variables for the June-August period. The GIMMS dataset used here covers a longer time period (1990-2016) but is limited to 8-km pixel resolution, whereas the MODIS NDVI time series goes from 2001 to 2020 and is at 1-km pixel resolution. Average greening trends are almost equally strong across the regions in the MODIS era but there is more variability in the GIMMS era. Weakest trends are found in European tundra across both the datasets; GIMMS shows strong trends particularly in Alaskan tundra. Stars in the trend values depict the significance of the trend (*= p<0.05, **=p<0.01, ***=p<0.001).

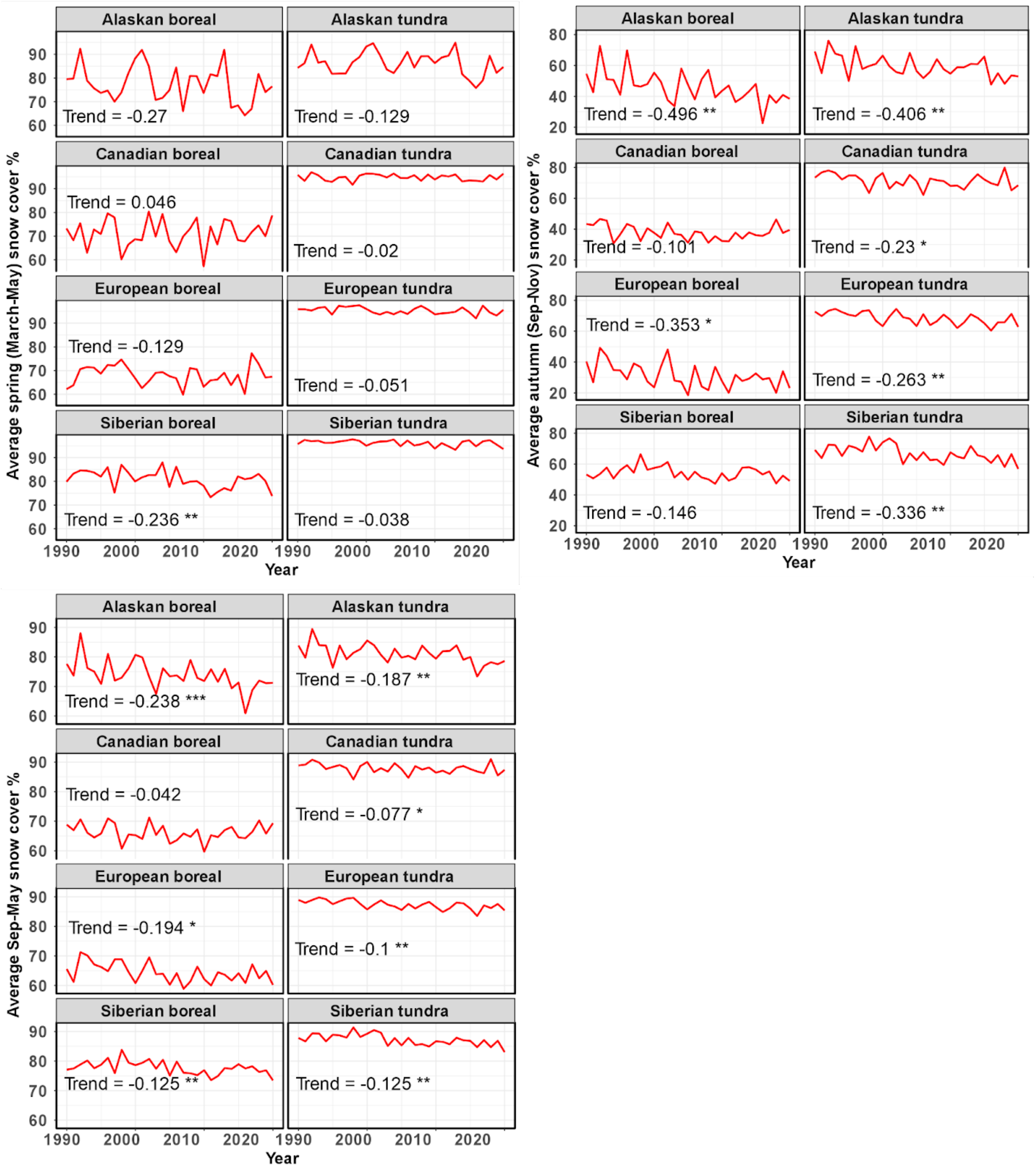

Supplementary Fig. 14. Trends (% $yr^{-1}$) for the snow cover variable included in the models show that all regions experience declining snow cover in spring, autumn, and the entire non-summer (September-May) season. However, snow cover trends are stronger and statistically significant primarily in the autumn season, except for the Siberian boreal region that experiences a strong statistically significant declining trend in the spring. Declines in snow cover are the steepest in Alaskan boreal and tundra regions. Stars in the trend values depict the significance of the trend (*= $p<0.05$, **=$p<0.01$, ***=$p<0.001$).

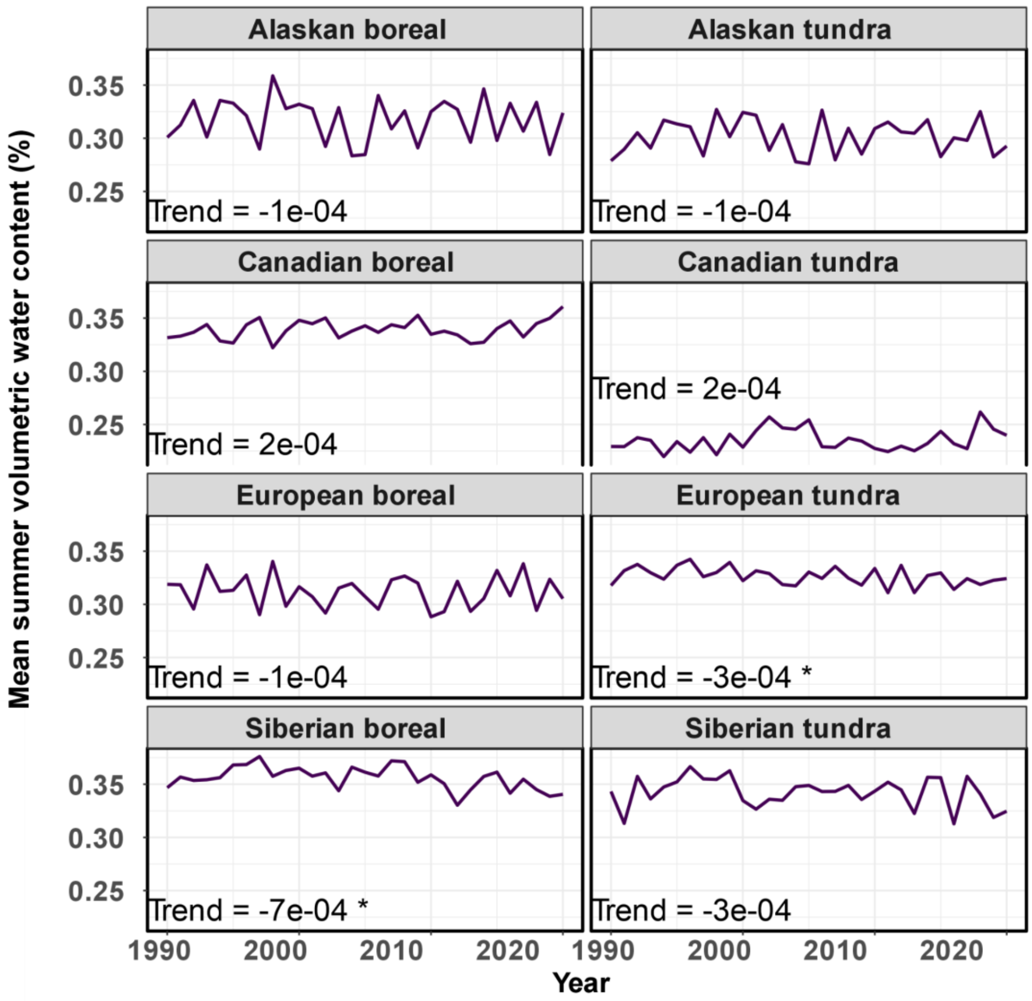

Supplementary Fig. 15. Trends (% yr$^{-1}$) for June-August soil moisture. Stars in the trend values depict the significance of the trend (*= p<0.05, **=p<0.01, ***=p<0.001).

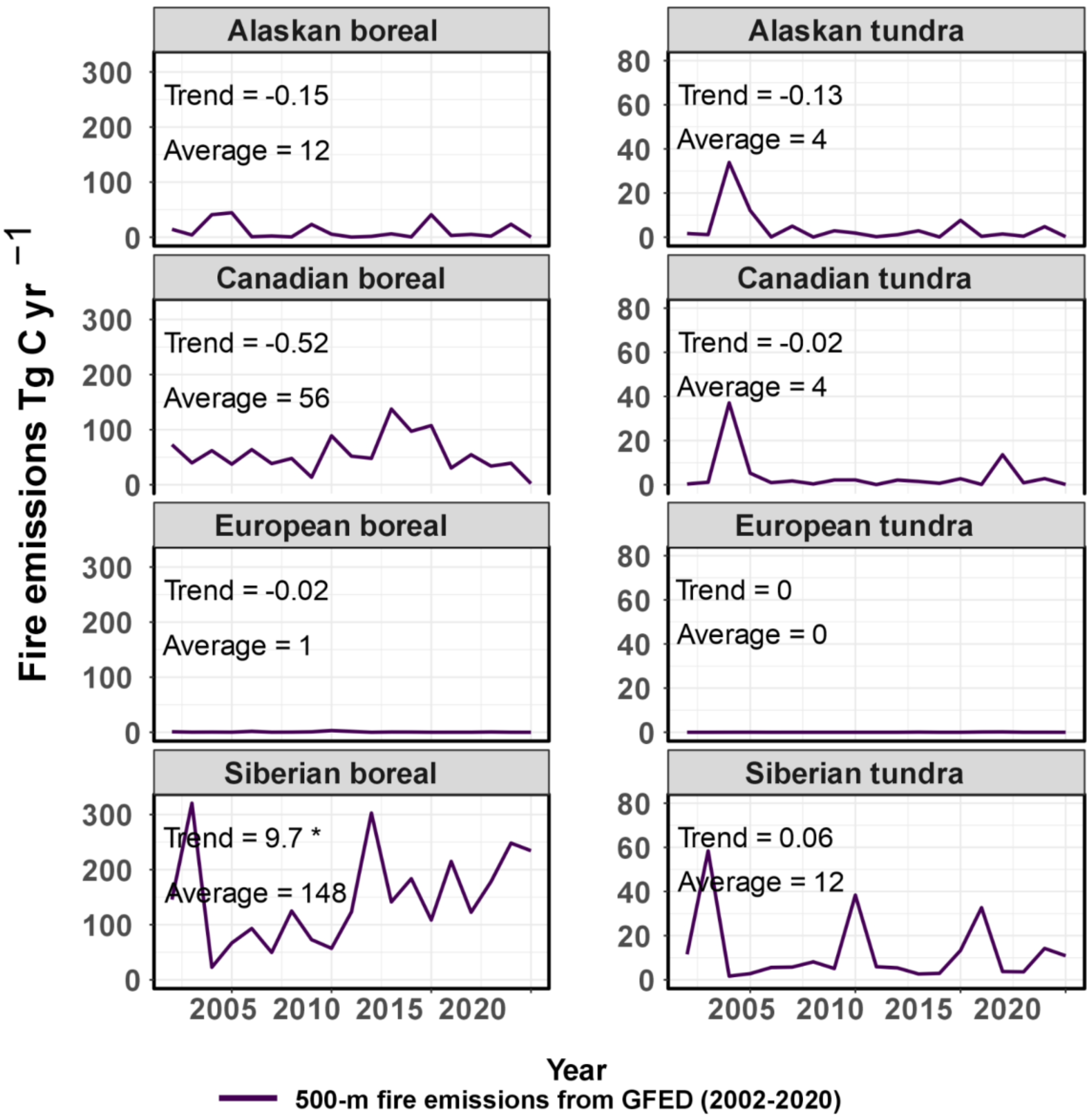

Supplementary Fig. 16. Annual fire emission budgets across the key domains. Stars in the trend values depict the significance of the trend (*= p<0.05, **=p<0.01, ***=p<0.001).

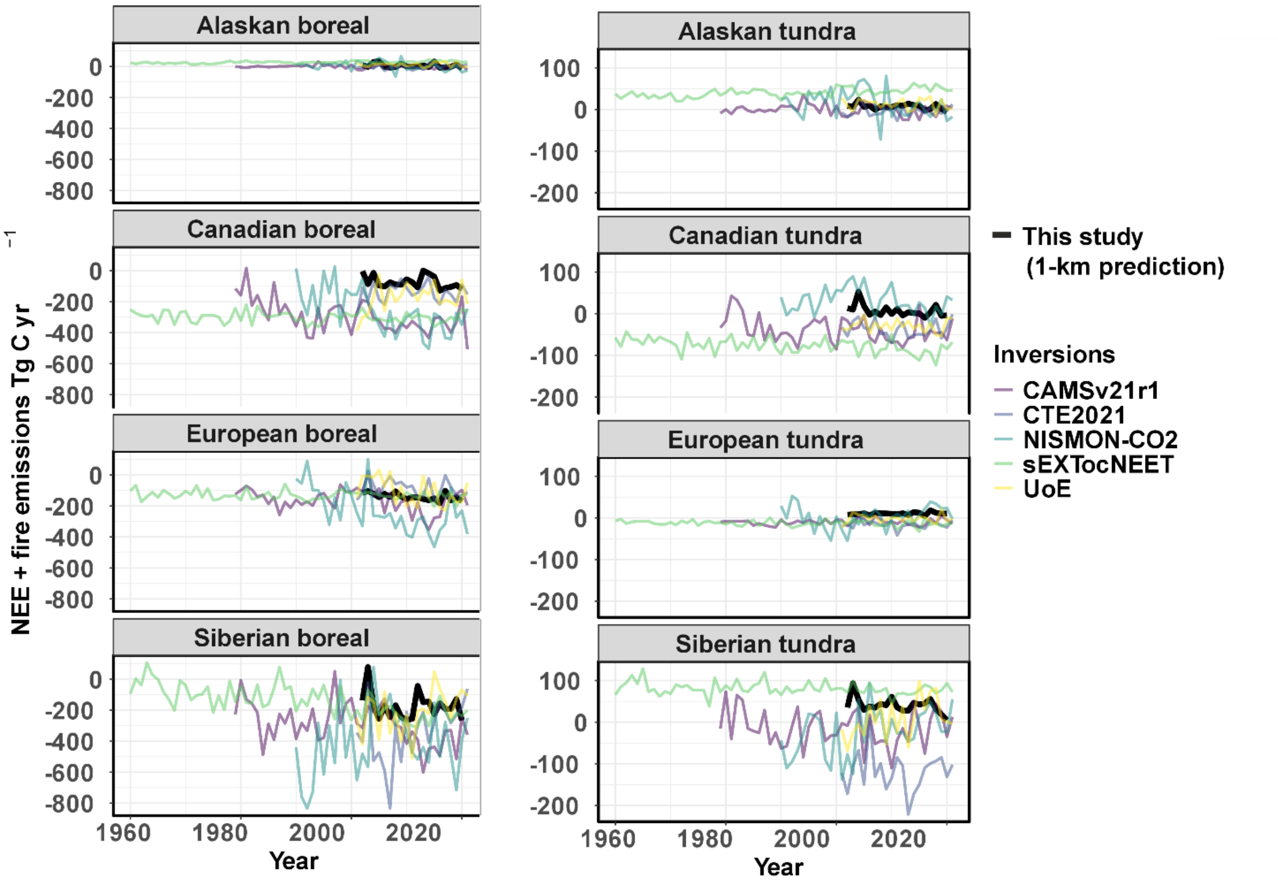

Supplementary Fig. 17. Time series of NEE + fire emissions from the 1-km predictions produced in this study and the atmospheric inversions.

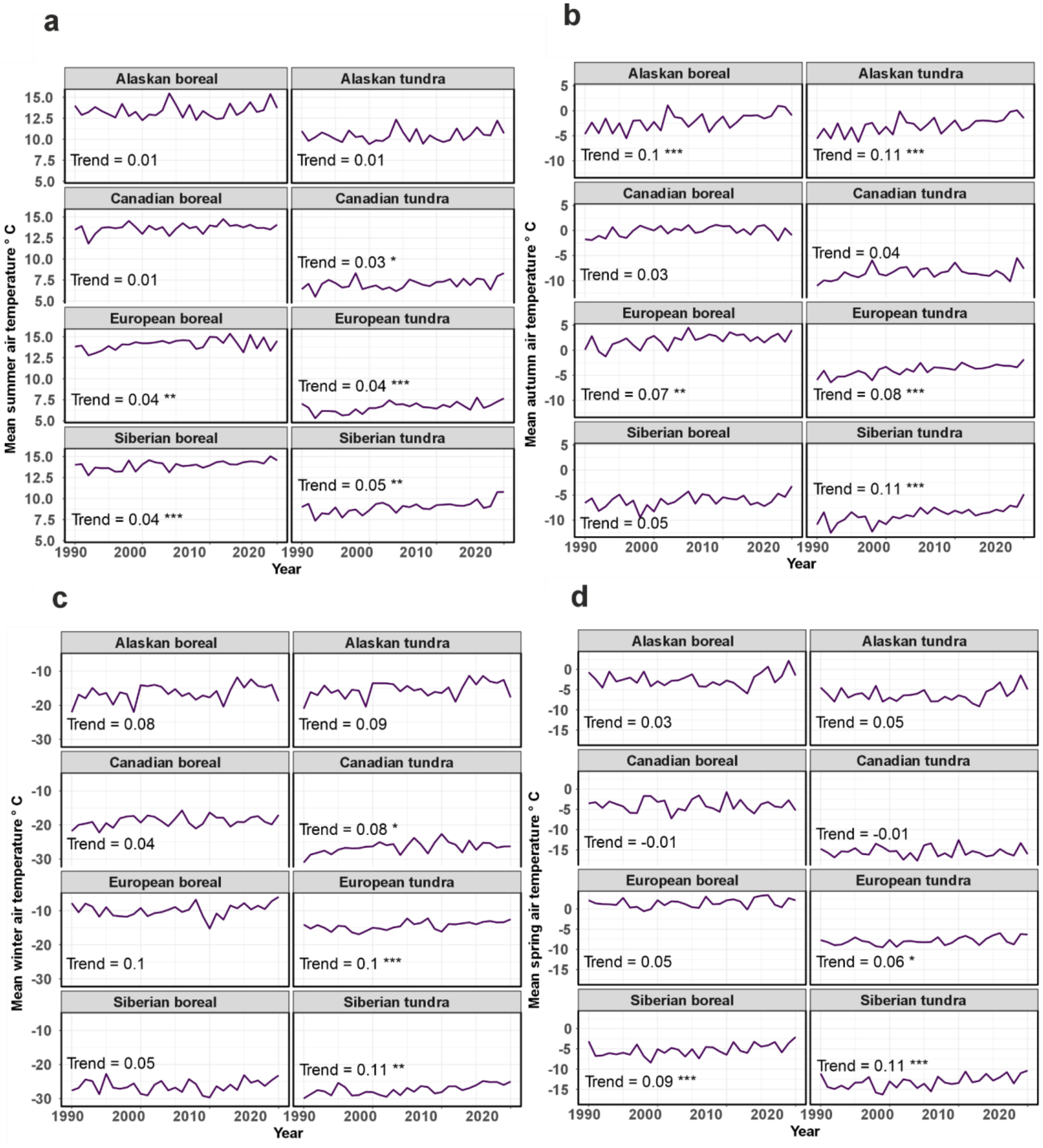

Supplementary Fig. 18. Trends (°C $yr^{-1}$) for the air temperature variables in different climatological seasons. Stars in the trend values depict the significance of the trend (*= p<0.05, **=p<0.01, ***=p<0.001).

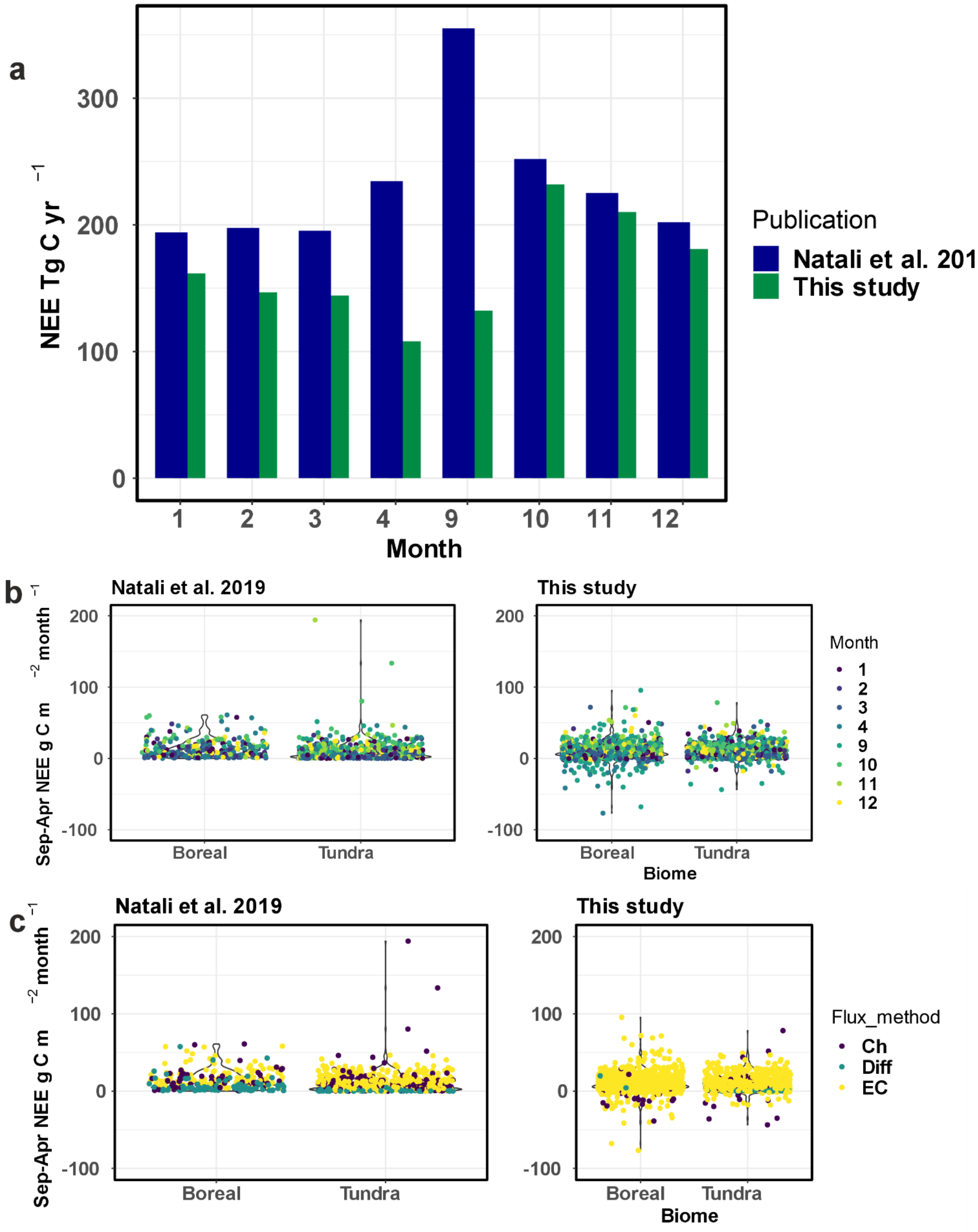

Supplementary Fig. 19. Comparison of the permafrost region non-growing season net ecosystem exchange (NEE) between Natali et al. (2019) and this study across monthly upscaled budgets (a), and in-situ monthly flux data visualized with months (b) and flux measurement methods (c). Natali et al. (2019) removed negative average monthly fluxes during the non-growing season to focus on net emissions, with a total number of site-months being 859 (in this study, site-months in the permafrost region totaled 1702). The October-April budget in this study was 1,181 Tg C $yr^{-1}$ compared with 1,501 Tg C $yr^{-1}$ in Natali et al. (2019) for the same period and domain.

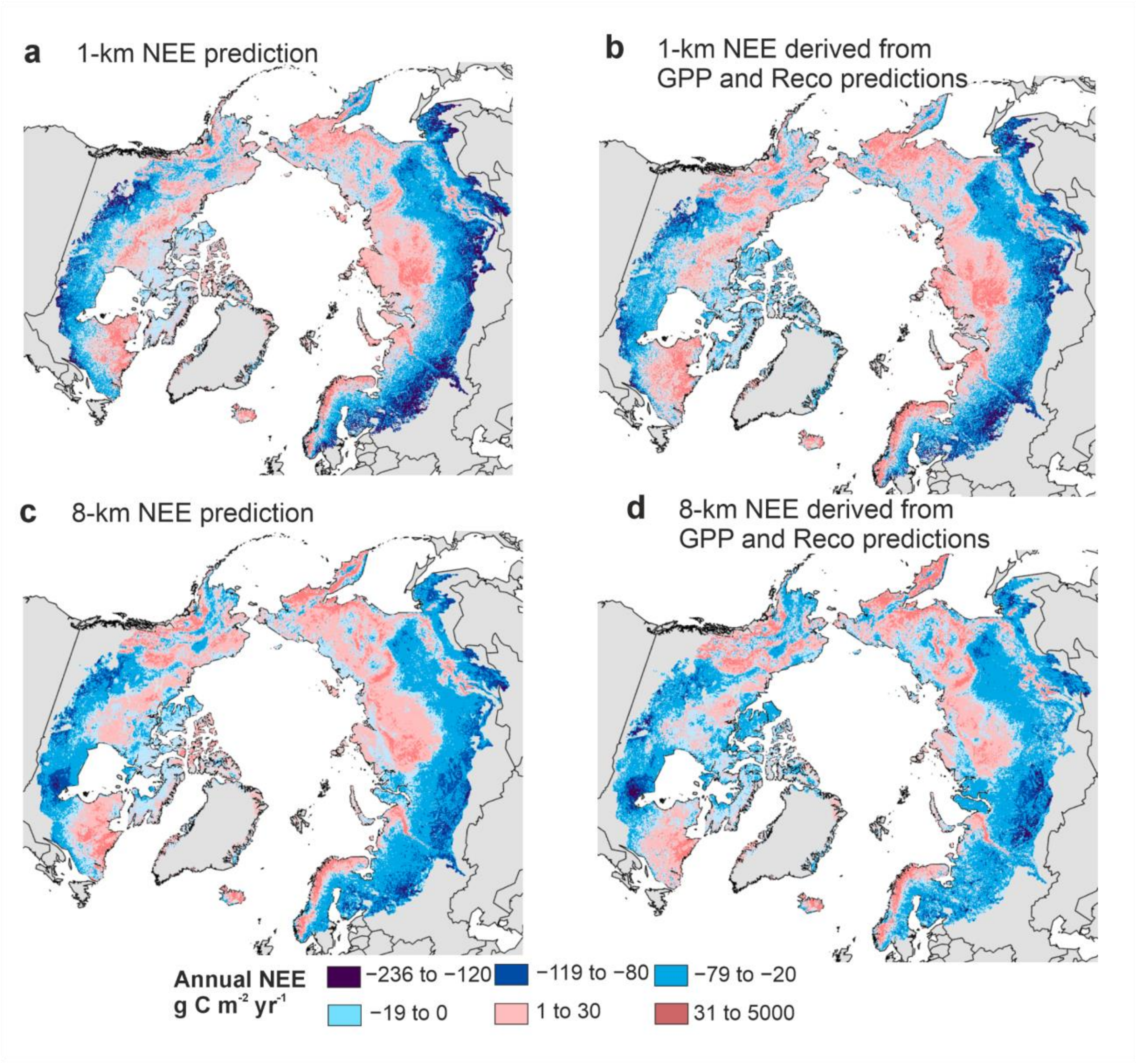

Supplementary Fig. 20. A comparison of upscaled NEE at 1 and 8-km predictions and from directly derived NEE or GPP-$R_{eco}$ derived NEE.

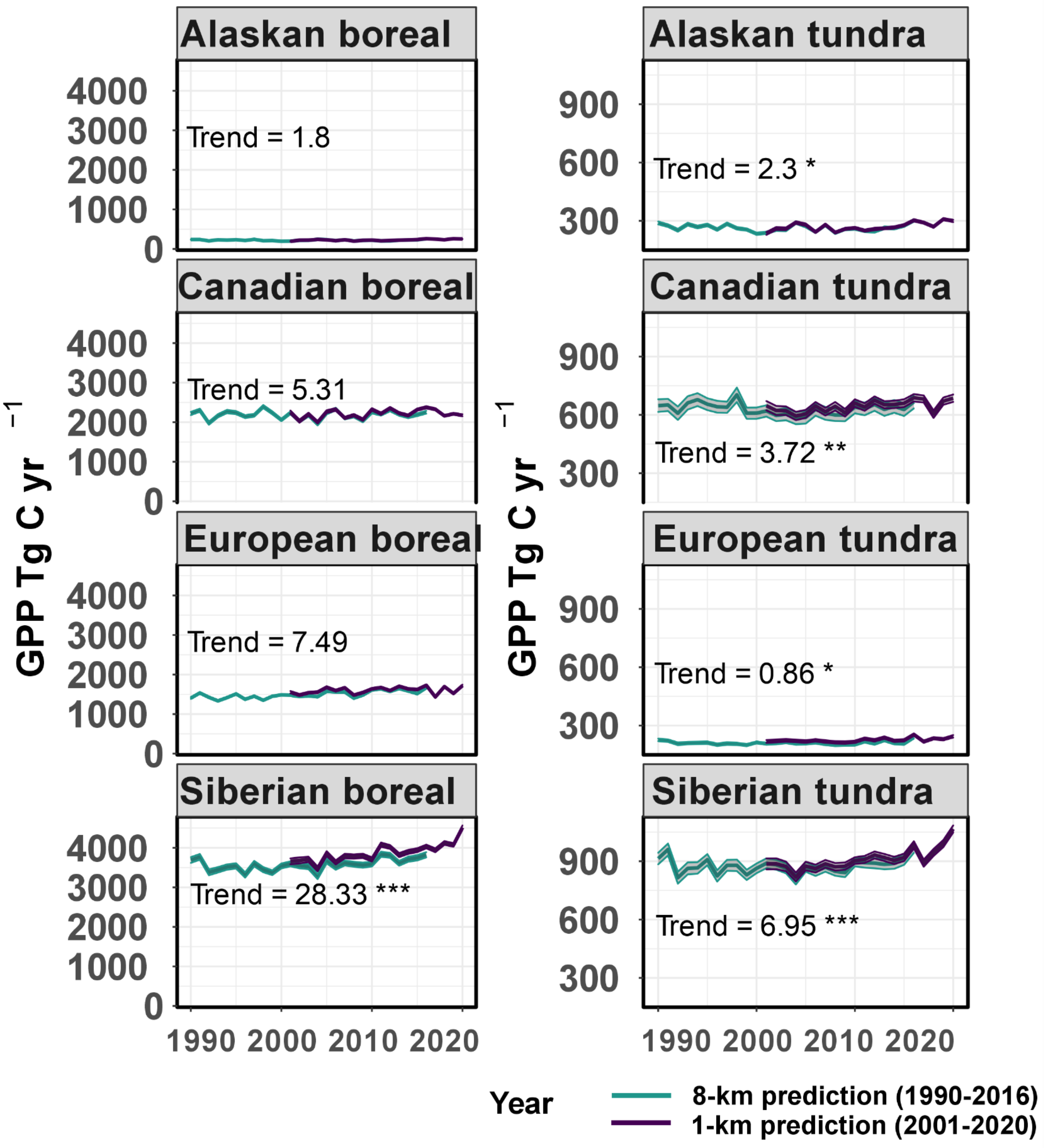

Supplementary Fig. 21. Time series of GPP across the key regions. Stars in the trend values depict the significance of the trend (*= p<0.05, **=p<0.01, ***=p<0.001).

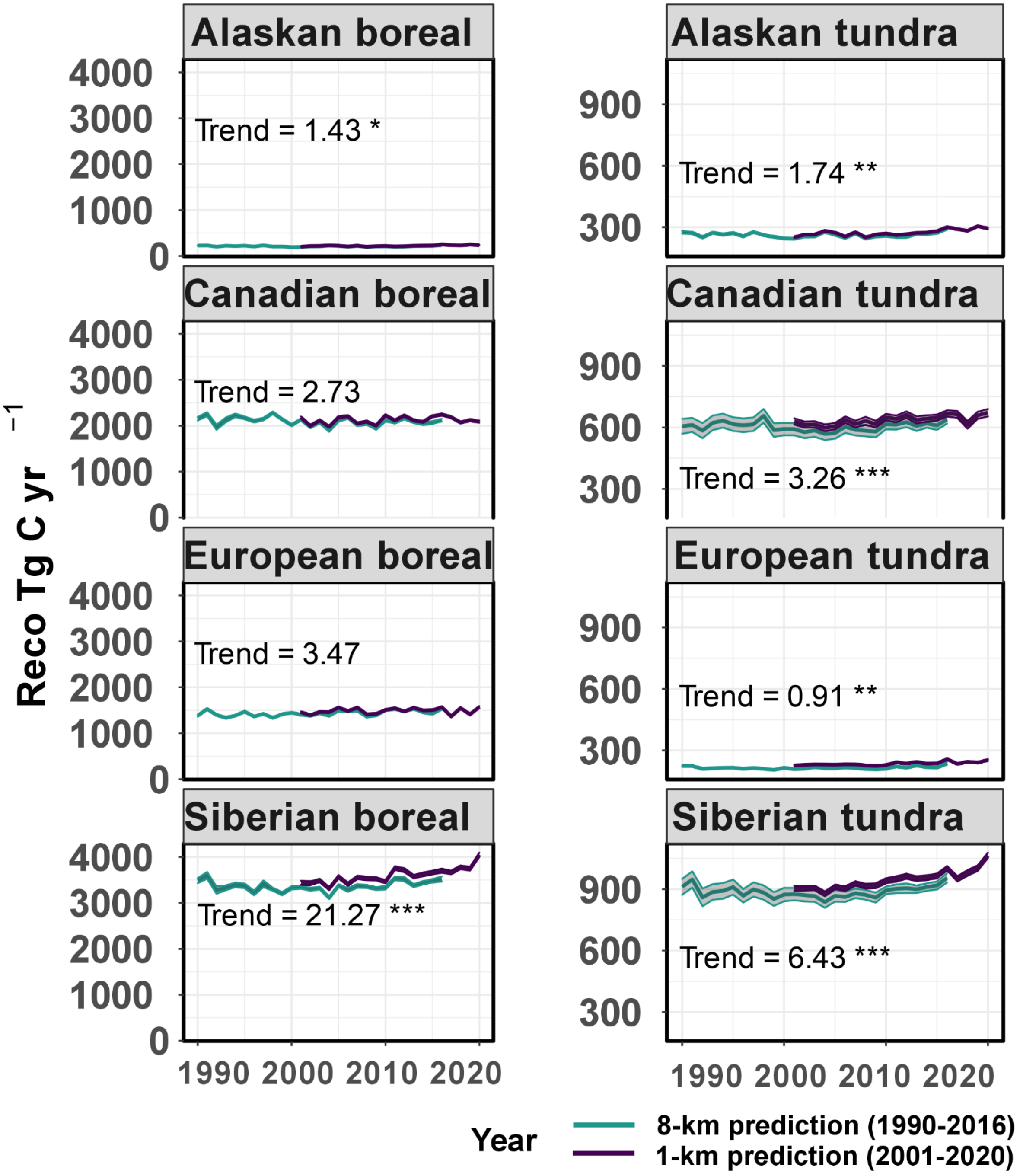

Supplementary Fig. 22. Time series of $R_{eco}$ across the key regions. Stars in the trend values depict the significance of the trend (*= p<0.05, **=p<0.01, ***=p<0.001).

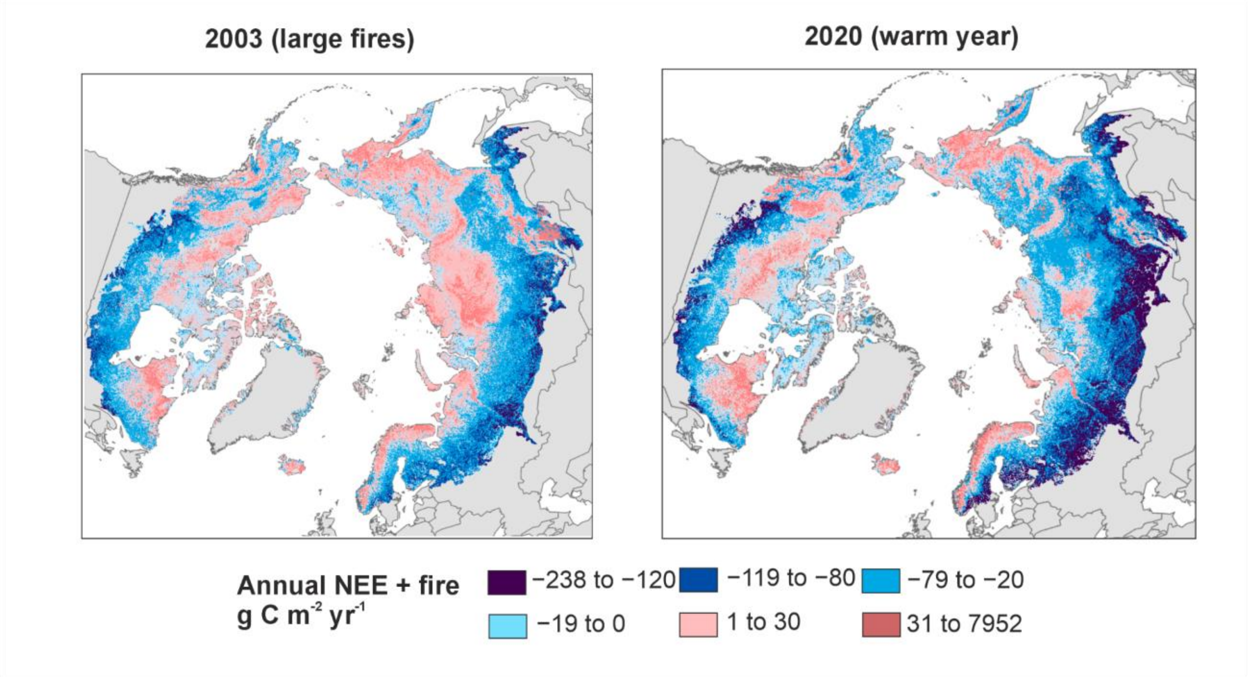

Supplementary Fig. 23. A visualization of how NEE + fire fluxes vary in 2003, when net $CO_2$ emission budget was the highest, and in 2020, when net $CO_2$ emission budget was the lowest.

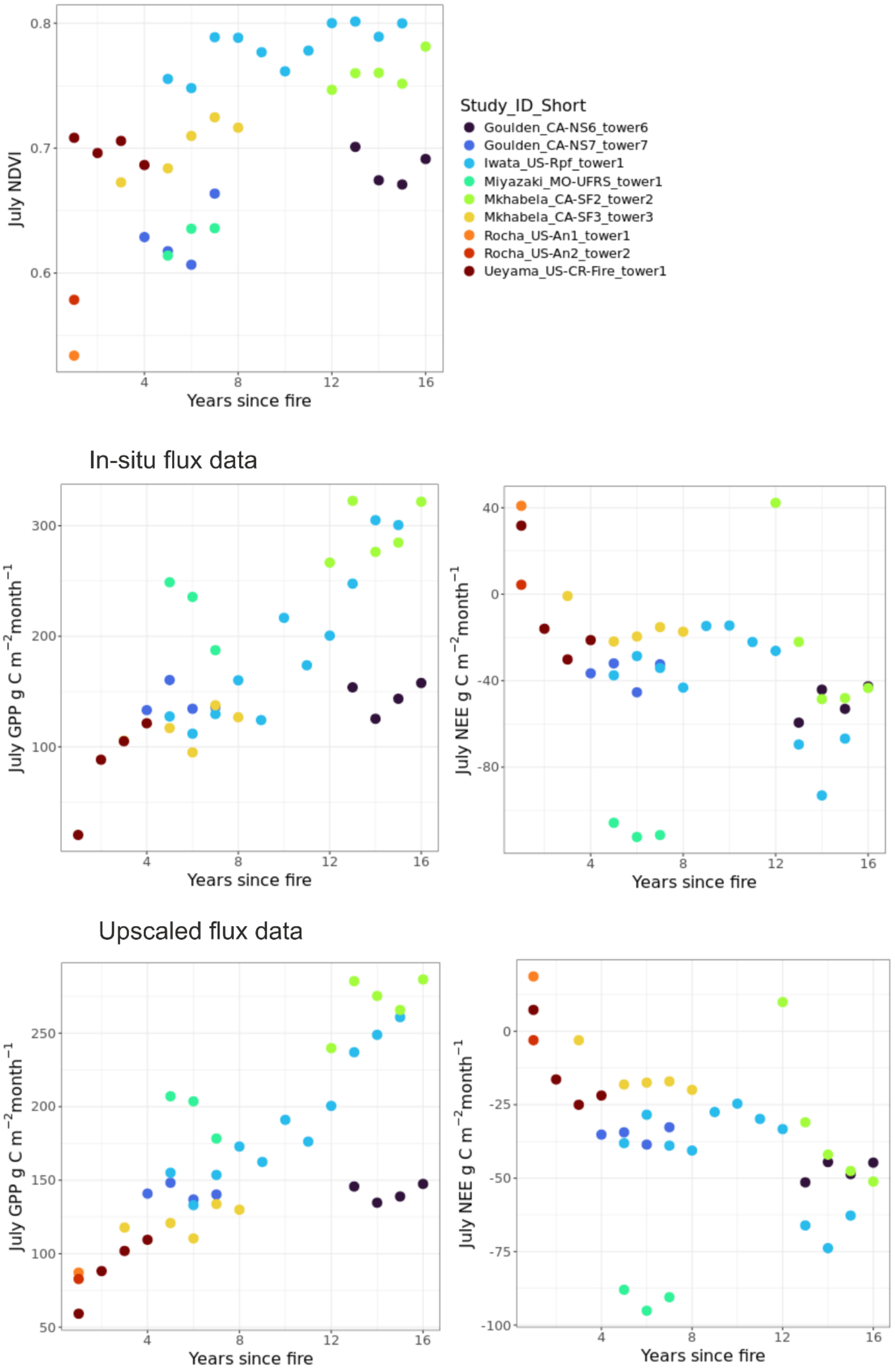

Supplementary Fig 24. Site-level remote sensing-based NDVI, and in-situ and upscaled flux data in recently burned sites showing a clear increasing NDVI, GPP and net uptake signal after the fire. The upscaled fluxes are showing the actual predictions and not those based on cross validation.

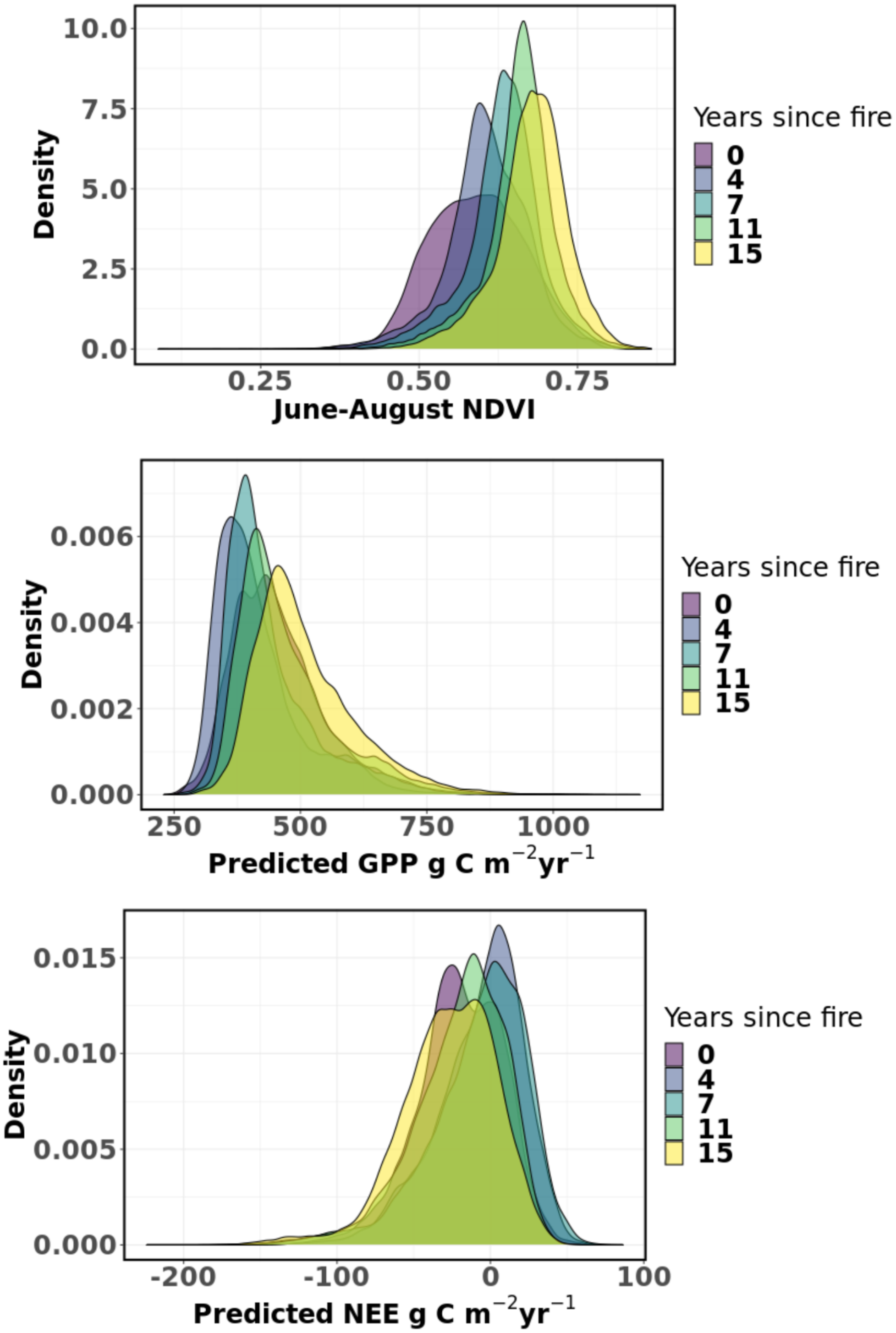

Supplementary Fig. 25. The Kernel density distribution of June-August NDVI, predicted annual GPP, and predicted annual NEE across pixels that were burned in 2004 from 0 to 15 years after the fire. Burned pixels were masked based on the pixels that had a direct fire emission of >200 g C $m^{-2}$ $yr^{-1}$ in the GFED-500m product and were located mainly in Alaska and northwestern Canada.

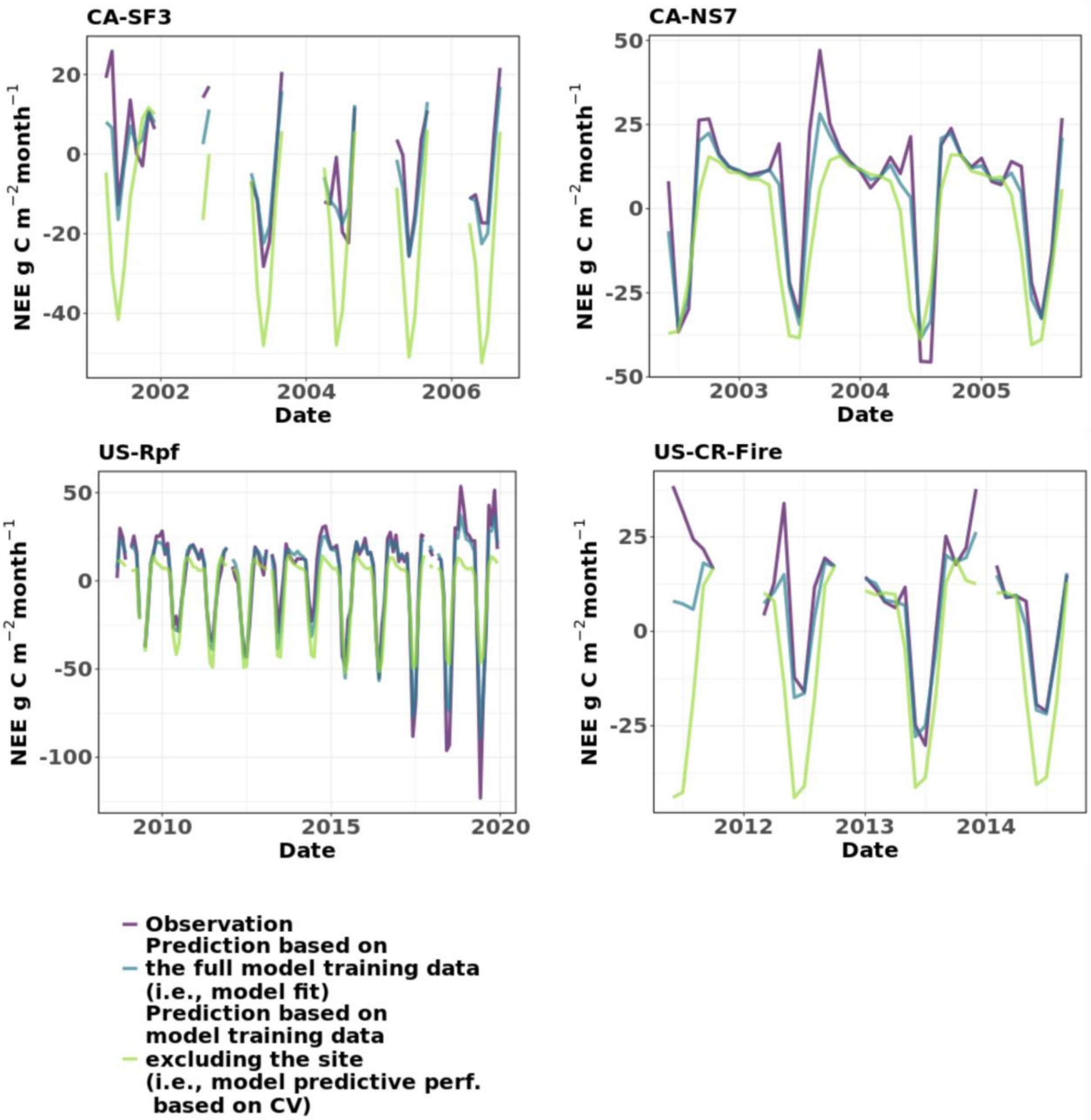

Supplementary Fig. 26. Time series of NEE from recently burned long-term (>3 years) sites and their agreement with model predictions. Model fit indicates how well the model trained with the entire model training data predicts the same data and model predictive performance shows how the models perform when a dataset excluding the specific site is used to train the model.

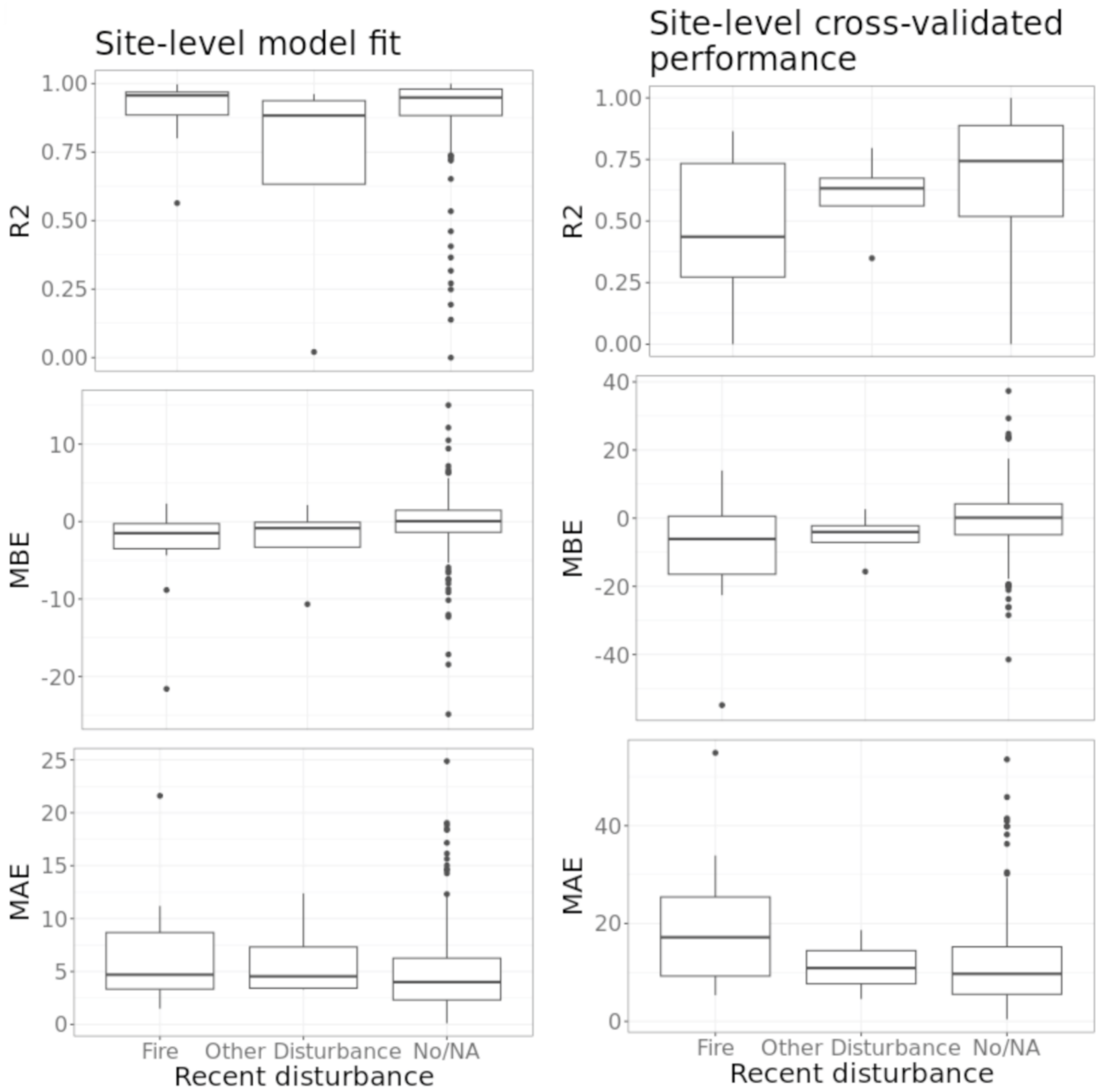

Supplementary Fig. 27. Model fit (i.e., no cross validation) and cross-validated predictive performance estimates for each site across sites with recent fire disturbance, other disturbance (e.g., permafrost thaw, drainage), or no disturbance or information about disturbance. The boxes correspond to the 25th and 75th percentiles, and the line within the box represents the median. The lines denote the 1.5 IQR of the lower and higher quartile, where IQR is the inter-quartile range. Coefficient of determination ($R^2$) describes the strength of the linear relationship between the observed and predicted fluxes. Mean bias error (MBE) characterizes the average bias between prediction and observation, with negative values indicating the model to underestimate NEE (i.e., overestimate net CO2 sinks or underestimate net CO2 sources). Mean absolute error (MAE) describes the absolute bias between prediction and observation, with larger values describing larger errors.

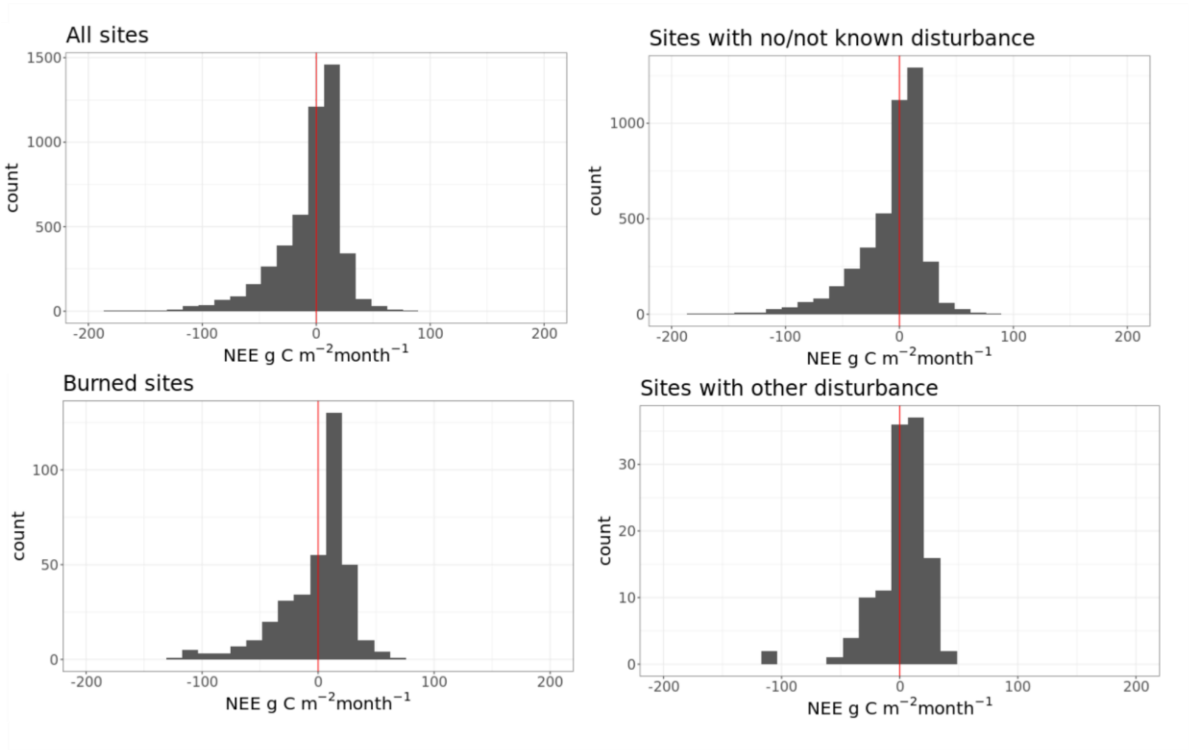

Supplementary Fig. 28. The distribution of monthly in-situ NEE data used to train the models. The red vertical line shows the zero flux.

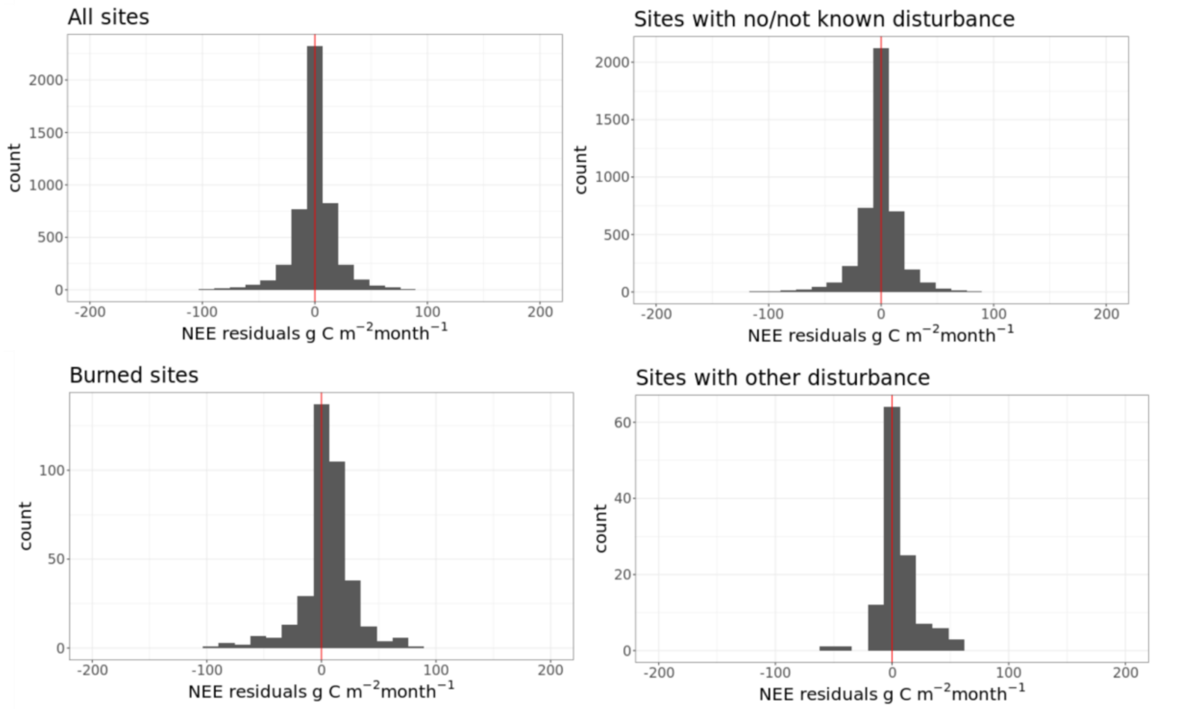

Supplementary Fig. 29. The distribution of monthly residuals of NEE. The red vertical line differentiates positive and negative residuals. A negative residual value represents the model overestimating NEE, i.e., underestimating net $CO_2$ sinks or overestimating net $CO_2$ sources.

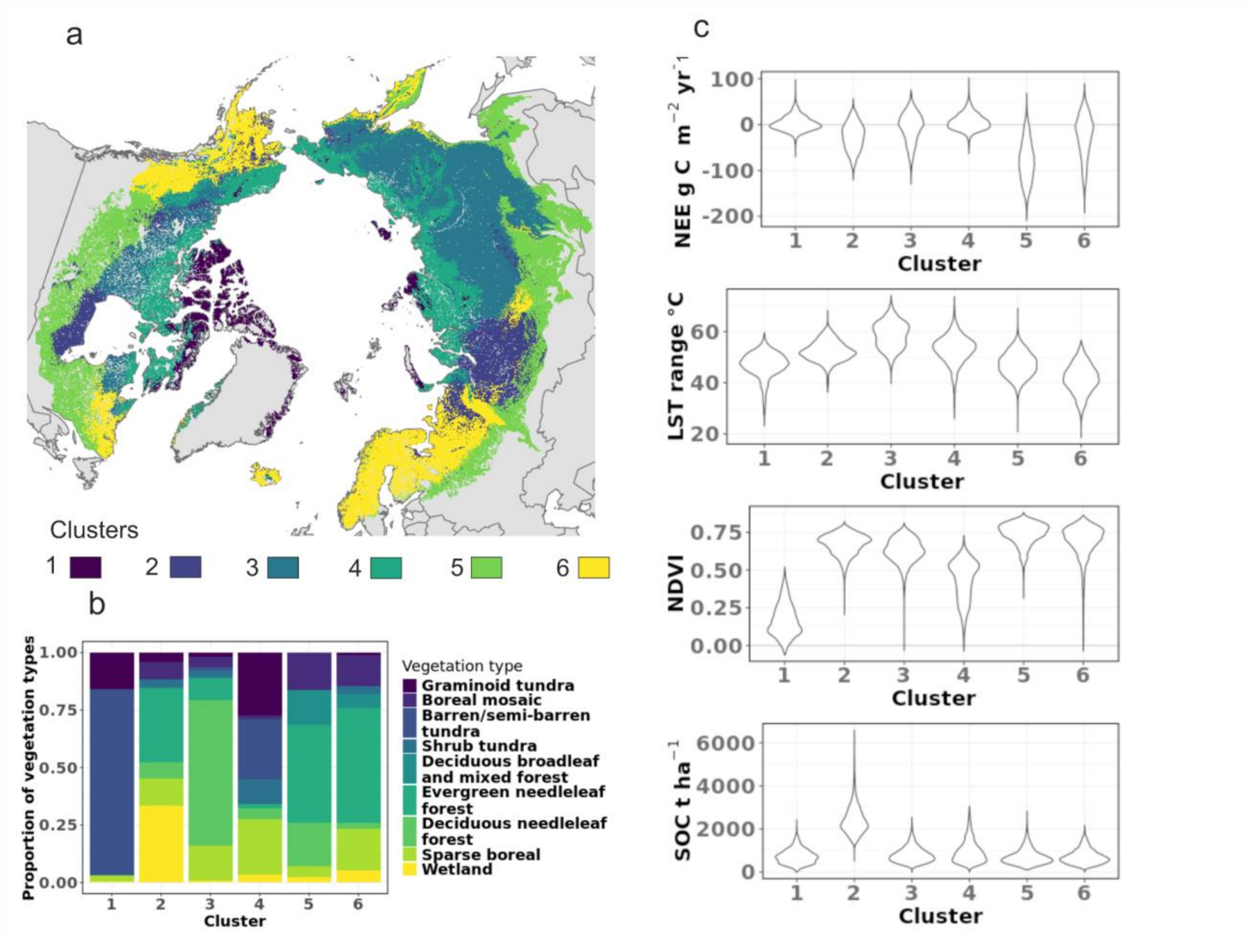

Supplementary Fig. 30. Environmental clustering of the ABZ based on the most important average annual geospatial predictors from upscaling (a), the percentage of vegetation types within each cluster (b), and the variability in key environmental conditions across the clusters based on a random spatial sample of 10,000 pixels per cluster (c).

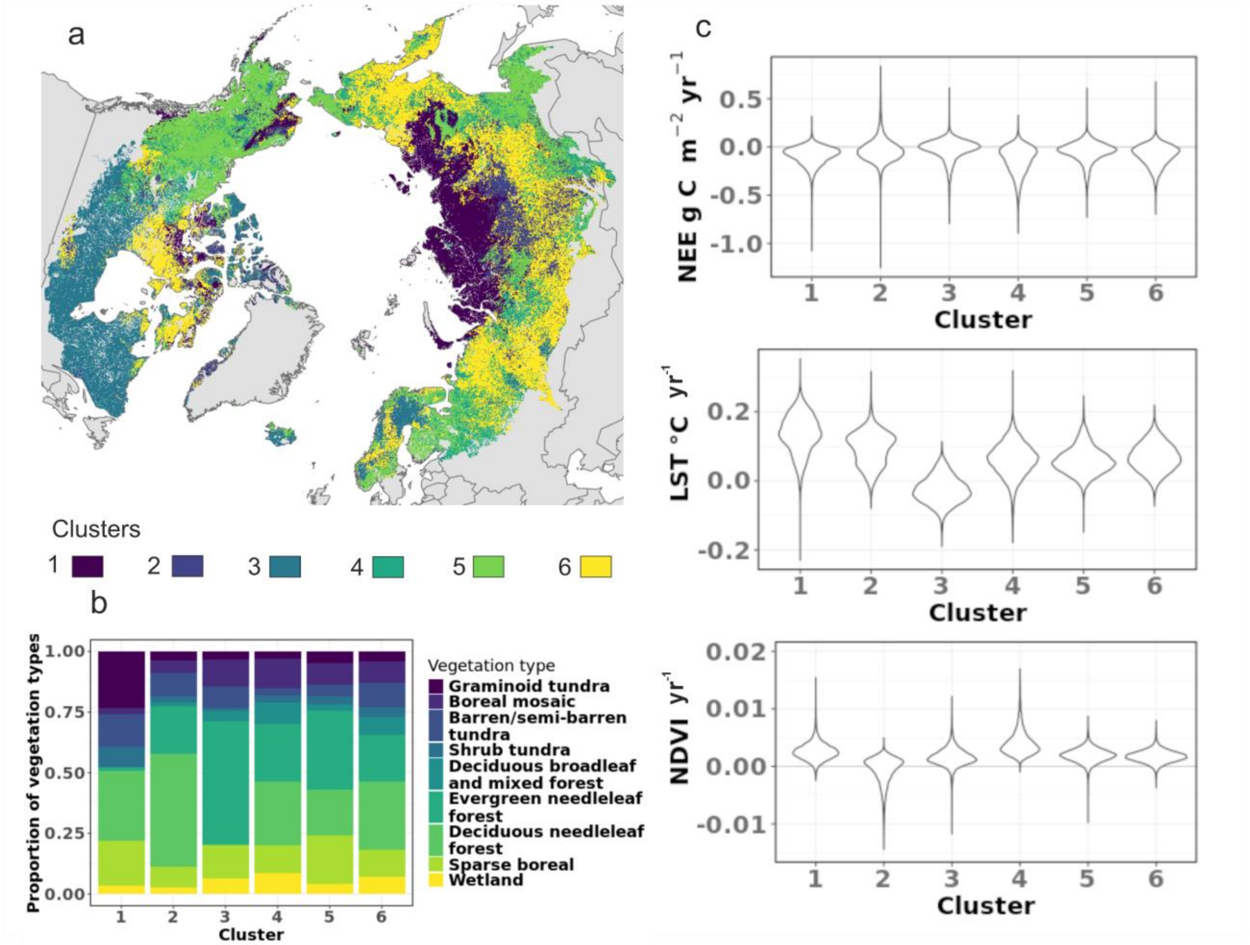

Supplementary Fig. 31. Environmental clustering of the temporal trends in ABZ based on the most important average annual geospatial predictors from upscaling (a), the percentage of vegetation types within each cluster (b), and the variability in key environmental trends (per year) across the clusters based on a random spatial sample of 10,000 pixels per cluster (c).

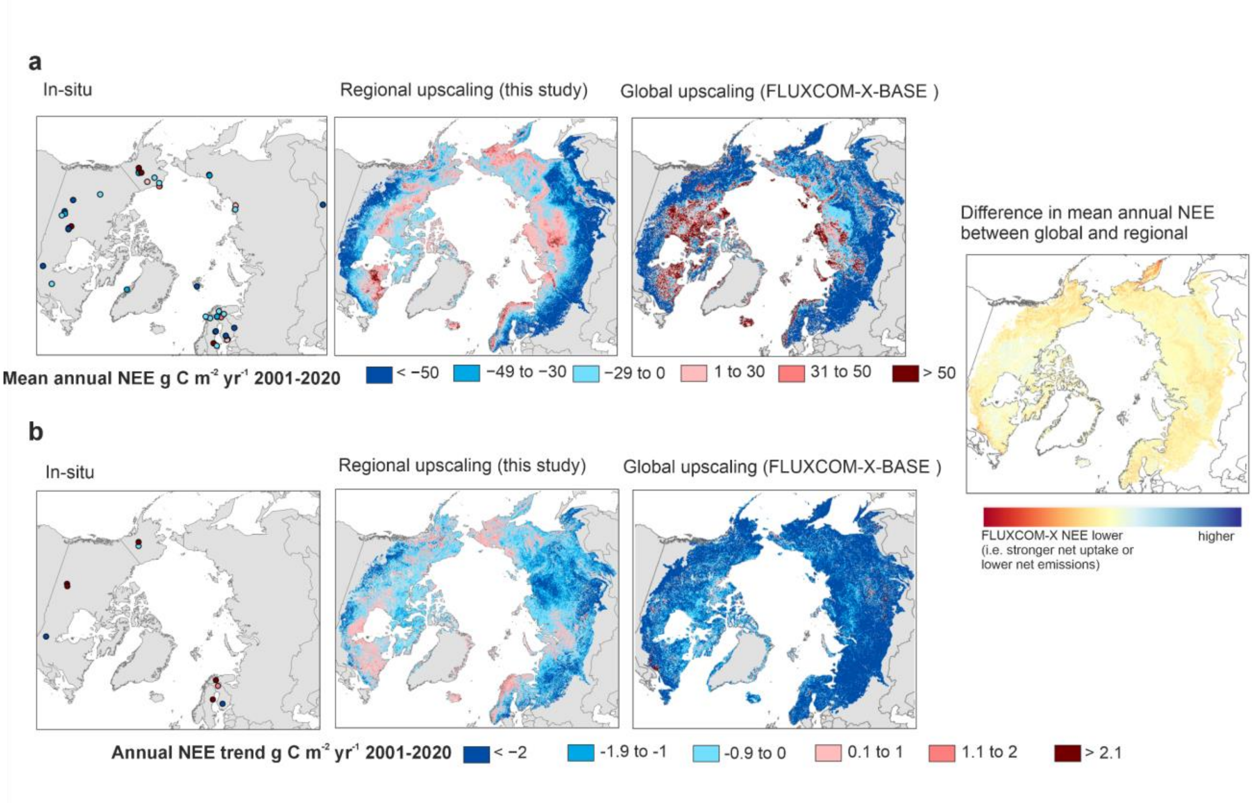

Supplementary Fig. 32. Maps showing the averages and trends in annual NEE in our upscaling and FLUXCOM-X-BASE, and the differences between those.

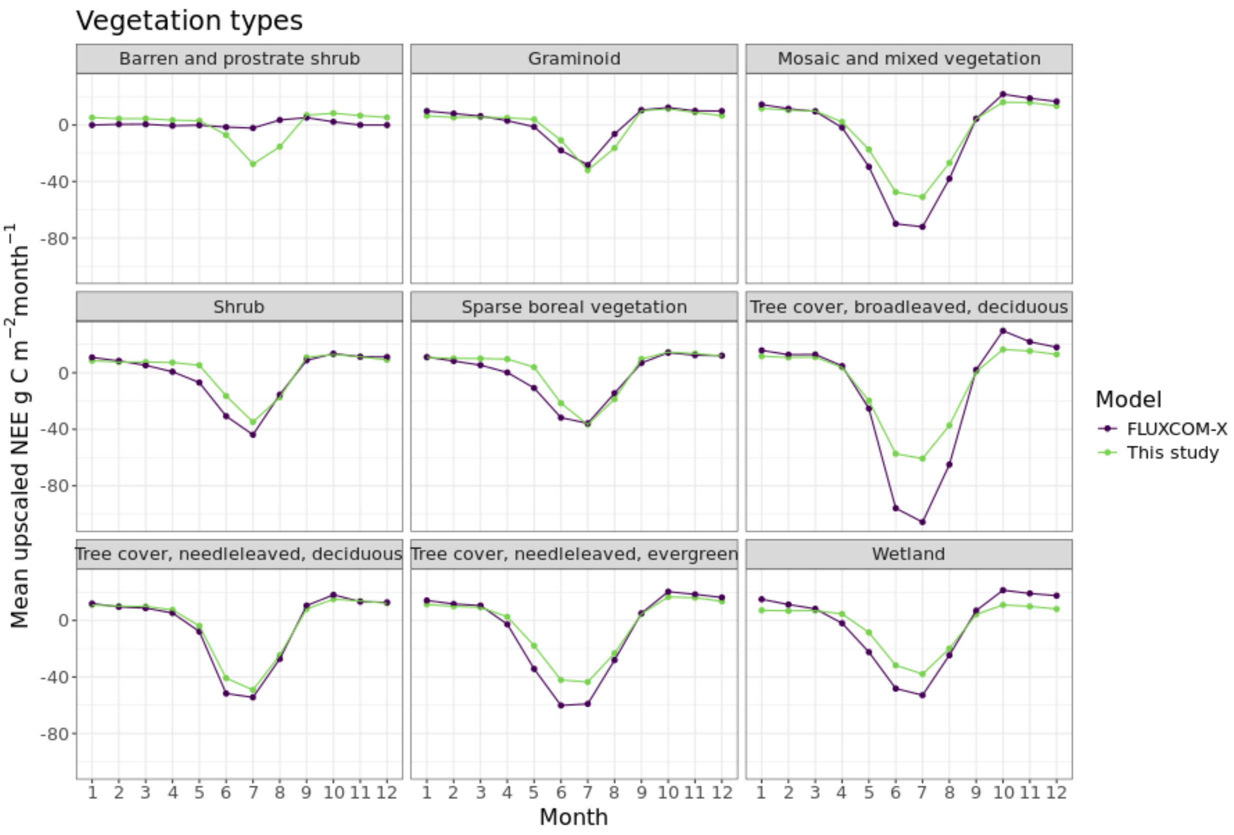

Supplementary Fig. 33. Mean upscaled monthly NEE fluxes in this study and FLUXCOM-X-BASE across key vegetation types.

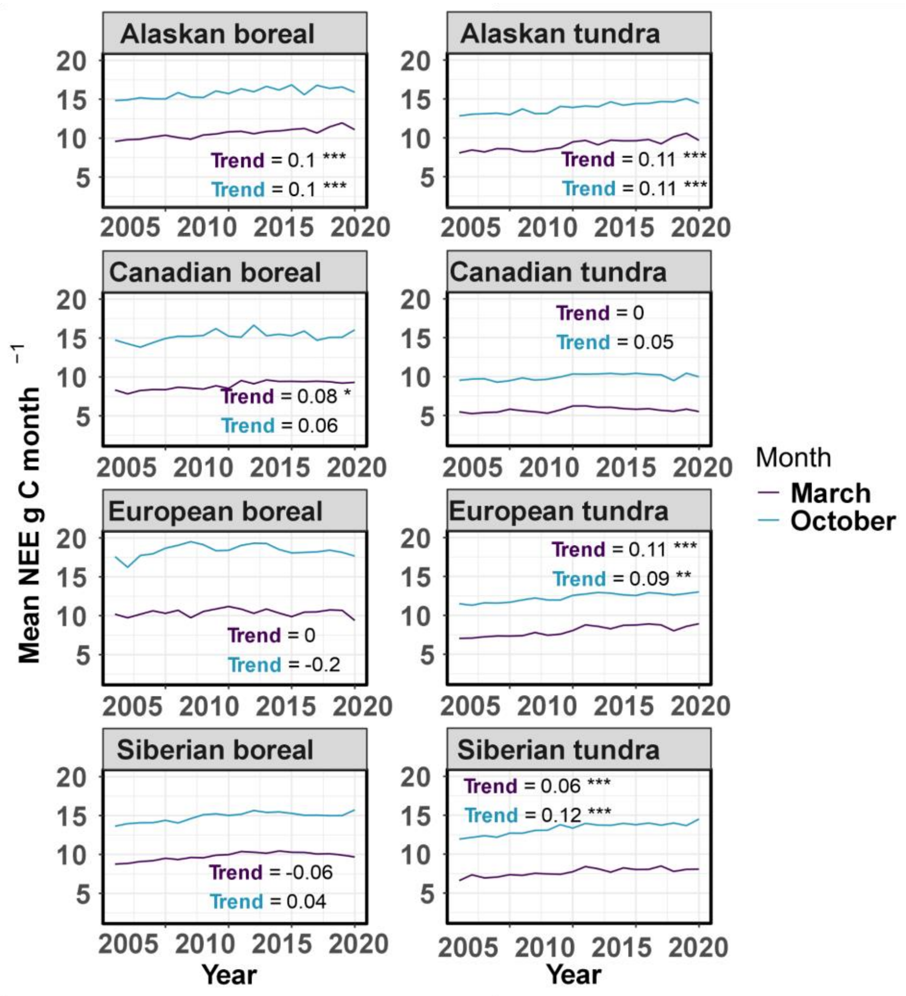

Supplementary Fig. 34. Trends in upscaled shoulder season mean NEE fluxes in key regions from 2001 to 2020.
