## Supplementary material for "An increasing Arctic-boreal CO_2_ sink offset by wildfires and source regions": Analysis codes

```
### load packages
library("dplyr")
library("caret")
library("parallel")
library("doParallel")
library("randomForest")
library("groupdata2")
library("itertools")
library("foreach")
library("doParallel")
library("parallel")
library("sp")
library("ggplot2")
library("caret")
library("dplyr")
library("purrr")
library("raster")
library("terra")
library("stringr")
library("vip")
library("pdp")
library("Metrics")
library("tdr")
library("viridis")
library("tidyr")
library("ggridges")
library("zyp")


### MODELING ###

### Load the model training data
d <- read.csv("/home/master/flux_upscaling_data/results/final/modeldata_avg_allsites_1km.csv")

# Define the factors
d$Study_ID_Short <- factor(d$Study_ID_Short)
d$ESACCI_cavm_general_ESAwaterfix_broadevfix_mixfix_cropfix_nowaterglacier_ESACCI_CAVM_mer
ged <-

as.factor(d$ESACCI_cavm_general_ESAwaterfix_broadevfix_mixfix_cropfix_nowaterglacier_ESACCI_C
AVM_merged)
d$Interval <- as.factor(d$Interval)


# Response variables
resp_vars <- c("NEE_gC_m2",  "GPP_gC_m2", "Reco_gC_m2") #

# Predictors
Baseline_vars_1km <- c("Interval", "srad_terraclimate_sites", "vpd_terraclimate_sites",

                       "Barrow_CO2_conc_Barrow_CO2conc",

"ESACCI_cavm_general_ESAwaterfix_broadevfix_mixfix_cropfix_nowaterglacier_ESACCI_CAVM_merge
d",
```

```
                    "Snow.cover_era5_soilmoist_temp_snow",
"Soil.temperature.level.1_era5_soilmoist_temp_snow",
                    "Volumetric.soil.water.layer.1_era5_soilmoist_temp_snow",

                    "NDVI_whittaker_constant_monthly_mean",

                    "LST_Day_1km_MOD11A2v006_LST_Day_sites_low_quality",


                    "dtm_cti_merit.dem_m_250m_s0..0cm_2018_v1.0_MERIT_topo_indices_250m",


                    "Percent_NonTree_Vegetation_MOD44B_sites", "Percent_NonVegetated_MOD44B_sites",
                    "Percent_Tree_Cover_MOD44B_sites",


                    "PHIHOX_M_sl1_250m_ll_SoilGrids", "SoilGrids_SOC_SoilGrids_SOCstock",


"UiO_PEX_PERPROB_5.0_20181128_2000_2016_NH_UiO_PEX_20181128_2000_2016_NH" #
Permafrost (static)



)



# Set up clusters for parallel processing
#  https://stackoverflow.com/questions/44774516/parallel-processing-in-r-in-caret
library(parallel)
# Calculate the number of cores
no_cores <- detectCores() - 1
library(doParallel)
# create the cluster for caret to use
cl <- makePSOCKcluster(no_cores)
registerDoParallel(cl)
clusterEvalQ(cl, .libPaths("/home/master/R/x86_64-pc-linux-gnu-library/4.2"))



### Loop through the response variables and tune the models
for (flux in resp_vars) {

  print(flux)

  modeldata3 <- d[,c("Study_ID_Short", "id", flux, Baseline_vars_1km)]
  modeldata2 <- na.omit(modeldata3)
  sapply(modeldata2, function(x) sum(is.na(x))) # no missing data


  # Create a list of row indices for cross validation splits (for leave-one-fold out)
  # leave one site out used because of the highly different number of observations in some factor data
levels

  set.seed(448)
```

```
folds <- groupdata2::fold(data= modeldata2,
                    k = length(unique(modeldata2$Study_ID_Short)), id_col = "Study_ID_Short")

# merge
folds_data <- subset(folds, select=c( ".folds", "id"))
modeldata2 <- full_join(modeldata2, folds_data, by="id")

# create a row ID
modeldata2$samplerow <- seq(1, length(modeldata2$Study_ID_Short), by=1)

indices2 <- list()
indices_not2 <- list()

for (k in unique(modeldata2$.folds)) {

  # k = 1
  subs <- subset(modeldata2, .folds==k)

  # list the row ids for the fold
  sites <- list(as.integer(subs$samplerow) )

  # list other row ids that don't belong to the k fold
  sites_not <- list(as.integer(modeldata2$samplerow[!(modeldata2$samplerow %in% unlist(sites))]))

  # check that all the rows are included
  length(sites_not[[1]]) + length(sites[[1]]) == length(modeldata2$Study_ID_Short)

  # append to list (caret wants these as lists)
  # row indices for each fold, used for model evaluation
  indices2 <- append(indices2, sites)
  # row indices for model training
  indices_not2 <- append(indices_not2, sites_not)

}


names(indices2) <- 1:length(indices2)
names(indices_not2) <- 1:length(indices2)



print("model tuning and feature selection starting")


# Model tuning parameters
tunecontrol2 <- trainControl(
  method = "cv",  # No need to specify this because we use index column which automatically does
leave-site/group-out CV
  verboseIter = FALSE,  # A logical for printing a training log. This could be set to FALSE in the final
model runs
  returnData = FALSE, # A logical for saving the data. This could be set to FALSE in the final model runs
  returnResamp = "final", # A character string indicating how much of the resampled summary metrics
should be saved.
  # p = 0.7, # For leave-group out cross-validation: the training percentage
  summaryFunction = defaultSummary, # a function to compute performance metrics across resamples.
```

```
   selectionFunction = "best", #  chooses the tuning parameter associated with the largest (or lowest for
"RMSE") performance.
   index = indices_not2, # a list with elements for each resampling iteration.  needs to be integer
   indexOut = indices2, # model evaluation
   allowParallel = TRUE, # parallel processing
   savePredictions="final") # an indicator of how much of the hold-out predictions for each resample
should be saved.


  # simple random forest
  rfe_fit2 = train(modeldata2[,Baseline_vars_1km], modeldata2[,flux],
                   trControl = tunecontrol2, # tuning parameters
                   method="rf", importance=TRUE, proximity = FALSE)

  print("model tuning done")

  ### Write the model out
  saveRDS(rfe_fit2, paste0("/home/master/abcfluxv1_modeling/results/", paste(flux, "1km_rf_train_loocv",
sep="_"), ".rds"))


}

# prediction to a raster is done with terra package and predict command:
# predict(rfe_fit2, c(pred_rast, pred_rast2, ...))
# where the predictor raster list has the predictors in the same cropped extent and resolution, and same
names as in rfe_fit2


### MODEL PERFORMANCE ###

for (i in resp_vars) {


  # Add data

  modeldata2 <- d[,c("Study_ID_Short", "id", i, Baseline_vars_1km)]
  modeldata2 <- na.omit(modeldata2)

  # create a row ID
  modeldata2$samplerow <- seq(1, length(modeldata2$Study_ID_Short), by=1)


  # Load model files
  mod <- readRDS(paste0("/home/master/abcfluxv1_modeling_bigfiles/results/", paste(i, "1km_rf",
"train_loocv",  sep="_"), ".rds"))
  mod2 = mod$finalModel #pull out the quantile regression object


  ### Predictive performance plots ###


  # Define x and y lab titles for the plot


  if (i=="GPP_gC_m2") {
```

```
    ylab = expression(paste("Observed GPP g C m"^{-2}, month^{-1}))
    xlab = expression(paste("Predicted GPP g C m"^{-2}, month^{-1}, " (1 km model)"))

  }

  if (i=="NEE_gC_m2") {
    ylab = expression(paste("Observed NEE g C m"^{-2}, month^{-1}))
    xlab = expression(paste("Predicted NEE g C m"^{-2}, month^{-1}, " (1 km model)"))

  }

  if (i=="Reco_gC_m2") {
    ylab = expression(paste("Observed Reco g C m"^{-2}, month^{-1}))
    xlab = expression(paste("Predicted Reco g C m"^{-2}, month^{-1}, " (8 km model)"))

  }



  # # Extract the final model details
  preds <- mod$pred %>%
    data.frame()


  # Merge
  obspred <- merge(modeldata2, preds, by.x="samplerow", by.y="rowIndex")



  # First plot scatterplots for each variable based on individual models
  # Max and min of several columns
  scale_max <- max(c(obspred$obs, obspred$pred))
  scale_min <- min(c(obspred$obs, obspred$pred))


  # Merge pred perf
  # access the results from the best performing model: mod$results[which.min(mod$results[, "RMSE"]), ]
  # but I will calculate these by myself
  R2 <- cor(obspred$obs, obspred$pred)^2
  rmse <- rmse(obspred$obs, obspred$pred)
  mae <- mae(obspred$obs, obspred$pred)
  mbe <- tdStats(obspred$pred,obspred$obs,  functions = "mbe") # Negative values indicate
overestimation
  r_stats <- data.frame(R2=R2, RMSE=rmse, MAE=mae, MBE=mbe)


  # Scatterplot with observed and predicted, colored by biome
  p1 <- ggplot(obspred, aes(x=pred, y=obs, colour=factor(Biome))) +
    geom_point(shape = 16,  size=3) + geom_abline(slope = 1) +

    annotate(label = sprintf("\n MAE = %.1f  \n MBE = %.1f    \n R2 = %.2f   \n RMSE = %.1f   \n ",
                         r_stats  %>% dplyr::select(MAE) %>% as.character() %>% as.numeric(),
                         r_stats  %>% dplyr::select(MBE) %>% as.character() %>% as.numeric(),
                         r_stats  %>% dplyr::select(R2) %>% as.numeric(),
                         r_stats  %>% dplyr::select(RMSE) %>% as.numeric()), geom = "text", x = -Inf, y = Inf,
size = 8, hjust = 0, vjust = 1) +
```

```
  theme_pub  + theme(legend.title=element_blank()) + scale_colour_viridis(discrete=TRUE) + xlab(xlab)
+ ylab(ylab)+
  xlim(scale_min, scale_max) + ylim(scale_min, scale_max)

 # Print out
 setwd("/home/master/abcfluxv1_modeling_bigfiles/figures/")
 print(p1)
 dev.copy(png, paste(i, "rf_1km_train_loocv_predperf_biome.png", sep="_"), width=500, height=400)

 dev.off()




 ### Scatterplot with bins
 p11 <- ggplot(obspred) + geom_bin2d(aes(x=pred, y=obs),bins=50) + geom_abline(slope = 1) +
   xlim(scale_min, scale_max) + ylim(scale_min, scale_max)  + theme_pub +
   scale_fill_viridis(direction=-1,  trans = 'log', labels = scales::number_format(accuracy = 1)) +
   annotate(label = sprintf("\n MAE = %.1f  \n MBE = %.1f    \n R2 = %.2f   \n RMSE = %.1f   \n ",
                        r_stats  %>% dplyr::select(MAE) %>% as.character() %>% as.numeric(),
                        r_stats  %>% dplyr::select(MBE) %>% as.character() %>% as.numeric(),
                        r_stats  %>% dplyr::select(R2) %>% as.numeric(),
                        r_stats  %>% dplyr::select(RMSE) %>% as.numeric()),
            geom = "text", x = -Inf, y = Inf, size = 8, hjust = 0, vjust = 1) +
   labs(fill="Log counts") + xlab(xlab) + ylab(ylab)

 print(p11)
 dev.copy(png, paste(i, "1km_train_loocv_predperf_bins.png", sep="_"), width=550, height=400)
 dev.off()


 ### Model fit
 library("randomForest")
 predtest <- predict(mod, obspred)
 obspred$predtest <- predtest


 p11 <- ggplot(obspred) + geom_bin2d(aes(x=predtest, y=obs),bins=50) + geom_abline(slope = 1) +
   xlim(scale_min, scale_max) + ylim(scale_min, scale_max)  + theme_pub +
   scale_fill_viridis(direction=-1,  trans = 'log', labels = scales::number_format(accuracy = 1))+
   labs(fill="Log counts") + xlab(xlab) + ylab(ylab)

 print(p11)
 dev.copy(png, paste(i, "1km_train_loocv_modelfit_bins.png", sep="_"), width=500, height=400)
 dev.off()


 ### VARIABLE IMPORTANCE ###



 all_varImp <- vi_permute(mod$finalModel, method="permute", metric="rmse",
                       pred_wrapper=predict, nsim=100, train=modeldata2, target=i, keep = TRUE)
 all_varImp <- subset(all_varImp, Importance!=0) #
```

```
  all_varImp$Variable <- ifelse(all_varImp$Variable=="Interval", "Month", all_varImp$Variable)
  importance_error <- attributes(all_varImp)$raw_scores %>% data.frame()
  importance_error <- subset(importance_error, permutation_1!=0) #
  importance_error$Variable <- all_varImp$Variable
  importance_error_wide <- pivot_longer(importance_error, cols=1:100, names_to = "nsim",
values_to="Importance")

  all_varImp <- importance_error_wide

  # plot
  p1 <- ggplot(all_varImp) + geom_boxplot(aes(x=Variable, y=Importance))   +
    coord_flip() + scale_x_discrete(limits = rev(levels(all_varImp$Variable)))+
    theme_pub2
  # Print out
  setwd("/home/master/abcfluxv1_modeling_bigfiles/figures/")
  print(p1)

  dev.copy(png, paste( i, "1km_rf_vip.png", sep="_"), width=1300, height=1100)
  dev.off()


  ### PARTIAL DEPENDENCE PLOTS ###

  nvars <- length(Baseline_vars_1km)


  library("randomForest")

  for (nvar in 1:nvars) {

    pd1 <- pdp::partial(mod, pred.var = Baseline_vars_1km[nvar], train=obspred)  # don't set plot = TRUE

    rug_data <- obspred[Baseline_vars_1km[nvar]]


    if (!is.factor(obspred[, Baseline_vars_1km[nvar]])) {

      pdp_plot1 <- ggplot(pd1, aes(x=pd1[, 1], y=pd1[, 2])) +
        geom_line(size=2) + theme_pub  +labs(y="yhat", x=Baseline_vars_1km_name[nvar]) +
        theme(legend.position = "none") +
        geom_rug(data=rug_data, aes(x=rug_data[, 1]), inherit.aes = FALSE) + ylim(c(ymin, ymax))
    } else if  (is.factor(obspred[, Baseline_vars_1km[nvar]])) {

      pdp_plot1 <- ggplot(pd1, aes(x=pd1[, 1], y=pd1[, 2])) +
        geom_point(size=4) + theme_pub  +labs(y="yhat", x=Baseline_vars_1km_name[nvar]) +
        theme(legend.position = "none") # + ggtitle(title)
    }


    # write out
    output_name <- Baseline_vars_1km[nvar]


    dev.copy(png, output_name, width=600, height=400)
    print(pdp_plot1)
    dev.off()
```

```
  } # nvars loop

} # resp vars loop


### MODEL UNCERTAINTY ###

## Bootstrapping the data

# Number of bootstrap samples
reps <- 20

# Take 20 stratified samples from the data
for (flux in resp_vars) {


  # dataset
  modeldata3 <- d[,c("Study_ID_Short", "id", flux, Baseline_vars_1km)]
  modeldata2 <- na.omit(modeldata3)
  sapply(modeldata2, function(x) sum(is.na(x))) # no missing data


  size <- nrow(modeldata2)
  mat <- data.frame(matrix(nrow=size, ncol=reps))
  modeldata2$rows <- seq(1, size, by=1)

  for (i in 1:reps) {
    print(i)
    sample_fraction_df<- modeldata2 %>% slice_sample(n = size, replace = TRUE) # has to be
replace=TRUE because otherwise the datasets will be identical because we are not changing the size of
the training data
    # Extract only the row numbers for the matrix
    mat[, i] <- sample_fraction_df$rows
  }

  write.csv(mat, paste0("/home/master/abcfluxv1_modeling_bigfiles/results/", flux,
"_bootstrapped_row_numbers_1km.csv"), row.names=FALSE)


}


### after this the model was trained with all these 20 different datasets,
# and fluxes predicted to the entire domain using the 20 different model versions


### ZONAL STATISTICS EXAMPLES ###
```

```
# download a prediction raster
r <- rast(f)


# biomes
biome <- vect("/home/master/masking_summary_rasters/Ecoregions2017_tundraboreal.shp")
biome <-  aggregate(biome, by='BIOME_NAME')
crs(biome) <- "EPSG:4326"


###  zonal statistics: mean

# whole region
abzmean <- global(r, mean, na.rm=TRUE) %>% data.frame() #have to add na.rm=TRUE, otherwise will
get na


#biomes
biomemean <- zonal(r, biome, mean, na.rm=TRUE)
biomemean$class <- biome$BIOME_NAME


### zonal statistics: budget

# calculate pixelwise budget and transform to Tg
r <- r*(res(r)*res(r)) *1.E-12

# whole region
abzsum <- global(r, sum, na.rm=TRUE) %>% data.frame() #have to add na.rm=TRUE, otherwise will get
na


#biomes
biomesum <- zonal(r, biome, sum, na.rm=TRUE)
biomesum$class <- biome$BIOME_NAME


### TREND ANALYSIS EXAMPLES ###

# NEE 1 km
trend <- zyp.trend.vector(d$NEE_1km, method="zhang",
                    conf.intervals=TRUE, preserve.range.for.sig.test=TRUE)


### MULTIVARIATE ENVIRONMENTAL DISSIMILARITY SURFACE ###

# environmental predictor rasters
predrasters <- stack(predictor_files)
names(predrasters) <- c("lst", "ndvi", "snow", "soiltemp", "srad", "ph", "soc", "perma")


# model training data (i.e., site-level predictor data)
```

```
d <-
read.csv("/home/master/flux_upscaling_data/results/final/modeldata_avg_allsites_1km.csv")
d <- subset(d, select=c("lst", "ndvi", "snow", "soiltemp", "srad", "ph", "soc", "perma"))

# MESS
dismo_output <- mess(predrasters, d)
```
